# An Integrated Spatially Resolved Mechanistic Model of Hierarchical Auxin–Cytokinin–Ethylene Crosstalk Underlying Root Growth Inhibition in Arabidopsis

**DOI:** 10.64898/2026.08.14.744203

**Authors:** M. Fenech, J.P. Fernandez-Moreno, G.A. Daubermann, A. Nawar, J.S. Taylor, H. Davis, S. Belcapo, A. Budnick, A.E. Yaschenko, C. Xu, H. Hand, J. Jackson, K. Vollen, K. Muller, M.M. Kater, D.S. Moura, J.T. Ascencio-Ibáñez, J.M. Alonso, A.N. Stepanova

**Affiliations:** Department of Plant and Microbial Biology, College of Agriculture and Life Sciences, North Carolina State University, Raleigh, North Carolina, United States of America; Departamento de Ciências Biológicas, Escola Superior de Agricultura “Luiz de Queiroz”, Universidade de São Paulo, Piracicaba, São Paulo, Brazil; Dipartimento di Bioscienze, Facoltà di Scienze e Tecnologie, Università degli Studi di Milano, Milan, Italy; Institute of Experimental Botany of the Czech Academy of Sciences, Rozvojová 263, 165 00 Prague 6, Czech Republic; Department of Molecular and Structural Biochemistry, College of Agriculture and Life Sciences, North Carolina State University, Raleigh, North Carolina, United States of America

**Author notes:** Corresponding authors: Correspondence to Jose M. Alonso or Anna N. Stepanova. Departamento de Engenharia Química e de Alimentos da Universidade Federal de Santa Catarina, Florianópolis, SC 88037-010, Brazil. Departamento Bioquímica y Biología Molecular, Edificio Severo Ochoa-C6, Campus de Rabanales, Universidad de Córdoba, 14071 Córdoba, Spain.

## Abstract

Decoding how plants integrate multiple hormone signals to coordinate growth requires tools capable of resolving pathway interactions at cellular resolution in living tissue. Here we present *ACE* (Auxin–Cytokinin–Ethylene) and *ACE2*, proof-of-concept single-locus reporters to simultaneously capture activity of multiple hormones. Deploying *ACE* alongside well-established reporters, exogenous hormone treatments, and reverse-genetic perturbations of hormone biosynthesis, signaling, and transport in three-day-old etiolated Arabidopsis seedlings, we dissect the spatiotemporal hierarchy governing primary root elongation and root apical meristem (RAM) size. We demonstrate that both ethylene- and cytokinin-triggered root growth inhibition involve a boost of TRYPTOPHAN AMINOTRANSFERASE OF ARABIDOPSIS1 (TAA1)-mediated auxin biosynthesis and AUXIN RESISTANT1 (AUX1)-dependent auxin redistribution. Two spatially distinct auxin responses underlie the respective root growth effects: ethylene expands TAA1-dependent auxin biosynthesis from the root vasculature into the epidermis and promotes AUX1-mediated auxin import into the transition and elongation zones to inhibit cell elongation, while cytokinin confines ethylene-dependent TAA1-boosted activity to the vasculature and drives auxin accumulation in lateral root cap cells to reduce RAM size. Together, these data establish a reciprocal regulatory loop between these hormones, positioning ethylene as a convergence node in auxin–cytokinin crosstalk, and cytokinin as a modulator of the ethylene–auxin interaction. Critically, the changes in cross-activated reporter patterns described for different genetic backgrounds, alongside quantitative assessment of hormone-specific inhibition of the mutants’ growth, were consistent with the multi-hormone network established over two decades of research, and added cell-type-resolved spatial detail and a proposed hierarchy for the etiolated seedling root. Finally, a second-generation reporter, *ACE2*, overcomes key technical limitations of *ACE*, expanding the platform’s capacity toward a higher-order multi-hormone monitoring system. These resources expand the Arabidopsis genetic toolkit and provide a generalizable framework instrumental for dissecting multi-hormone signaling hierarchies at the cellular level.

## Main

Plants continuously integrate signals from multiple phytohormone pathways to coordinate growth, organ patterning, and environmental responses. Among the most consequential of these interactions is the crosstalk among auxin, cytokinin, and ethylene—three hormones whose individual signaling cascades (the nuclear SCF^TIR1/AFB^–Aux/IAA–ARF module^1^, the AHK–AHP–ARR phosphorelay^2,3^, and the EIN2/EIN3 de-repression cascade^4^) are well-characterized, yet whose combined outputs remain difficult to predict from single-pathway analyses (Figure 1A-C). Decades of genetic and biochemical work have established numerous bilateral interactions (Figure 1D): ethylene promotes auxin biosynthesis^5–11^ and transport^11–14^ and bolsters cytokinin signaling through EIN3-mediated repression of type-A *ARRs*^15^; cytokinin induces ethylene^16–18^ and auxin^19–21^ biosynthesis, and modulates polar auxin transport^19,22–29;^ auxin enhances ethylene biosynthesis^30–32^ and signaling^33^ contributing to cells sensitization to ethylene^6^; auxin interaction with cytokinin is complex and context-dependent^34^, and therefore, it can inhibit^21,35^ or induce^36,37^ cytokinin biosynthesis and signaling^38–40^ to fine-regulate the development of root apical meristem^26,28,29,41–43^ (RAM; Figure 1E) and organ initiation^25,34,40^. Given the dense interconnection between these pathways, understanding how a given stimulus—or mutation—propagates through the system demands simultaneous, spatially resolved readout of all three activities with single-cell resolution.

**Figure 1.**
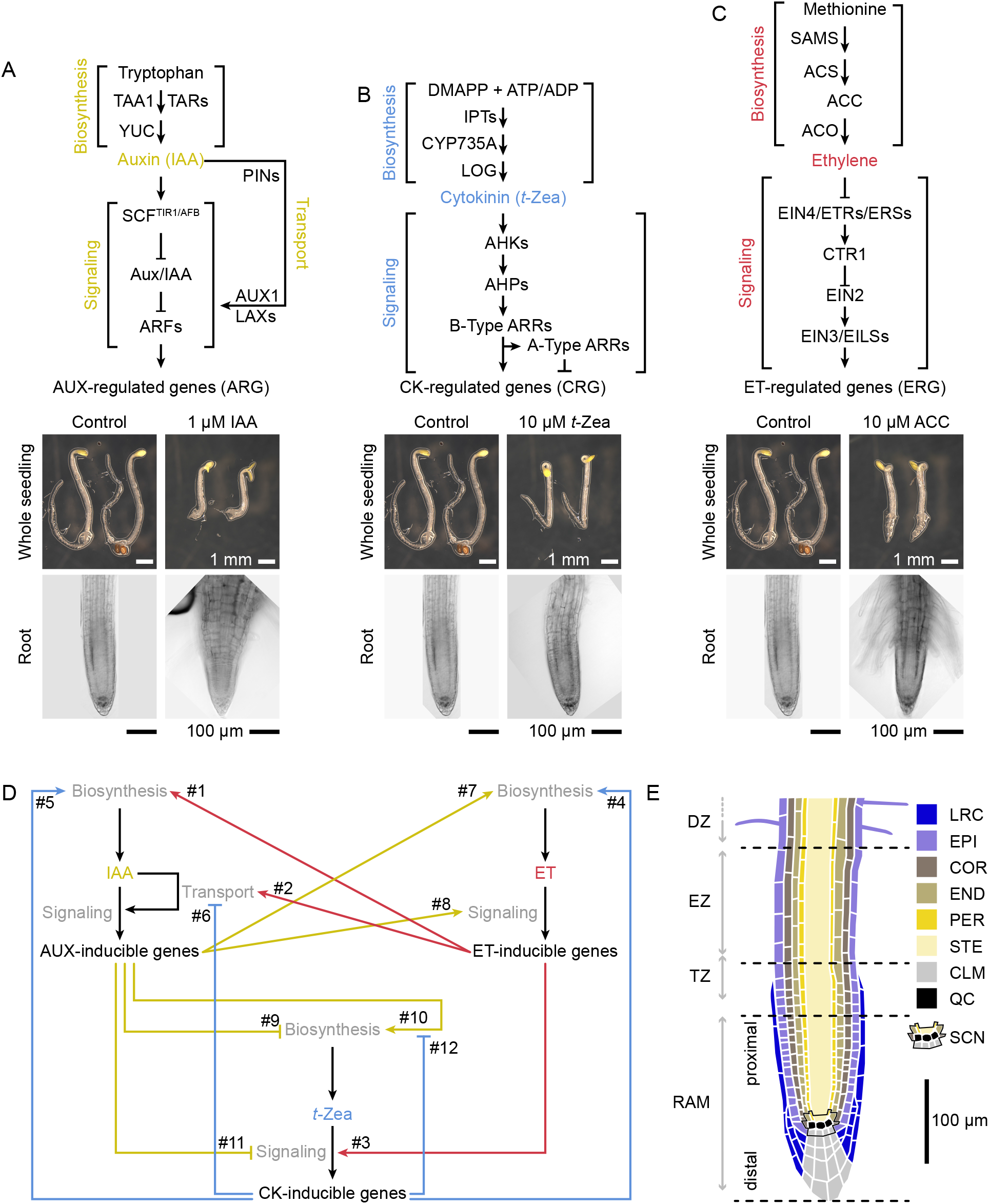
The intricate network of interactions between auxin, ethylene, and cytokinin pathways. (A) Schematic representation of biosynthesis and signaling pathways of auxin indole-3-acetic acid (IAA), cytokinin *trans*-zeatin (*t*-Zea) (B), and ethylene (ET) (C), and the phenotypes of three-day-old seedlings exposed to these growth regulators. Arabidopsis seedlings were germinated on control or hormone-supplemented growth media in the dark. Same control images are presented alongside hormone treatments for better comparison. (A) Previously reported interactions between auxin, ethylene, and cytokinin signaling pathways. Colors represent the hormone that triggers the signaling: red for ethylene, blue for cytokinin, yellow for auxin. Numbers tagged with “#” next to an arrowhead in the figure map indicate the literature references supporting given connections: #1:^5–11;^ #2:^11–14;^ #3:^15^; #4:^16–18;^ #5:^19–21;^ #6:^19,22–29;^ #7:^30–32;^ #8:^6,33^; #9:^21,35^; #10:^36,37^; #11:^38–40;^ #12:^36^. (B) Longitudinal schematic of the Arabidopsis root structure. The root apical meristem (RAM) comprises the cells with active division, and the stem cell niche (SCN) is a microenvironment within the RAM that drives the establishment and maintenance of cell types. The non-meristematic region includes the transition zone (TZ), where cells begin to increase in length; the elongation zone (EZ), where cells undergo rapid elongation; and the differentiation zone (DZ), where root hairs begin to form. The major cell types depicted in the root schematic include lateral root cap (LRC), epidermis (EPI), cortex (COR), endodermis (END), pericycle (PER), vascular tissue (stele, STE), quiescent center (QC), and columella (CLM). The SCN is outlined in black according to the literature^113^. Colors represent different cell types. Scale bars represent the same size within analogous images within a panel. TAA1: TRYPTOPHAN AMINOTRANSFERASE OF ARABIDOPSIS1, TARs: TAA1-RELATED, YUC: YUCCA (FLAVIN-DEPENDENT MONOOXYGENASE), SCFTIR1/AFB: E3 ubiquitin ligase complex [SCF: SKP1/-CUL1/RBX1; TIR1: TRANSPORT INHIBITOR RESPONSE1, AFB: AUXIN SIGNALING F-BOX], AUX/IAA: AUXIN/INDOLE-3-ACETIC ACID INDUCIBLE, ARF: AUXIN RESPONSE FACTOR, PINs: PINOID-LIKE PROTEINs, AUX1: AUXIN RESISTANT1, LAX: LIKE-AUX1, SAMS: S-ADE-NOSYL-L-METHIONINE SYNTHETASE, ACS: 1-AMINOCYCLOPROPANE-1-CARBOXYLATE (ACC) SYNTHASE, ACO: ACC OXIDASE, EIN2/3/4: ETHYLENE INSENSITIVE2/3/4, ETRs: ET RESPONSEs, ERSs: ETHYLENE RESPONSE SENSORs, CTR1: CONSTITUTIVE TRIPLE RESPONSE1, EILs: EIN3-LIKE PROTEINs, DMAPP: Dimethylallyl pyrophosphate, ATP/ADP: Adenosine Triphosphate/Diphosphate, IPTs: ISOPENTENYL TRANSFERASEs, CYP735A: CYTOCHROME P450 MONOOXYGENASE FAMILY 735, SUBFAMILY A, LOG: LONELY GUY, AHKs: ARABIDOPSIS HISTIDINE KINASEs, AHPs: ARABIDOPSIS HISTIDINE PHOSPHOTRANSFER PROTEINs, ARRs: ARABIDOPSIS RESPONSE REGULATORs (B-Type: positive regulators of CK signaling; A-Type: negative regulators).

Our understanding of how cytokinin, auxin, and ethylene work together to control root growth and development has advanced substantially^26,44^, yet the complexity of these regulatory networks continues to reveal new mechanisms of interaction. Phytohormone-responsive synthetic reporters have been the primary tool for mapping hormone activity *in vivo*. Landmark transcriptional reporters including *DR5/DR5rev/DR5v2* (auxin)^45–47^, *TCS/TCSn/TCSv2* (cytokinin)^39,48,49^, and *EBS/EBS-S10/EBSn* (ethylene)^6,50–52^ have each contributed substantially to defining the spatiotemporal distributions of individual hormone signals. However, because each monitors a single pathway, inferring interaction hierarchies has required combining separate transgenic lines—an approach that is genetically laborious, subject to line-to-line variation, and incapable of capturing co-activity patterns within individual cells. Therefore, previous works studying complex developmental processes where hormone interplay has a substantial regulatory role have addressed that limitation by generating single-locus dual reporters^53–57^.

Here we develop *ACE*, a GoldenBraid-compatible, proof-of-concept sensor to monitor <u>a</u>uxin, <u>c</u>ytokinin, and <u>e</u>thylene activity simultaneously in Arabidopsis, uncovering design constraints that are addressed in an improved version, *ACE2*. By integrating this early version of *ACE* reporter with other transcriptional and translational reporters upon targeted genetic perturbations, we successfully map the auxin-ethylene-cytokinin signaling hierarchy governing root elongation in etiolated seedlings. We demonstrate that root growth inhibition triggered by both ethylene and cytokinin involves the spatial regulation of TRYPTOPHAN AMINOTRANSFERASE OF ARABIDOPSIS1 (TAA1)-mediated auxin biosynthesis and requires AUXIN RESISTANT1 (AUX1)-mediated indole-3-acetic acid (IAA) import. This process occurs specifically within the lateral root cap (LRC) and epidermis cells (EPI) of the root transition zone (TZ) and expands into the EPI cells within the elongation and differentiation zones (EZ, DZ, respectively; Figure 1E). Importantly, we demonstrate that regulatory mechanisms underpinning root shortening and RAM size in response to ethylene or to cytokinin are different from one another, providing a comprehensive model for multi-hormone signal integration in Arabidopsis roots at cellular resolution. Finally, we present several variants for *ACE2* reporter, a second-generation single-locus triple-hormone *ACE* reporter that overcomes limitations of the original construct and exhibits improved nuclear targeting and cytokinin-channel output compared with its first-generation counterpart.

## Results

### Construction and validation of the GoldenBraid-compatible library and *ACE* reporter

To enable simultaneous monitoring of three hormone pathways from a single genomic locus, we assembled a GoldenBraid (GB)-compatible library of sequence-diversified synthetic DNA parts, including codon-optimized coding sequences (CDSs) for three spectrally distinct fluorescent proteins (FP) (3xYPet, mTagBFP*, mCherry), and subcellular targeting signals for the nucleus, mitochondria, and peroxisomes (Supplementary Table S1; mTagBFP* corresponds to a hybrid version of mTagBFP that contains mTagBFP2-like amino acid substitution in mTagBFP chromophore moiety but does not include the mTagBFP2 N-terminus). Additionally, we generated a library of Cauliflower mosaic virus (CaMV*) 35S* promoter (*35Sp*)-derived minimal promoters (*minPro*) and terminators (*ter*) as well as the minimal promoters from Figwort mosaic virus (FMV DxS strain subgenomic VI (*19S*)^58,59^) and Horseradish latent virus (HRLV full-length transcript^60^), supplemented with viral-sourced full promoters from Strawberry vein banding virus (SVBV subgenomic transcript VI^61,62^), FMV (FMV M3 strain subgenomic transcript *V*I (*19S*)^63,64^), and Cassava vein mosaic virus (CVMV full-length transcript^63,65^) for constitutive gene expression (Supplementary Figure S1A-E). Systematic sequence diversification of all repeated elements—CDSs, *35SminPro*, *35Ster*, and subcellular targeting signals—was implemented to minimize transcriptional interference and reduce the risk of epigenetic silencing, two key failure modes^66^ in multi-reporter assemblies (Supplementary Figure S1).

From this library, we assembled—and validated by *Nicotiana benthamiana* agroinfiltration—*ACE*, a single-locus triple reporter in which the auxin-responsive *DR5v2*^47^, cytokinin-responsive *TCSn*^48^, and ethylene-responsive *EBSn*^52^ synthetic promoters drive expression of *3xYPet*^67^ (with the respective protein fused to 3xSV40 peptide for nuclear import^51,68^), *mTagBFP*\* ^69^ (fused to MitCOx4^70,71^ for mitochondrial import), and *mCherry*^72,73^ (fused to the KSRM^74^ Peroxisomal Targeting Signal 1 (PTS1)), respectively (Figure 2A; Supplementary Figures S1F and S2). Stable Arabidopsis transformants maintained *ACE* reporter activity through the T5 generation, confirming transgenerational stability without evidence of silencing. Exogenous application of saturating concentrations of individual hormones to three-day-old etiolated seedlings confirmed the expected inducibility: auxin IAA activated the yellow FP 3xYPet, cytokinin *trans*-zeatin (*t*-Zea) activated the blue FP mTagBFP*, and the ethylene precursor ACC activated the red FP mCherry (Figure 2B), each accompanied by expected morphological responses including hypocotyl and root shortening and altered hook angle (Figure 1A-C). Critically, cross-activation was also detected: high IAA and *t*-Zea concentrations induced *EBSn* activity in the root apical meristem (RAM), transition and elongation zones (TZ and EZ, respectively), but more prominently in the differentiation zone (DZ) where no fluorescence was observed under control conditions (Figure 2B); and ACC and *t*-Zea induced spatially distinct *DR5v2* activity, with ACC activating *DR5v2* in the RAM, EZ and DZ, and *t*-Zea inducing it in the proximal RAM and the DZ (Figure 2B). This cross-reactivity, rather than being a limitation, constitutes the primary informational output of *ACE*: combined with hormone-specific mutants, it enables cell-type-resolved dissection of the hierarchy governing hormone signaling interactions in these root tissues.

**Figure 2.**
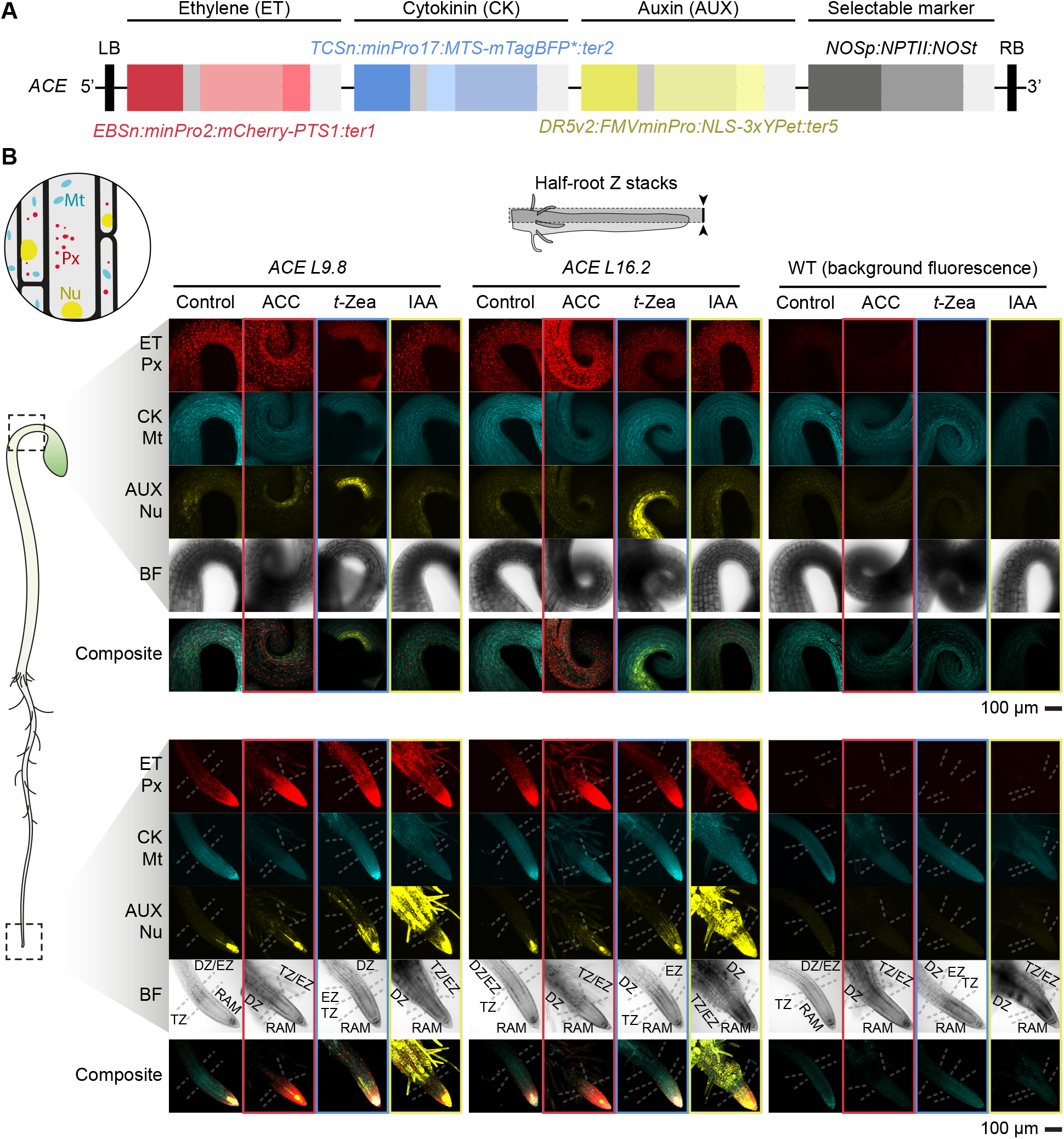
Characterization of the *ACE* triple hormone reporter in stable Arabidopsis transgenic lines. (A) *ACE* construct structure. Fragments not at scale. (B) Reporter activity in three-day-old etiolated seedlings under endogenous concentrations of hormones (AT) and upon exogenous hormone supplementation reveals hormone crosstalk between hormone signaling pathways, which is a desirable feature of this reporter. Circular schematic shows the expected subcellular localization of the fluorescent proteins (Mt: mitochondrial mTagBFP*, Px: peroxisomal mCherry, Nu: nuclear 3xYPet). All microscopy images were acquired in the same experimental session. Confocal microscopy images show stacked pictures from equatorial to cortical planes. *L9.8* and *L16.2* refer to two independent single-insertion lines. Scale bar applies to all images within the panel. Single-channel stacks were contrast-enhanced to facilitate detection of low-expressing cells, whereas composite images (all three channels) were adjusted to enable simultaneous visualization of all reporters. LB: left T-DNA border; RB: right T-DNA border; *PTS1: Peroxisomal Targeting Signal 1* (N-KS-RM-C). *MTS: Mitochondrial Targeting Signal* (MitCox4); *NLS: Nuclear Localization Signal* (3xSV40); ACC = 10 µM 1-aminocyclopropane-1-car-boxylic acid; *t*-Zea = 10 µM *trans*-zeatin; IAA = 1 µM indole-3-acetic acid; ET: ethylene reporter (*EBSn:mCherry-PTS1*); CK: cytokinin reporter (*TCSn:MTS-mTagBFP\**); AUX: auxin reporter (*DR5v2:3xSV40-3xYPet*); mTagBFP*: Hybrid version of mTagBFP that contains mTagBFP2-like amino acid substitution in mTagBFP chromophore moiety but does not include mTagBFP2 N-terminal; BF: bright field.

### *ACE* resolves a self-reinforcing cytokinin–ethylene–auxin loop in the root transition zone

To probe hormone interaction hierarchy systematically, we studied the crosstalk between auxin, cytokinin, and ethylene by leveraging mutants defective in hormone biosynthesis, signaling, and transport. We introgressed *ACE* into cytokinin receptor (*ahk3 cre1*^75^), auxin biosynthesis (*wei8^7^*), auxin overproduction (*sur2*^76^), auxin import (*aux1*^77,78^), and ethylene signaling (*ein2*^79^, *ctr1*^80^) mutants, and monitored all three reporters in response to non-saturated equi-effective individual hormone concentrations that achieve ∼50% root length reduction compared to control in three-day old wild-type (WT) etiolated seedlings. Furthermore, we supplemented this assay by quantifying the growth inhibitory effect of these hormone concentrations on each mutant (Supplementary Figure S3). These measurements provide a more comprehensive assessment of hormone activity than solely considering fluorescence activity, since the growth inhibitory effect of a hormone may not only depend on the signaling pathway that a given mutant disrupts.

Because the *TCSn* transcriptional unit in *ACE* showed lower-than-expected^48^ performance (see *Technical limitations* section below), we supplemented *ACE* analysis with that of a previously validated, upgraded version of the cytokinin reporter, *TCSv2:3xmVenus-N7*^49^, to ensure accurate readout of cytokinin signaling (Figure 3A; Supplementary Figure S4A). In WT seedlings, exogenous *t*-Zea enhanced *TCSv2* reporter activity in the RAM and root vasculature (stele cells; STE) and simultaneously activated *EBSn* in the RAM, TZ, EZ, and DZ of the root, while *DR5v2* was predominantly activated in the LRC (Figure 3A; Supplementary Figure S4A). Reciprocally, IAA and ACC each induced *TCSv2* specifically in the TZ and EZ (Figure 3A; Supplementary Figure S4A) although this cytokinin reporter expression decreased in the distal RAM, underscoring the spatial organization of hormone signaling interactions in the root and highlighting the TZ/EZ region as a spatial crosstalk hub between auxin, cytokinin, and ethylene. In the cytokinin receptor double mutant, *ahk3 cre1*, *TCSv2* activity in the vasculature decreased compared to WT, and its induction by *t*-Zea, IAA, and ACC was abolished, while basal auxin and ethylene reporter activities were maintained (Figure 3A; Supplementary Figure S4A). Notably, the *DR5v2* response to IAA was slightly reduced in the DZ of the mutant, whereas ACC-induced *EBSn* activity was unaffected and *t*-Zea-mediated induction of *EBSn* was milder than in WT (Figure 3A; Supplementary Figure S4A). These results point to auxin and ethylene acting mainly at the level of or upstream of the cytokinin receptors, which is supported by previous evidence where auxin promotes the induction of *IPT* cytokinin biosynthesis genes in Arabidopsis roots^36^. Phenotypically, *ahk3 cre1* root elongation was prominently insensitive to *t*-Zea and mildly insensitive to ACC but responded normally to IAA (Figure 3B; Supplementary Figure S3). These results define a self-reinforcing hormone signaling loop in the root in which cytokinin activates ethylene and auxin, which in turn sustain cytokinin activity through AHK3 and CRE1 receptors (Supplementary Figure S4B). This self-reinforcing loop seems to be required for normal auxin sensitivity in the differentiation zone. Importantly, the WT-like root response of *ahk3 cre1* to IAA and the very mild insensitivity of this mutant to ACC (Supplementary Figure S3) indicate that the AHK3- and CRE1-dependent, IAA- and ethylene-induced boost in cytokinin signaling observed in the TZ of WT roots (Figure 3A; Supplementary Figure S4A) is fully dispensable for auxin- and partially dispensable for ethylene-mediated root growth inhibition. In other words, disruption of this regulatory loop by the cytokinin receptor mutant does not prominently affect the growth inhibitory effects of ethylene and auxin, but it interferes with normal auxin response in the DZ, suggesting that complex tissue-specific interactions between these three hormones may regulate developmental processes beyond cell elongation.

**Figure 3.**
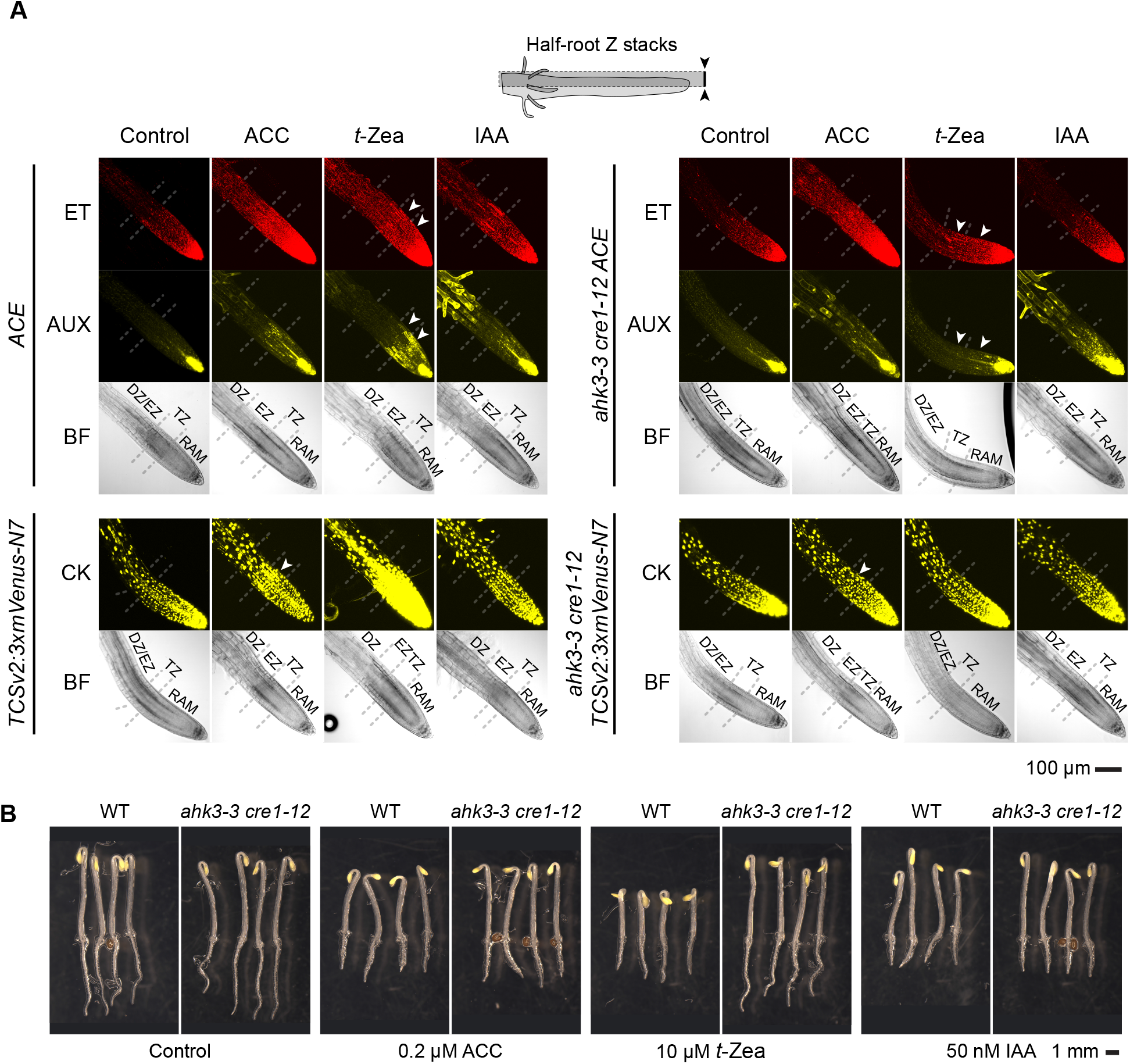
AHK3 and CRE1 are required for *t*-Zea-mediated increase in auxin and ethylene signaling in Arabidopsis. (A) Cytokinin-insensitive *ahk3 cre1* mutant shows normal basal activity and inducibility of ethylene and auxin reporters but suppresses the induction of cytokinin reporter *TCSv2* by exogenous ACC, *t*-Zea, and IAA in roots of three-day-old etiolated seedlings. Arrowheads point at root sections where reporter expression differs between treatments or genotypes. (B) *ahk3 cre1* roots show strong insensitivity to *t*-Zea and a mild, yet significant, insensitivity to ACC. Quantitative comparison and statistical assessment of hormone-specific inhibitory effect on growth between WT and mutant are available in Supplementary Figure S3. All microscopy images were acquired in the same experimental session. Confocal microscopy images show stacked pictures from the equatorial plane at the quiescent center to the cortical plane of the lateral root cap. Scale bar applies to all images within the panel. ACC = 0.2 µM 1-aminocy-clopropane-1-carboxylic acid; *t*-Zea = 10 µM *trans*-zeatin; IAA = 50 nM indole-3-acetic acid; ET: ethylene reporter (EBSn:mCherry-PTS1); CK: cytokinin reporter (*TCSv2:3xmVenus-N7*)^49^; AUX: auxin reporter (*DR5v2:3xSV40-3xYPet*); BF: bright field; DZ: differentiation zone; EZ: elongation zone; TZ: transition zone; RAM: root apical meristem. Root zone boundaries are approximate.

### TAA1-dependent auxin biosynthesis is the primary executor of ethylene- and cytokinin-mediated root growth inhibition

To establish the epistatic relationship between upstream ethylene and cytokinin signals and auxin output, we examined *ACE* activity in *wei8* mutant, whose TAA1-impaired activity leads to auxin biosynthesis deficiency (Figure 1A). Basal *DR5v2* and *EBSn* activities were reduced in *wei8* compared to WT (Figure 4A; Supplementary Figure S5A). Critically, although ACC induced *EBSn* activity normally, neither ACC nor *t*-Zea increased *DR5v2* like WT (Figure 4A; Supplementary Figure S5A). Consistently, *wei8* roots were mildly insensitive to ACC- and *t*-Zea-mediated shortening (Figure 4B; Supplementary Figure S3). These findings suggest that a reduction in baseline auxin biosynthesis desensitizes cells to ethylene, as evidenced by reduced basal *EBSn* reporter activity in the *wei8* mutant. However, when treated with exogenous ACC, the previously reported induction of TAR2 and other downstream auxin biosynthesis genes by ethylene^7,9^ can partially compensate for this defect. Even low concentrations of ACC (0.2 µM) restored downstream *EBSn* reporter induction to WT levels (Figure 4B; Supplementary Figure S3). Nevertheless, this backup mechanism is insufficient to restore normal ethylene-mediated root growth inhibition in *wei8*, which ultimately depends on TAA1-driven auxin biosynthesis as its primary downstream effector (Supplementary Figure S5B). The same logic applies to cytokinin: *t*-Zea induces auxin reporter activity in the LRC in a TAA1-dependent manner (Figure 4A; Supplementary Figure S5A), and the partial *t*-Zea insensitivity of *wei8* roots (Supplementary Figure S3) confirms that cytokinin-triggered root growth inhibition similarly relies on TAA1-mediated IAA production.

**Figure 4.**
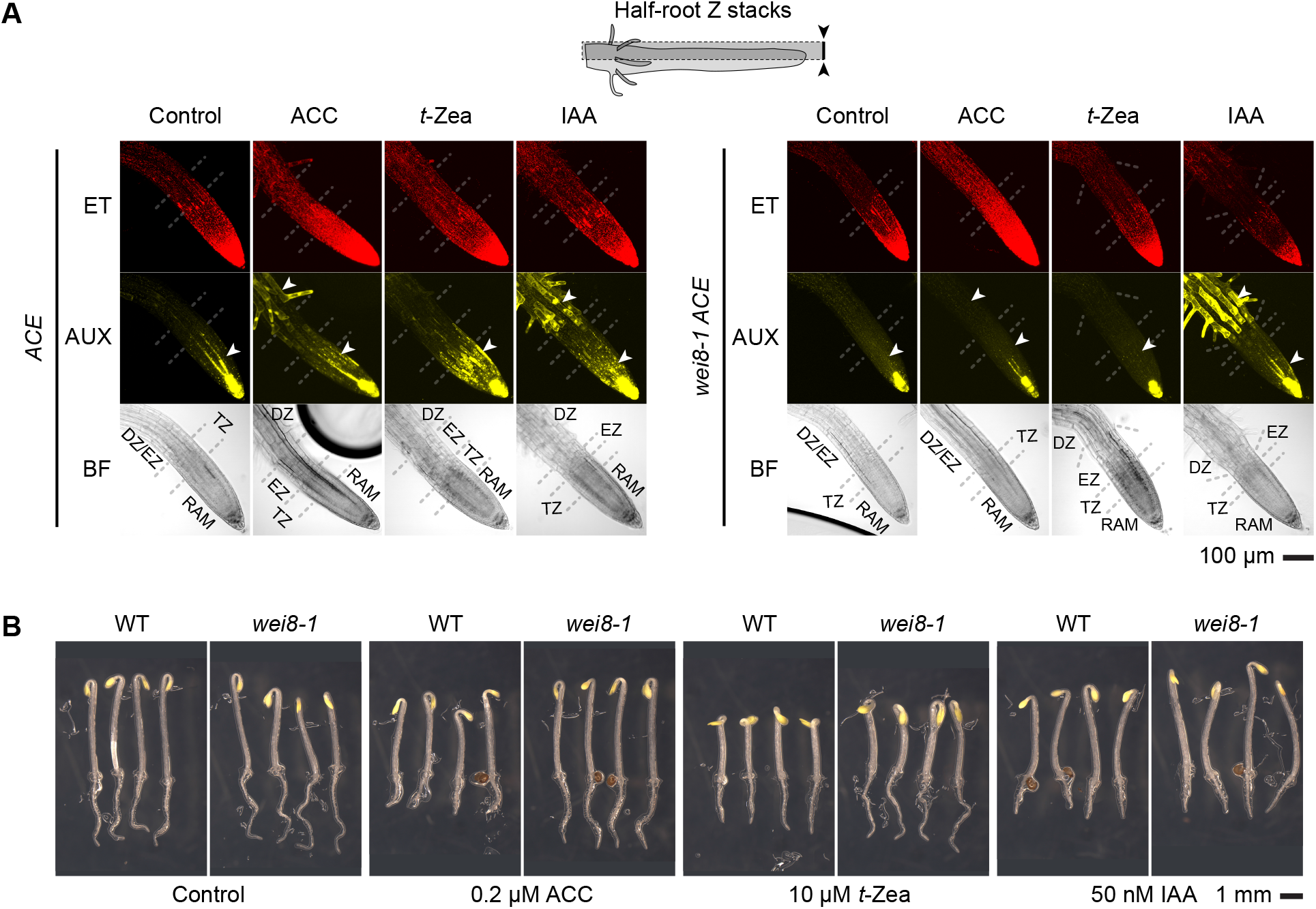
TAA1 is required for ACC- and *t*-Zea-mediated increase in auxin signaling in Arabidopsis. (A) TAA1-deficient *wei8* mutant has weaker basal AUX activity but normal ET signaling in roots of three-day-old etiolated seedlings. Arrowheads point at root sections where reporter expression differs between treatments or genotypes. (B) *wei8* roots show mild, yet significant, insensitivity to ACC and *t*-Zea. Quantitative comparison and statistical assessment of hormone-specific inhibitory effect on growth between WT and mutant are available in Supplementary Figure S3. All microscopy images were acquired in the same experimental session. Confocal microscopy images show stacked pictures from the equatorial plane at the quiescent center to the cortical plane of the lateral root cap. Scale bar applies to all images within the panel. ACC = 0.2 µM 1-aminocy-clopropane-1-carboxylic acid; *t*-Zea = 10 µM *trans*-zeatin; IAA = 50 nM indole-3-acetic acid; ET: ethylene reporter (*EBSn:mCherry-PTS1*); AUX: auxin reporter (*DR5v2:3xSV40-3xYPet*); BF: bright field. Root zone boundaries are approximate.

In the auxin overproducing mutant *sur2*, elevated basal auxin correlated with enhanced *EBSn* and *DR5v2* induction by both ACC and *t*-Zea in the root DZ and RAM, respectively (Supplementary Figure S6A), suggesting that high basal auxin sensitizes roots to ethylene and cytokinin. On the other hand, *sur2* root growth inhibition displayed exaggerated sensitivity to *t*-Zea but not to ACC (Supplementary Figure S3, S6B), suggesting that the interactions between auxin–cytokinin and auxin–ethylene signaling are highly dependent on developmental context and hormone concentration.

To determine whether a cytokinin-induced boost in auxin biosynthesis requires intact ethylene signaling, we examined *ACE* reporter activity in the ethylene-insensitive mutant *ein2* (Figure 5). As expected^52^, in *ein2*, ACC failed to induce *EBSn* and, therefore, *DR5v2* activity (Figure 5A, C). Remarkably, *t*-Zea also failed to boost *DR5v2* activity in *ein2* (Figure 5A, C), indicating that EIN2-dependent ethylene signaling is required, under these conditions, for the cytokinin-to-auxin biosynthesis relay (Supplementary Figure S7) and recapitulating the *DR5v2* response to *t*-Zea of *wei8* mutant. Consistent with these results, analysis of the TAA1 translational reporter in the *wei8* background (*wei8 TAA1p:YPet-gTAA1*^8,81^) showed that *t*-Zea-induced TAA1 accumulation partially overlapped with that of the ACC-induced pattern—specifically in the root vasculature, but not in the LRC or EPI cells, where ACC-induced YPet-TAA1 maxima are observed (Supplementary Figure S8).

**Figure 5.**
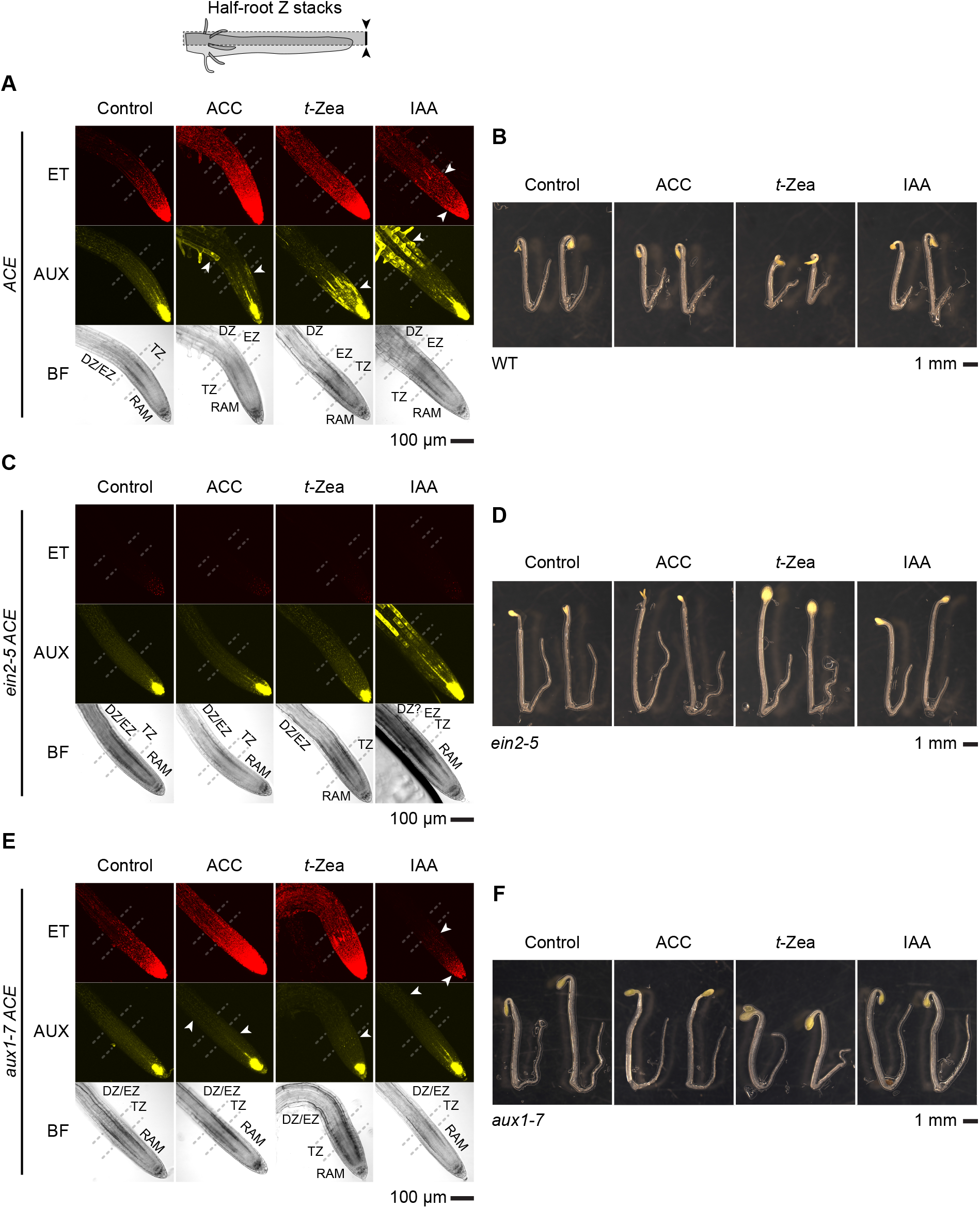
Cytokinin-mediated switch of auxin maxima in Arabidopsis requires functional ethylene signaling and depends on auxin import for full output. (A) *ACE* reporter activity in WT background. (B) Three-day-old WT etiolated seedlings grown in unsupplemented and hormone-supplemented media. (C) Auxin signaling activity in the ethylene-insensitive mutant *ein2* is comparable to that of WT (despite fewer root hairs in *ein2*, the same cells responded to exogenous IAA in both genotypes), but ethylene signaling is completely abolished in *ein2*. Moreover, EIN2 is required for cytokinin-and ACC-mediated increase in auxin signaling. (D) *ein2* roots are insensitive to exogenous ACC, show mild, yet significant, insensitivity to exogenous IAA (similar to that of the mild auxin-insensitive *tir1* mutant, supporting previous findings that ethylene sensitizes the root to auxin^6^; Supplementary Figure S3) and to *t*-Zea (similar to that of *wei8*, suggesting that TAA1 and EIN2 are necessary for *t*-Zea-mediated root growth inhibition; Supplementary Figure S3). The fact that *ein2* suppresses cytokinin-induced auxin biosynthesis, but its roots are not completely insensitive to *t*-Zea, suggests that only part of cytokinin-mediated decrease in root length relies on EIN2-dependent increase in auxin biosynthesis, compatible with an active ethylene-dependent EIN2-independent signaling pathway^114,115^. (E) The import-deficient *aux1* mutant has weaker auxin activity and shows severely impaired IAA-, ACC-, and *t*-Zea-mediated increase in AUX activity, and weaker EBSn response to ACC in the EZ/DZ, leading to unchanged root architecture in response to exogenously applied hormones in three-day-old etiolated seedlings. Arrowheads point at root sections where reporter expression differs between treatments or genotypes. (F) *aux1* roots show strong insensitivity to ACC and IAA and mild, yet significant, insensitivity to *t*-Zea, similar to that of *wei8*. Quantitative comparison and statistical assessment of hormone-specific inhibitory effect on growth between WT and mutants are available in Supplementary Figure S3. All microscopy images were acquired in the same experimental session. Confocal microscopy images show stacked pictures from the equatorial plane at the quiescent center to the cortical plane of the lateral root cap. Scale bar applies to all images within the panel. ACC = 0.2 µM 1-aminocyclopropane-1-carboxylic acid; *t*-Zea = 10 µM *trans*-zeatin; IAA = 50 nM indole-3-acetic acid; ET: ethylene reporter (*EBSn:mCherry-PTS1*); AUX: auxin reporter (*DR5v2:3xSV40-3xYPet*); BF: bright field. Root zone boundaries are approximate.

Finally, to further investigate possible synergistic interactions between ethylene and these other two hormones, we next examined the effects of the hormones on the ACE activity in the constitutive ethylene signaling mutant, *ctr1*. The *ctr1* mutant displayed elevated basal *EBSn* and *DR5v2* activity under control conditions, with a partially saturated hormonal response phenotype—consistent with the expected activation of the ethylene-mediated auxin response in this mutant (Supplementary Figures S3, S9). Noteworthy, high basal ethylene signaling activity in *ctr1* roots was further enhanced in the vascular tissue of DZ upon exogenous *t*-Zea treatment, concomitant with a remarkable increase of auxin activity in the LRC of the RAM and TZ along with the EPI cells of the DZ. However, exogenous ACC and IAA treatments increased auxin activity more prominently in the EPI cells of *ctr1* non-meristematic tissue (TZ/EZ, DZ) than in the LRC (Supplementary Figure S9). On the other hand, we examined the effects of exogenous IAA on *ACE* activity in the ethylene insensitive mutant *ein2*. IAA application induced root thickening and root hair formation in WT but not in *ein2*, with *ein2* roots showing mild, yet significant, insensitivity to IAA (Figure 5B, D). This weak resistance was comparable to that of the auxin perception-deficient mutant *tir1*^82^, supporting previous findings that ethylene sensitizes the root to auxin^6^ (Supplementary Figure S3) and indicating that some auxin-mediated morphological responses also involve downstream activation of ethylene signaling.

Together, these results suggest that while the root TZ maintains a self-reinforcing hormone signaling loop (Supplementary Figure S7B), the three signals interact in a hierarchical fashion within the RAM and EZ. In these tissues, cytokinin acts upstream of ethylene to regulate TAA1-mediated auxin biosynthesis and subsequent root growth inhibition. However, we considered that a basic linear signaling model would probably be an oversimplification for the following reasons: (1) our results show two distinct patterns of auxin activity upon exogenous hormone treatments, with the *t*-Zea-induced pattern occupying a higher hierarchical position than the patterns induced by ACC, IAA, or their corresponding constitutively active genetic backgrounds (Supplementary Figures S6 [*sur2*] and S9 [*ctr1*]); (2) the spatial patterns of ethylene- and cytokinin-induced TAA1 accumulation only partially overlap; (3) despite ethylene insensitivity abolishing *t*-Zea-increased auxin activity in *ein2* roots, its resistance to *t*-Zea-triggered growth inhibition is incomplete; (4) it is consistently reported that cytokinin inhibits PIN-mediated IAA efflux^24^ and decreases AUX1 accumulation in the EPI cells within the EZ, but not in the LRC^26^—consistent with *t*-Zea inducing auxin accumulation in the LRC (Figures S4A, S5A, S6B, S7A)—and that (5) auxin maxima is established through the combined action of local auxin biosynthesis^8,9^, its transport *via* auxin carriers^8,83^ and plasmodesmata^84^, and auxin degradation^85,86^. It was previously reported that cytokinin inhibits root growth through ethylene-dependent and -independent mechanisms^26^, which is consistent with (a) *t*-Zea-induced ethylene-mediated auxin biosynthesis (Figure 5A-D; Supplementary Figure S7A,B) and (b) altered auxin distribution due to reportedly decreased auxin carrier accumulation^19,22–29^. Hence, *t*-Zea-treated *ein2* roots are insensitive to (a), but limited distribution of endogenous auxin still impairs *ein2* root growth (b).

### AUX1-dependent auxin influx mediates cytokinin-induced auxin redistribution from vascular source to root tip sink tissues

The spatial discordance between *t*-Zea-induced *TAA1* accumulation in the vasculature (source cells) and the corresponding *DR5v2* maxima in the LRC (sink cells) may suggest that auxin transport from source to sink cells underlie the formation of these auxin maxima. According to the inverted fountain model of auxin flux in the primary root, IAA is transported rootward through the vasculature into the stem cell niche (SCN), from which it is redistributed shootward through the LRC and into epidermal (EPI) cells of the elongation and differentiation zones (EZ and DZ)^83,84,87^. In this model, polar PIN transporters determine the direction of auxin efflux, whereas nonpolar AUX1/LAX influx carriers regulate sites of auxin accumulation^83,84^. Cytokinin is known to inhibit auxin distribution along this transport pathway^24,84^. Consistent with this, we observed sustained auxin accumulation in LRC cells alongside low *DR5v2* activity in the vasculature of *t*-Zea-treated plants (Figures 2–5; Supplementary Figures S4A, S5A, and S7A,C). Given the observation that root vasculature cells potentially serve as the auxin source in *t*-Zea-treated seedlings (Supplementary Figure S8) these results suggest minimal intracellular auxin retention and limited auxin import in the vascular cells. In contrast, the strong auxin response in the LRC suggests net import activity in these cells over a limited IAA efflux, resulting in net auxin accumulation in the LRC within the TZ.

Because AUX1 is the major component leading to auxin accumulation, to test our hypothesis, we introduced *ACE* into the auxin influx carrier mutant *aux1* (Figure 5E, F; Supplementary Figure S7C). Under control conditions, *aux1* showed reduced *DR5v2* activity in the columella (CLM) and vasculature of the proximal RAM compared to its WT counterpart, but its *EBSn* activity was indistinguishable from WT (Figure 5A, E; Supplementary Figure S7C). Loss of auxin influx abolished the IAA-, ACC-, and *t*-Zea-induced increases in *DR5v2* activity (Figure 5A, E; Supplementary Figure S7C), translating into strong root insensitivity to ACC and IAA and mild insensitivity to *t*-Zea (Figure 5B, F; Supplementary Figure S3). Furthermore, the *EBSn* response to ACC was weaker in *aux1* than in WT in the EZ/DZ, and its root architecture was indistinguishable from control (AT) conditions (Supplementary Figure S7C), confirming that AUX1 is essential for ACC-triggered auxin redistribution that drives inhibition of root elongation (Supplementary Figure S7D).

Having established that AUX1-dependent transport is required for cytokinin-induced auxin accumulation in outer root tissues, we next investigated the origin of this auxin. Although *t*-Zea restricted *DR5v2* maxima to the LRC in WT roots (Figures 2-5; Supplementary Figures S4A, S5A, S7A,C), TAA1 accumulation in these tissues remained unchanged relative to control conditions (Supplementary Figure S8). Moreover, the *t*-Zea-induced *DR5v2* response was completely abolished in *wei8* mutants lacking functional *TAA1*, despite the presence of the related paralogs *TAR1* and *TAR2*^7^. Together, these observations argue against cytokinin-induced local auxin biosynthesis in LRC and EPI cells. Instead, they support a model in which IAA is transported shootward from the SCN through the LRC, consistent with the transport model^83,84,87^. Auxin then accumulates in the LRC due to their limited auxin efflux^24^, and EPI cells in the TZ/EZ fail to import auxin due to limited uptake activity and theoretically scarce apoplastic IAA^26^.

Collectively, the mutant analyses support a spatially organized, hierarchical model (Figure 6), where *t*-Zea activates cytokinin signaling in roots leading to subsequent induction of ethylene production and EIN2-dependent signaling that in turn stimulates TAA1-mediated auxin biosynthesis (Figure 6A). Importantly, the use of the *ACE* reporter allowed us to identify tissue specific differences between ethylene and cytokinin treated plants in this general hormone cascade that leads to distinct auxin activity patterns in primary roots. Thus, ethylene treatment stimulates TAA1-mediated auxin biosynthesis in the vasculature along the primary root and in the LRC and EPI cells from the TZ to the DZ. On the other hand, exogenously applied *t*-Zea restricts TAA1 accumulation to the vascular cells despite the induction pattern of ethylene reporter being comparable between *t*-Zea and ACC treatments, suggesting that prior induction of cytokinin signaling (Figure 3; Supplementary Figure S4) promotes the repression of TAA1 accumulation in the outer cell layer of the root. In the absence of exogenous cytokinin, the resulting auxin is then redistributed from the source *TAA1*-expressing cells into the SCN, from where AUX1 mediates auxin reflux shootwards *via* LRC cells as stipulated by the inverted fountain model^87^. Remarkably, we show that *t*-Zea-triggered auxin patterning is hierarchically higher than the patterns established by ACC or IAA (Figure 6B); in the presence of exogenous cytokinin, constitutive activation of auxin (Supplementary Figure S6A,B) or ethylene (Supplementary Figure S9A,B) does not recapitulate exogenous application of IAA or ACC, respectively, but enhances auxin activity patterns triggered by *t*-Zea. Some authors have linked cytokinin treatment of roots with impaired IAA efflux^19,22–25,28^. This repressive role of cytokinin on the auxin transport pathway is consistent with our observation of *t*-Zea-mediated restriction of TAA1 accumulation to the vasculature and with reduced auxin efflux from the LRC, leading to IAA accumulation in this tissue. Rather than a cell-autonomous effect of auxin within LRC cells, this accumulation likely reflects impaired shootward auxin flux—a consequence of cytokinin-mediated downregulation of polar auxin transport—with the resulting reduction in net auxin supply to the meristematic epidermis and TZ contributing to RAM size reduction, where auxin normally sustains cell proliferation. An additional, ethylene-independent contribution of cytokinin to RAM size reduction through direct promotion of TZ cell differentiation cannot be excluded, and would account for the partial *t*-Zea insensitivity of *ein2* roots. Ethylene-triggered root growth inhibition bypasses the cytokinin step and directly stimulates TAA1 in LRC and EPI cells, explaining the partial but not complete overlap between ethylene and cytokinin phenotypes across these mutant backgrounds (Supplementary Figure S3). Finally, the partial resistance of *ein2* roots to *t*-Zea (Supplementary Figure S3) and the IAA-mediated AHK3/CRE1-dependent induction of cytokinin signaling being dispensable for the IAA-triggered root inhibition (Figure 3A; *ahk3 cre1* in Supplementary Figure S3; Supplementary Figure S4A) are consistent with cytokinin regulating root development independently from ethylene but not from auxin, most likely, by altering auxin distribution^24,26,28^ (Figure 6C).

**Figure 6.**
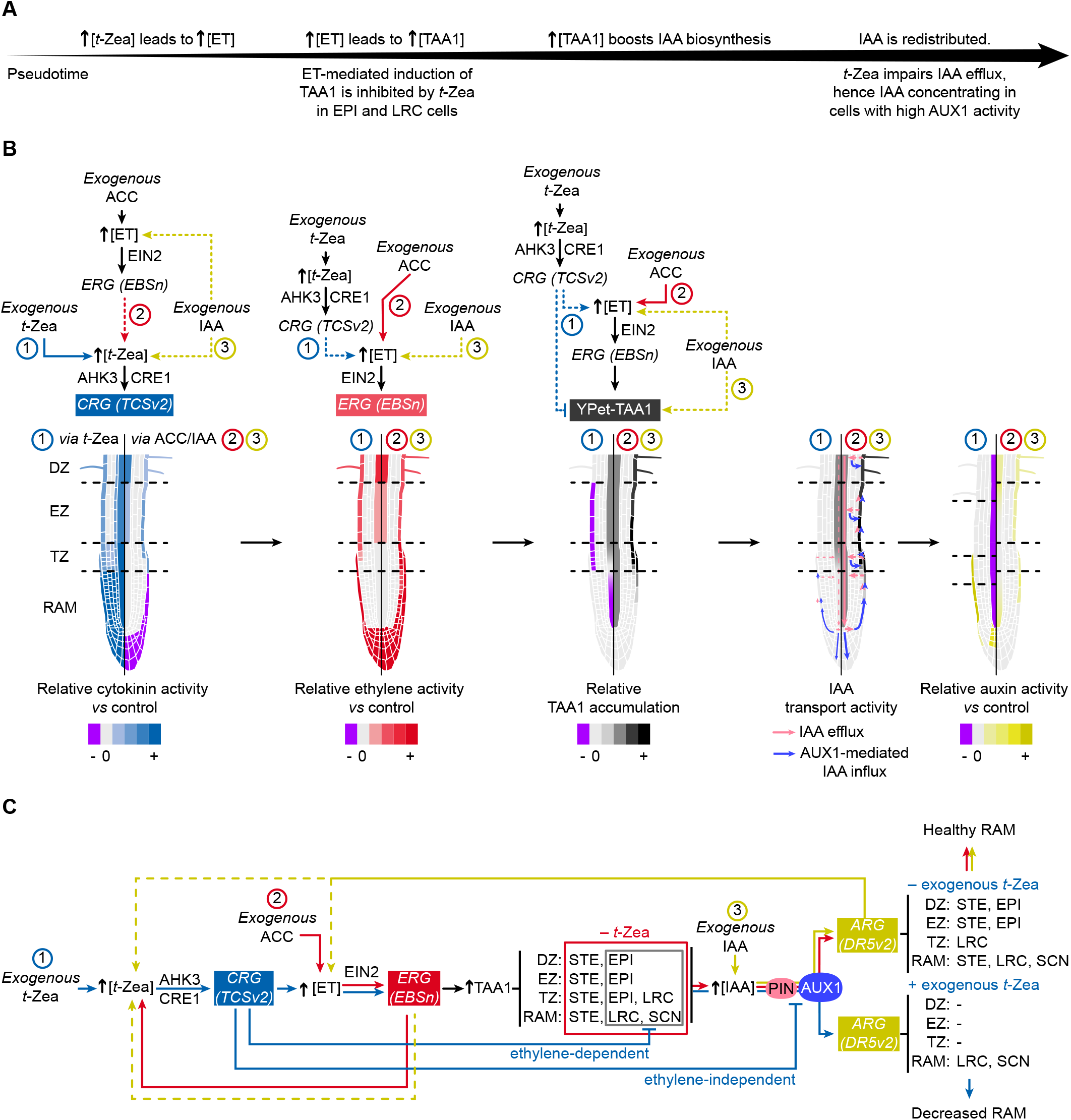
Integrated model of hormone pathway interactions among auxin, cytokinin, and ethylene in the roots of three-day-old etiolated Arabidopsis seedlings. (A) Spatially resolved hormone crosstalk in Arabidopsis roots upon exogenous *t*-Zea treatment, leading to decreased RAM size. Color represents relative change in hormone activity or TAA1 accumulation compared to unsupplemented controls: purple indicates decreased expression/accumulation, light grey (0) indicates no change, and a graded scale from “0” to “+” indicates increasing levels of expression/accumulation. IAA transport activity patterns were inferred from the results reported here (Figure 5E,F; Supplementary Figure S7C,D) and supplemented with previous _studies_24,26,28,29. (B) Network model of hormone signaling activity, including feedback loops, triggered by exogenously applied *t*-Zea (1), ACC (2), or IAA (3) (numbered as in the figure). Unlike panel A, root zones and cell types indicate detected activity rather than relative change compared to control. TAA1-mediated auxin biosynthesis and AUX1-mediated auxin distribution are central to RAM size regulation in response to ethylene and cytokinin. Together, they integrate these signals into a coordinated response by differentially patterning auxin activity. Signaling pathways are color-coded as follows: blue arrows mark *t*-Zea-mediated connections, red arrows signify ACC-mediated connections, and yellow arrows correspond to IAA-mediated connections. Dashed lines represent connections that remained unresolved in this work. Hormones and signaling components: *t*-Zea: trans-zeatin; ACC: 1-aminocyclopropane-1-carboxylic acid; ET: ethylene; IAA: indole-3-acetic acid; EIN2: ETHYLENE INSENSITIVE2; AHK3, CRE1/AHK4: ARABIDOPSIS HISTIDINE KINASEs; TAA1: TRYPTOPHAN AMINOTRANSFERASE OF ARABIDOPSIS1; AUX1: AUXIN RESISTANT1. Reporters: *CRG (TCSv2)*: cytokinin-regulated genes (represented by *TCSv2* reporter); *ERG (EBSn)*: ethylene-regulated genes (represented by *EBSn* reporter); *ARG (DR5v2)*: auxin-regulated genes (represented by *DR5v2* reporter). Anatomical zones: RAM: root apical meristem; SCN: stem cell niche (quiescent center and immediately adjacent cells); TZ: transition zone; EZ: elongation zone; DZ: differentiation zone. Cell types: STE: stele cells (vasculature); EPI: epidermal cells; LRC: lateral root cap cells.

### Technical limitations of *ACE* and design of the improved *ACE2* reporter

The characterization of *ACE* revealed three technical limitations that informed second-generation reporter design. First, the *TCSn:35SminPro17:mTagBFP*:35Ster2* transcriptional unit underperformed: while the activity of individual DNA parts was confirmed in transient expression assays in *Nicotiana benthamiana* (Supplementary Figure S1A-F), this specific *minPro/ter* combination appears to be incompatible within the assembled construct context, despite the early characterization of *minPro* and *ter* parts not revealing this behavior (Supplementary Figure S1B,D). Second, the 3xSV40 nuclear localization signal (NLS) was inefficient in stable Arabidopsis transformants (Supplementary Figure S2), limiting spatial FP segregation. Third, high autofluorescence in the mTagBFP2 emission window partially masked cytokinin reporter signal in root cells (Supplementary Figure S2A), while peroxisomal reporter detection was further hampered by chlorophyll autofluorescence in photosynthetic tissues (Supplementary Figures S10). Broadband filter-based microscopy proved inadequate for spectral separation (Supplementary Figures S11, S12, S13), with single-wavelength laser excitation being the preferred means for reliable multi-channel imaging with this system.

A potential concern in targeting *ACE* reporters to specific organelles is that organelle distribution or abundance might be altered by hormonal shifts. To test this, we monitored peroxisomal dynamics in control and hormone-treated seedlings. Using both *EBSn:mCherry-PTS1* and the established [*35Sp:NeonGreen-PTS^PEX^*^26^] - [*UBQ10p:mRuby3-PTS1*] dual peroxisomal markers^88^, we observed an apparent increase in *UBQ10p:mRuby3-PTS1* activity—but not in *35Sp:NeonGreen-PTS^PEX^*^26^—of roots exposed to IAA (Supplementary Figure S13A-C). Notably, the changes in root architecture induced by different hormone supplementation may partially explain an apparent increase in *UBQ10p*-driven marker, since integrated single-cell root atlas (https://rootcellatlas.org/) shows that *UBQ10* expression is higher in the DZ (Supplementary Figure S13D) which is enlarged upon IAA supplementation. Because the *35Sp:NeonGreen-PTS^PEX^*^26^ expression remained unchanged in hormone-treated plants, we conclude that the ACC-triggered induction of *EBSn:mCherry-PTS1* reflects increased promoter activity rather than a shift in peroxisomal abundance.

Importantly, we aimed to enhance the sensitivity of the auxin reporter by generating an *ACE_DD2_* variant in which the auxin reporter is driven by a hybrid *DR5_DR5v2* promoter. However, the analysis of two independent lines showed no major improvement in auxin reporter sensitivity or inducibility compared to *DR5* or *DR5v2* alone (Supplementary Figures S15, S16), suggesting that the reporter performance is more strongly influenced by the 35S*minPro/35Ster* duo compatibility rather than by the number of *DR5* copies. Therefore, this design was not pursued further due to potential silencing concerns arising from the repetitive promoter architecture^51^.

To address *ACE* limitations, we designed *ACE2* featuring: (i) replacement of *3xSV40* with the *N7*^89^ NLS for efficient nuclear targeting; (ii) substitution of *TCSn:35SminPro17* with a GoldenBraid-assembled version of the original *TCSv2:35SminPro0:ΩTMV*^49^ for improved cytokinin transcriptional output; and (iii) reassignment of subcellular targeting to capitalize on compartment-specific autofluorescence profiles. In *ACE2*, cytokinin-inducible promoter *TCSv2* drives *mTagBFP2-NLS* (low background), ethylene-activated *EBSn* drives *mCherry-MTS* (negligible red autofluorescence), and auxin-controlled *DR5v2* drives single *YPet-PTS1* (Supplementary Figure S16). To enable facile crossing into publicly available T-DNA insertion mutant collections, *ACE2* versions were assembled with either BASTA or kanamycin resistance markers under *NOS*-derived or *NOS*-free, virus-sourced regulatory elements (*ACE2-M, with M = 1, 2, or 3* referring to the distinct variants of the selectable marker as depicted in Supplementary Figure S16), minimizing cross-silencing risk^66^. For each *ACE2*’s version of the selectable marker (M), two variants were produced: *ACE2-MS* (spatially *<u>s</u>egregated* reporters: peroxisome/nucleus/mitochondria) and *ACE2-MA* (spatially *<u>a</u>ggregated*, all-nuclear reporters), providing complementary tools for studies requiring either organelle-resolved or nucleus-confined multi-hormone readout.

Validation of *ACE2-3A* and *ACE2-3S* T1 lines transferred to high hormone concentrations confirmed robust activity of all three transcriptional units and efficient N7-mediated nuclear targeting, hence outperforming *ACE* in the subcellular spatial arrangement of the three reporters (Supplementary Figure S17A,B). In pursuit of reproducing the original cytokinin reporter expressing a 3xmVenus FP tandem, we compared the performance of the cytokinin-responsive transcriptional unit in the context of *ACE2* expressing one (*ACE2-3A1*) or three (*ACE2-3A3*) nucleotide-diversified copies of mTagBFP2 in unsupplemented media. We found that increasing the number of mTagBFP2 copies did not prominently improve the basal activity of the cytokinin reporter compared to its single-FP expressing counterpart (Supplementary Figure S17C). Since expressing three nucleotide-diversified copies exhausts the mTagBFP2 variants that can be included in other transcriptional units due to the high similarity between DNA sequences of the FPs employed here, we further characterized the simpler *ACE2-3A1* variant (Supplementary Figure S18A), which represents a technically improved platform ready for the full biological characterization described for *ACE*.

Under control conditions, the activity of *ACE2* reporter in two independent one-insertion homozygous T4 lines recapitulated the activity patterns described for the first-generation *ACE* reporter (Supplementary Figure S18B). Moreover, all three reporters were inducible by their exogenously applied substrate and displayed the cross-activation of the reporters in a similar manner as described for *ACE*. The lower intensities of *DR5v2* and *TCSv2* reporters induced by exogenously applied hormones were consistent with the expression of single-copy FPs in *ACE2* as opposed to the triple *YPet* and *mVenus* copies in *ACE* (Figure 2) and the original *TCSv2* (Figure 3) reporters, respectively. This new design allows the future stacking of other reporters driving the expression of additional nucleotide-diversified *YPet*, *mTagBFP2*, and *mCherry* versions with *ACE2* to expand the array of signals that can be monitored in parallel while also minimizing gene silencing. Thus, for example, parallel visualization of developmental or stress responses of interest along with that of auxin, cytokinin, and ethylene activities can shed light on the role of plant hormones in plant adaptation and phenotypic plasticity.

## Discussion

### Auxin as the convergent executor of multi-hormone root growth control

By combining the *ACE* reporter with genetic perturbation of hormone biosynthesis (*wei8*, *sur2*), transport (*aux1*), and signaling (*ahk3 cre1*, *ein2*, *ctr1*) alongside established cytokinin and auxin reporters, we dissected a hierarchical model of phytohormone crosstalk controlling primary root elongation in etiolated Arabidopsis seedlings. A central, unifying finding emerges: despite engaging distinct upstream cascades, both ethylene and cytokinin inhibit root growth predominantly through TAA1-dependent auxin biosynthesis in source cells, and IAA uptake into sink cells.

Ethylene acts via EIN2-dependent signaling to stimulate TAA1 expression in LRC and EPI source cells, enabling local IAA production that suppresses root elongation. Cytokinin engages a longer relay: it activates ethylene production and EIN2 signaling, which in turn induces TAA1 in vascular source tissues; the resulting IAA is then redistributed to LRC and EPI sink cells through an AUX1-dependent transport pathway (Figure 6). The obligate requirement for EIN2 in cytokinin-triggered auxin biosynthesis—and the insensitivity of *ein2* and *aux1* roots to *t*-Zea—positions ethylene signaling as a required intermediary between cytokinin perception and auxin-mediated root growth inhibition. This functional relay explains why root growth responses to cytokinin partially phenocopy those of ethylene and are jointly suppressed by *wei8* and *aux1*. Critically, the reduced RAM size of *ein2* and *aux1* mutants in response to *t*-Zea despite their partial resistance to root growth inhibition suggests that EIN2 and AUX1 are required for cytokinin-mediated growth inhibition but not for cytokinin-induced meristematic cell-differentiation rate^41^, consistent with previous work^26^.

Integrating our results with the known inhibitory effect of cytokinin on auxin distribution within auxin transport pathway^24,84^ is consistent with previous reports where cytokinin impairs auxin efflux and decreases AUX1 accumulation in the EPI cells within EZ but not in the LRC, hence explaining the net AUX1-dependent IAA accumulation in LRC in response to *t*-Zea (Figure 5; Supplementary Figure S7). In contrast, the low *DR5v2* activity in vasculature cells despite their role as the auxin source in response to *t*-Zea could be interpreted as minimal intracellular auxin retention in the vascular cells. However, that would need to be explained by an efficient net efflux of newly synthesized IAA into the apoplast and impaired uptake of apoplastic IAA. While the latter would be consistent with cytokinin’s inhibitory role in LAX2 accumulation^26,29^, the former would contradict the reported cytokinin-inhibited accumulation of PIN carriers in roots^24^. The previously described cell-type-specific inhibitory effect of cytokinin in auxin activity in the vascular cells^26,28,37^ (where *TCSv2* is strongly induced; Figures 3A and 6A) could explain the observed decrease of *DR5v2* activity despite active TAA1-mediated auxin biosynthesis and impaired IAA efflux.

The self-reinforcing loop in the root—where cytokinin, ethylene, and auxin mutually sustain each other’s signaling—is functionally separable from the root growth-inhibitory relay. Thus, the fact that the roots of the cytokinin receptor mutant *ahk3 cre1* respond normally to IAA despite lacking this positive feedback suggests that although some hormone interactions are absolutely required for specific responses, such as root growth inhibition, others may serve modulatory roles such as buffering signal amplitude (referred to as oscillator^26^) or coordinating developmental state transitions^24,28^.

In summary, our results are consistent with a model where cytokinin regulates auxin activity in a tissue-specific manner, with *t*-Zea inhibiting the action of newly TAA1-synthesized auxin in the vasculature that potentially accumulates over time due to impaired efflux, while local TAA1-mediated auxin biosynthesis provides enough IAA to maintain the SCN upon impaired auxin transport, as previously reported^8^. This model builds on prior evidence that ethylene promotes auxin biosynthesis^5–11^ and that cytokinin modulates auxin levels^19,21–29,43^ by providing spatial and epistatic resolution unavailable from single-pathway analyses. Critically, *ACE* enables these conclusions by revealing cross-activation patterns. Specifically, both ACC and *t*-Zea induce *DR5v2* in the root when the relevant upstream signals and transport routes are intact. This finding could suggest confounded hormone response overlap. Instead, it turns ACE into a diagnostic tool for dissecting hormone signaling interactions in the root. Distinguishing signaling pathway interactions from developmental outputs requires a specific kind of readout. It must be simultaneous, multi-hormone, and cell-type-resolved. *ACE* and *ACE2* provide exactly that.

### Generalizable principles and platform extensions

The *ACE* and *ACE2* platforms introduce a generalizable design logic for high-order multiplexed reporters. Sequence diversification of repeated genetic elements minimizes cross-silencing; subcellular targeting provides an orthogonal dimension to spectral separation, theoretically extending three fluorescent channels to nine independent readouts (Supplementary Figure S19). This framework is compatible with reporters for additional hormone classes—brassinosteroids^90^, gibberellins^91^, abscisic acid^92,93^, jasmonate^94^, salicylate^95^, strigolactones^96^—for which *cis*-regulatory DNA elements have been identified and single-cell transcriptomics continue to refine, making a nonuple or higher-order hormone reporter a tractable engineering goal. The newly developed GB-compatible stacking platform from our laboratory^97^ further facilitates assembly of large multi-transcriptional-unit constructs.

The dose-dependent, biphasic effect of IAA on *EBSn* activity—inhibition at 50 nM, activation at 1 µM (Figures 2–5; Supplementary Figures S4A, S5A, S6, S7A, S9A,B, S14, S16)—reveals that the auxin–ethylene interaction is governed by concentration thresholds rather than monotonic proportionality. Quantitative dissection of this non-linearity using *ACE/ACE2* combined with ethylene biosynthesis mutants will be important for understanding how the ratio—not merely the absolute level—of hormone activity shapes developmental decisions, an assessment that single-hormone reporters could not enable in isolation.

Beyond root biology, *ACE*’s capacity to correlate spatially resolved hormone activity with cell identity positions it as a natural complement to single-cell transcriptomics developmental atlases^98^. In the longer term, coupling multi-hormone reporters with cell-type-specific perturbations could reveal how developmental competence states—including those relevant to callus differentiation and regeneration in recalcitrant crop species—are encoded in the dynamic balance of phytohormone activities.

## Methods

### Biological material

All Arabidopsis genotypes are in Col-0 background. Except for *ACE*- and *ACE2*-harboring lines, all reporters (*DR5*:*GFP*^46^, *TCSv2:3xmVenus-N7*^49^, [*35Sp:NeonGreen-PTS^PEX^*^26^] - [*UBQ10p:mRuby3-PTS1*]^88^, *wei8-2 TAA1p:YPet-gTAA1*^8,81^) and mutant backgrounds (*wei8-1^7^*, *sur2^5^, tir1*^82^, *aux1-7*^77^, *ein2-5*^99^, *ctr1-1*^80^, *ahk3-3 cre1-12*^75^) were previously described.

### Growth conditions

Seeds were surface-sterilized in 50% (v/v) commercial bleach supplemented with 0.01% (v/v) Triton X-100 for 5 min, followed by four washes with sterile deionized water. Sterilized seeds were stratified for 3 days at 4 °C prior to the start of each assay. Seeds were resuspended in sterile 0.6% (w/v) low-melting-point agarose and sown aseptically on the surface of sterile control medium containing 4.33 g L⁻¹ Murashige and Skoog salts, 10 g L⁻¹ sucrose, and 6 g L⁻¹ Bacto Agar (unless otherwise stated), pH 6.0 (1 M KOH). Germination was induced by 2 h of continuous white light at room temperature, and plates were placed horizontally in a dark growth chamber at 22 °C for 3 days prior to imaging. For organ measurements following exogenous hormone supplementation, seedlings were arranged horizontally on plain agar plates and imaged with an Epson Perfection V600 scanner; images were processed in FIJI/ImageJ. To assess reporter inducibility, high concentrations of IAA, ACC, or *t*-Zea were applied as described in their respective figure captions (Figure 2; Supplementary Figures S14, S15, S16, S18). To compare *ACE* activity and inducibility across genetic backgrounds, hormone concentrations were adjusted to achieve ∼50% reduction in WT root length (Supplementary Figure S3) as follows: [ACC] = 0.2 µM, [*t*-Zea] = 10 µM; [IAA] = 50 nM. Leaves of four-week-old Arabidopsis rosettes were collected from plants germinated and grown for 7 days on AT plates under continuous light, then transferred to soil (1:1 mix of Sun Gro Horticulture Professional Growing Mix and Jolly Gardner PRO-LINE C/B Growing Mix) in standard 4 × 6 flats and grown for 3 additional weeks under long-day conditions (16 h light/8 h dark, 22 °C, fluorescent bulbs EIKO F54T5/HO/850) prior to imaging.

Seeds from *Nicotiana benthamiana* were surface-sterilized for 15 min using 50% bleach spiked with 0.01% of Triton X-100^8^ and washed five to seven times with deionized water under sterile conditions. Sterilized seeds were then grown in autoclaved soil in a walk-in growth chamber at 25 °C of temperature under a 16 h/8 h light/dark cycle^51^. Four-week-old plants were then used for leaf agroinfiltration assays. *ACE* plasmids were transformed into homemade electrocompetent *Agrobacterium tumefaciens* GV3101 cells as described^100^, and the resulting colonies were validated by PCR. Positive *Agrobacterium* clones were then tested by *N. benthamiana* leaf agroinfiltration assays, performed as described^101^. For the triple auxin-cytokinin-ethylene induction, 10 µM *t*-Zea (PhytoTechnology Lab) and 10 µM ACC (PhytoTechnology Lab) were mixed with the agroinfiltration solution and inoculated simultaneously with the plasmid. About 30 min before the 72h post-agroinfiltration, a solution with 0.1 µM IAA (PhytoTechnology Lab) spiked with 0.001% of Triton X-100 was painted with a brush over the agroinfiltrated area. After that period, leaves were used to check for fluorescence using a DFC365 FX camera and a Zeiss Axioplan microscope (20x objective; 100% excitation power; exposure time: 600 ms (YPet and mCherry) and 800 ms (mTagBFP*)).

### GB-compatible library

An *ad hoc* script was used to generate three Arabidopsis codon-optimized CDSs for each fluorescent protein and subcellular-targeting peptide variant (DOI: 10.5281/zenodo.21797666). Two additional scripts generated *35Sp*-derived minimal promoters (*minPro*) (DOI: 10.5281/zenodo.21797744) and terminators (*ter*) (DOI: 10.5281/zenodo.21797824) with intact TATA-box and polyadenylation site positions, respectively, and preserved GC content. All DNA parts were amplified by PCR with primers presented in Supplementary Table S1, subcloned into the GB-compatible entry plasmid *pUPD2,* confirmed by Sanger sequencing, and individually validated in the context of tester transcriptional units in transient expression assays in 4-week-old *Nicotiana benthamiana* leaves assessing luminescence induction strength (Supplementary Figure S1A-E), fluorescent protein activity, and subcellular targeting efficiency (Supplementary Figure S1F) as described^51^. The assembly of these entry clones into functional transcriptional units, and then into larger multi-transcriptional modules, was done following GB2.0 rules^102^ as previously described^51,52^, and final assemblies were confirmed by whole-plasmid sequencing.

As repetitive sequences are often incompatible with commercial DNA synthesis, we employed a cloning strategy different from the standard GB2.0 aimed to circumvent this issue for cytokinin-inducible *TCSn* promoter, auxin-inducible *DR5*-related promoters, and full-length CVMV promoter. Cytokinin-inducible *TCSn* promoter was initially cloned alongside the *35SminPro0* into the *pUPD2* entry vector from *pUC18_TCSn:35SminPro0:LUC:NOSter*^48^ (Supplementary Figure S20A,B). To make this inducible promoter fully modular and compatible with the collection of minimal promoters (*minPro*), it was recloned a second time using a non-GoldenBraid type IIS restriction enzyme, AcuI (NEB CutSmart R0641S). As the repetitive nature of short *TCSn* repeats prevents designing oligos directly within its sequence, we leveraged the ability of AcuI to cut 16bp away from the recognition site and leave 2bp sticky ends: oligos on the 3’-end of *TCSn* F2, R2 and R3, Supplementary Table S1A) were designed to create a 2bp sticky end cutting over the 5’-end of *35SminPro0* sequence (Supplementary Figure S20C,D). These oligos also included the recognition site for BsmBI (NEB CutSmart R05580L) and the *pUPD2* cloning overhang as previously described for other hormone-inducible promoters^51,52^ (Supplementary Figure S20C,D). The *minPro*-free *TCSn* sequence was thus obtained by PCR amplification and a restriction-ligation cycle using AcuI and T4 DNA ligase (NEB M0202L) (Supplementary Figure S20E) and subcloned into *pUPD2* entry vector following a second restriction-ligation cycle as described for the GoldenBraid cloning strategy^98,51^ (Supplementary Figure S20F).

Auxin-inducible proximo-distal (A1-A2) promoters *DR5_×10_*, *DR5v2_×10_* and *DR5_×10_-DR5v2_×10_* were cloned from *pGREEN II_DR5×10-dTomato / DR5v2-3nGFP*^47^ (Supplementary Figure S21). To amplify the individual proximo-distal (A1-A2) *DR5_×10_* and *DR5v2_×10_* promoter sequences, both forward oligos (F1 and F2, Supplementary Table S1A) harbored the 5’-A1 GB code and a 3’-end sequence complementary to the first *DR5* or *DR5v2* repeat sequence, respectively, to ensure specificity (Supplementary Figure S21A). The reverse oligo (R1, Supplementary Table S1A) was shared by both promoters, as it was designed over the common 3’-multisite cloning sequence, and it harbors the 3’-A2 GB code (Supplementary Figure S21A). For amplifying the combined proximo-distal (A1-A2) *DR5_×10_-DR5v2_×10_* promoter sequence, individual *DR5_×10_* and *DR5v2_×10_* sequences were amplified using a similar approach changing the GB code of F2 and R2 oligos: the new reverse oligo for the *DR5_×10_* sequence harbored a 3’-A1 GB code (R2, Supplementary Table S1A), and the new forward oligo for the *DR5v2_×10_* sequence included a 5’-A2 GB code instead (F3, Supplementary Table S1A, Supplementary Figure S21B). Each of these oligos were designed to also include a recognition site for BsmBI and the *pUPD2* cloning overhang as was previously described^51^.

The full-length CVMV promoter sequence was split into two GBlocks for commercial synthesis to comply with the quality thresholds established by the vendor (Integrated DNA Technology). Two oligo pairs (F1, F2, R1, and R2, Supplementary Table S1A) were designed to amplify both fragments independently and harbored BsmBI recognition sites in R2 and F2 oligos to create 4bp sticky ends to allow scar-less fragment assembly into *pUPD2* clone during the regular restriction-ligation cycle (Supplementary Figure S21C).

### Quantification of the transcriptional activity of *minPro* and *ter* libraries by dual-luciferase assays

Four leaf-disk pools (0.25 inch diameter, four leaves each) per construct (Supplementary Figure S1B,D) were flash frozen in liquid nitrogen and ground with 100 µL of 1-mm glass beads for 4-5 seconds in a Vivadent dental shaker, then mixed with 250 µL room-temperature 1x Passive Lysis Buffer (Promega). Samples were vortexed until thawed, kept on ice and in the dark, and centrifuged at maximum speed for 15 minutes at 4 °C to pellet tissue debris. 100 µL of each supernatant was transferred to a well in a 96-well PCR plate for multichannel pipetting during the assay. Luciferase activity was measured in a black 96-well plate using a 96-well GloMax luminometer with automatic dual injectors. To limit signal degradation, only four 10-µL samples were loaded at a time *via* multichannel pipette, with remaining samples kept dark on ice. The luminometer injected 40 µL LARII (Promega) to measure firefly luciferase (LUC), followed by 40 µL Stop&Glo to quench LUC and activate Renilla luciferase (REN), with a 2-second integration time and 10-second read time for both reactions. These experiments were repeated three times.

### Quantification of the transcriptional activity of the library of full viral promoters by dual-luciferase assays

For each construct, three independent biological replicates were collected from three independently infiltrated plants. Each biological replicate (a pool of two 0.25-inch-diameter leaf disks excised from the same infiltrated leaf of a single plant) (Supplementary Figure S1E) was flash frozen in liquid nitrogen, mixed with 100 µL ice-cold thawed 1x Passive Lysis Buffer (Promega) and ground with sterile blue pestles. Samples were on ice for about 15-20 minutes while processing the different replicates from each construct, allowing the debris to pellet. 20 µL of each supernatant was transferred to a well from an opaque 96-wells plate and kept on ice and in darkness. For the luminescence quantification, 100 µL LARII (Promega) were loaded at a time in each well using a multichannel pipette to measure firefly luciferase (LUC). Likewise, 100 µL Stop&Glo were then loaded at once to each well using a multichannel pipette to quench LUC and activate Renilla luciferase (REN). Reads were measured for 10 min with an interval of 1 min between reads and 10 sec of initial orbital shaking. Luminescence and Kinetics were used as read mode and type, respectively, in a SpectraMax®iD5 device with SoftMax® Pro 7.0.3 software (Molecular Devices, USA) (unmentioned parameters were set as default).

### Image acquisition and data analysis

Flower organs and expanded siliques were dissected under a Nikon SMZ645 stereomicroscope using Dumoxel Style 5 forceps. Embryos were isolated from expanded siliques, mounted in 15% glycerol, and imaged. Fluorescence from whole seedlings, flower organs, and siliques was acquired on a Leica Thunder Imager M205FA with Leica DMC6200 (bright field) and DFC9000 sCMOS (fluorescence) cameras, using fixed filter sets: Blue (#10450571; excitation ∼395–415 nm, emission 435–485 nm), GFP (#10447408; excitation 450–490 nm, emission 500–550 nm), and mCherry (#10450195; excitation 540–580 nm, emission 592–667 nm). Embryo fluorescence imaging used standard Zeiss HE filter sets (DAPI/Filter Set 49 for mTagBFP2; GFP/Filter Set 38 HE for YPet; DsRed/Filter Set 43 HE for mCherry). Confocal imaging was performed on a Zeiss LSM 880 with Plan-Apochromat 20×/0.8 NA (air) and 63×/1.4 NA (oil-immersion) objectives; pinhole = 1 AU. Images were acquired at 512 × 512 (20×, Z-steps = 1 µm) or 1024 × 1024 (63×) pixels with line-sequential scanning and frame averaging (2× at 20×; 4× at 63×). Fluorophores were excited at 405, 488, and 561 nm; emission was detected by PMT and spectral detectors (mTagBFP2: 415–489.8 nm; YPet: 499–561.2 nm; mCherry: 561.2–685.6 nm) at 16-bit depth. All imaging parameters were held constant within experiments. Five or more biological replicates for every given genotype x treatment combination were looked at prior to the acquisition of a representative specimen.

Images were analyzed in FIJI/ImageJ, using the Z-stack *maximum intensity* projection method where applicable with uniform brightness/contrast adjustments applied within each experiment. Data visualization and statistical analyses were performed in RStudio using the *car*^103^, *pwr*^104^, *rstatix*^105^, *dunn.test*^106^, and *multcompView*^107^ packages. Outliers were removed by Tukey’s criteria (x < Q1 − 1.5 × IQR or x > Q3 + 1.5 × IQR). Normality and homoscedasticity were assessed by Shapiro–Wilk^108^ and Levene’s^109^ tests, respectively. Statistical power was calculated as 1 − β. For homoscedastic datasets with power > 0.8, one-way ANOVA with Tukey’s post hoc test (α = 0.05) was used regardless of normality. Non-conforming datasets were log₁₀-transformed and re-assessed; datasets still failing parametric assumptions were analyzed by Kruskal–Wallis^110^ with Dunn’s multiple comparisons^111,112^ for α = 0.05.

## Supplementary Material Legends

**Supplementary Table S1.** *pUPD2* entry clones and selectable markers created for the different GB parts assembled into transcriptional reporters. (A) Hormone-inducible promoters. GB: GoldenBraid. *10xDR5* and *10xDR5v2* come from *pGREEN II_DR5×10-dTomato* / *DR5v2-3nGFP*^47^ and were independently amplified *via* PCR using primers containing GoldenBraid (GB) overhangs for parts A1 and A2, respectively. Both amplicons were then assembled in the *pUPD2* vector in a single-pot reaction, with the PCR fragment *DR5ₓ₁₀* (GB A1-A2) amplified with primers F1+R1 and the PCR fragment *DR5v2ₓ₁₀* (GB A1-A2) amplified with primers F2+R1 to generate two alternative proximo-distal promoters (Supplementary Figure S21A). The PCR fragment *DR5ₓ₁₀* (GB A1) was amplified with primers F1+R2, whereas the PCR fragment *DR5v2ₓ₁₀* (GB A2) was amplified with primers F2+R3, and the two PCR products were combined together to generate a third proximo-distal promoter (Supplementary Figure S21B). *TCSn* came from *pUC18_TCSn:35SminPro0:LUC:NOSter*^48^. The repetitiveness of the *TCSn* sequence necessitated its subcloning as *pUPD2_TCSn*:*35SminPro0*, with primers F1+R1 used to amplify the promoter. To make the *TCSn* construct fully modular and compatible with different *minPro* of the collection, the *35SminPro0* was removed with the help of a non-GB Type IIS enzyme AcuI by leveraging PCR1 that used primers F1+R3 and PCR2 that relied on primers F2+R2, and combining the two PCR products in *pUPD2* using AcuI, BsmBI, and DNA ligase (Supplementary Figure S20). Original *TCSv2:35SminPro0:TMV* was described in^49^. TMV: Tobacco mosaic virus translational enhancer. (B) Full promoters from the Cauliflower mosaic virus *35S* (CaMV *35Sp*; 1027 bp)^51^, the Strawberry vein banding virus Sgt VI^61,62^ (SVBV VIp; 988 bp), the Figwort mosaic virus M3 strain Sgt VI (*19S*)^63,64^ (FMV *19Sp*; 416 bp), and the Cassava vein mosaic virus Flt^63,65^ (CVMV *Fltp*; 2895 bp). DNA sequences were amplified from GBlock synthetic DNA. The synthesis of *fullProCVMV* was split into two GBlocks to be commercially synthesized, with the two fragments amplified (PCR1 used primers F1+R2 and PCR2 relied on primers F2+R1) and combined during regular GB subcloning protocol into *pUPD2* (Supplementary Figure S21C). Sgt: subgenomic transcript; Flt: full-length transcript; VI: sixth open reading frame (ORF). (C) Core promoters. Newly designed DNA variants of the minimal promoter from CaMV *35S* gene^51,52^ were amplified from GBlock synthetic DNA composed of multiple (three to ten) DNA parts flanked by part-specific GB codes and stacked together as a cost-cutting measure. Construct-specific oligos that include a tail with the GB cloning overhang were designed to amplify *minPro* sequences prior to their *pUPD2* subcloning. Alternatively, sequences of oligos from our lab collection were used as sequence stuffers in the design of the multisequence GBlocks which enabled selective PCR amplification of the DNA fragments flanked by the *pUPD2* cloning overhang (when allowed by vendor’s synthesis thresholds). Minimal promoters from FMV SxD strain Sgt VI (*19S*) ^58,59^ and Horseradish latent virus (HRLV) Sgt VI^60^ deduced from previously described full promoters were also amplified from GBlock synthetic DNA. Sgt: subgenomic transcript; VI: sixth open reading frame (ORF). (D) Subcellular localization signals, N-terminal/C-terminal. *Mitochondrial Targeting Signal (MTS)* derived from the *CYTOCHROME C-OXIDASE SUBUNIT 4*^70,71^ was amplified from GBlock synthetic DNA. The *MitCox4* sequence was synthesized with the GB cloning overhangs and later amplified with oligos from our lab collection used as stuffers in the synthesis of the multisequence GBlocks. *Peroxisomal Targeting Signal 1 (PTS)* was previously described^74^. The original *Nuclear Localization Signal (NLS) N7* was previously described^89^. (E) Reporter proteins. Initially, mTagBFP* was used in *ACE* as blue fluorescent protein (hybrid version of mTagBFP that contains an mTagBFP2-like amino acid substitution in the mTagBFP chromophore moiety but does not include the mTagBFP2 N-terminus). Later, it was upgraded to full mTagBFP2^69^ in *ACE2*. FP DNA sequences were amplified from GBlock synthetic DNA. The mCherry C-terminal CDS sequence was synthesized with the GB cloning overhangs and amplified using oligos from our lab collection used as stuffers in the synthesis of multisequence GBlocks. (F) Terminators. Newly designed DNA variants of the terminator from CaMV *35S* gene^51,52^ were amplified from GBlock synthetic DNA composed of multiple (three to ten) DNA parts flanked by part-specific GB codes and stacked together as a cost-cutting measure. Individual terminator amplification was carried out in a similar manner as described above for *minPro*. (G) *NOS*- and *35S*-free selectable markers. *CVMV Fltp*: 2895 bp-promoter from the CVMV *full-length transcript*^63,65^ was amplified from GBlock synthetic DNA. *FMV 19Sp*: 416 bp-promoter from the FMV M3 strain Sgt VI *(19S)*^63,64^ was amplified from GBlock synthetic DNA. *HSP18.2ter*: terminator from *Arabidopsis thaliana HEAT SHOCK PROTEIN18.2*^116^. (H) Other *pUPD2* clones previously reported. A1-C1 (TU) refers to the full transcriptional unit (*promoter:CDS:terminator*) subcloned into an entry vector, *pUPD*, from the GoldenBraid collection (GB0023 or GB0184). They were all described in a previous work^52^. (I) The GoldenBraid website is given as reference for the *pUPD* entry clone GB0096 from the GoldenBraid collection. The *minPro*-less *35S* proximodistal promoter (A1-A2) sequence was amplified from the *pUPD* entry clone GB0030 (*pUPD_fullPro35S*) from the GoldenBraid collection. The *RENILLA* CDS sequence from the clone GB0109 (*pEGB_35S:REN:tNOS*) was redomesticated into *pUPD2* entry clone to modify the flanking GB codes and make them compatible with the other DNA elements to assemble.

**Supplementary Figure S1.** Collection of minimal promoters (*minPro*) and terminators (*ter*) assessed for transcriptional activity in transient expression assays in agroinfiltrated *Nicotiana benthamiana* leaves. (A) DNA sequence comparison of 22 minimal promoters (twenty of which were derived from CaMV *35Sp* plus those derived from FMV DxS strain subgenomic transcript VI *(19S)* or HRLV full-length transcript) relative to CaMV *35SminPro0* (*minPro0*) visualized using MEGAX^117^ software without sequence alignment. The position of the *TATA* box (highlighted in yellow) is preserved in all *35Sp*-derived promoters. (B) Construct structures and normalized activities of fifteen minimal promoters relative to *minPro0* (black-edged vertical bar; horizontal black line at y-axis=1) were assessed by transient assays in *Nicotiana benthamiana* leaves agroinfiltrated with [*DisProx35Sp:minProX:LUCIFERASE:ter0*] *–* [*35Sp:RENILLA:NOSter*]. Vertical bars represent the average LUC/REN ratios, and data points indicate individual pseudoreplicates (n=4). *DisProx35Sp* refers to CaMV *minPro0-less 35Sp*. For visual reference, significant differences between a given *minProX* and *minPro0* are highlighted in bold letters, with magenta and green indicating lower or higher activity, respectively. (C) DNA sequence comparison of the twenty CaMV *35Ster*-derived terminators compared to *35Ster0* (*ter0*) visualized using MEGAX^117^ without sequence alignment. The polyadenylation signals are highlighted in yellow and their position is preserved in all terminators. (D) Construct structures and normalized activities of twenty terminators relative to *ter0* (black-edged vertical bar; horizontal black line at y-axis=1) assessed by transient assays in *Nicotiana benthamiana* leaves agroinfiltrated with [*35Sp:LUCIFERASE:terX*] *–* [*35Sp:RENILLA:NOSter*]. Vertical bars represent the average LUC/REN ratios and data points indicate individual pseudoreplicates (n=4). For visual reference, significant differences between a given *terX* and *ter0* are highlighted in bold letters, with magenta and green indicating lower or higher activity, respectively. (E) Construct structures and normalized activities of three viral-sourced promoters relative to CaMV *35Sp* (black-edged vertical bar; horizontal black line at y-axis=1) were assessed by transient assays in *Nicotiana benthamiana* leaves agroinfiltrated with [*Full_proX:LUCIFERASE:ter0*] *–* [*35Sp:RENILLA:NOSter*]. *Full_proX:* full promoters from the Strawberry vein banding virus (SVBV; 988 bp) Sgt VI^61,62^, FMV M3 strain (416 bp) *19S* (Sgt VI)^63,64^, or Cassava vein mosaic virus (CVMV; 2895 bp) Flt^63,65^ (Sgt: subgenomic transcript; Flt: full-length transcript; VI: sixth open reading frame). Vertical bars represent the average LUC/REN ratio, and data points indicate individual pseudoreplicates (n=3) representing technical variability. Different letters denote statistically significant differences between means (one-way ANOVA, ɑ=0.05). (F) Transient expression of *ACE* reporter in *N. benthamiana* leaves upon hormone infiltration. ACC = 10 µM 1-aminocyclopropane-1-carboxylic acid, *t*-Zea = 10 µM *trans*-zeatin, IAA = 0.1 µM indole-3-acetic acid. *PTS1: Peroxisomal Targeting Signal 1 (N-KSRM-C). MTS: Mitochondrial Targeting Signal (MitCox4); NLS: Nuclear Localization Signal (3xSV40)*. mTagBFP*: Hybrid version of mTagBFP that contains mTagBFP2-like amino acid substitution in mTagBFP chromophore moiety but does not include the mTagBFP2 N-terminus.

**Supplementary Figure S2.** Subcellular localization of the *ACE* reporter in Arabidopsis root tip cells. (A) Two independent insertion lines of the *ACE* reporter (*L9.8* and *L16.2*) reveal subcellular localization of mCherry and mTagBFP* consistent with peroxisomes (Px) and mitochondria (Mt), respectively, whereas YPet signal is not confined to the nucleus (Nu). (B) Comparison of mCherry localization in the *ACE* reporter with the previously described dual peroxisomal marker [*35Sp:NeonGreen-PTS^PEX^*^26^] - [*UBQ10p:mRuby3-PTS1*]^88^ imaged in separate roots supports peroxisomal targeting of mCherry. White boxes represent the area of the root that is magnified within the same panel for a given genotype and reporter. Three-day-old etiolated seedlings were germinated on unsupplemented media. All microscopy images were acquired in the same experimental session. Confocal microscopy images show stacked pictures from the equatorial plane at the quiescent center to the cortical plane of the lateral root cap. Scale bar applies to all images within the panel. ET: ethylene reporter (*EBSn:mCherry-PTS1*); CK: cytokinin reporter (*TCSn:MTS-mTagBFP\**); AUX: auxin reporter (*DR5v2:3xSV40-3xYPet*); mTagBFP*: Hybrid version of mTagBFP that contains mTagBFP2-like amino acid substitution in mTagBFP chromophore moiety but does not include mTagBFP2 N-terminus; BF: bright field; WT: wild-type.

**Supplementary Figure S3.** Quantitative analysis of relative organ sizes in three-day-old etiolated wild-type (WT) and mutant seedlings of Arabidopsis defective in ethylene, cytokinin, or auxin biosynthesis, perception, signaling, or transport in response to exogenous hormone treatments. All genotype x treatment combinations were grown in the same experimental session. Black dots represent independent organ measurements of individual seedlings and horizontal red bars across them represent the mean values. Numbers at the bottom of the plot (in grey) represent the number of biological replicates (n) for a given organ and treatment. Different letters indicate statistically significant differences among genotypes for a given organ within each treatment. Multiple comparison analyses were performed as described in the methods section under *Image acquisition and data analysis*. ACC = 0.2 µM 1-aminocyclopropane-1-carboxylic acid; *t*-Zea = 10 µM *trans*-zeatin; IAA = 50 nM indole-3-acetic acid.

**Supplementary Figure S4.** Detailed view of AHK3- and CRE1-dependent responses to exogenous hormone treatments in signaling pathway crosstalk in Arabidopsis roots. (A) Equatorial Z-stack of *ACE* and *TCSv2* activities in longitudinal sections of the same Arabidopsis roots displayed in Figure 3A. Scale bar applies to all images within the panel. ACC = 0.2 µM 1-aminocyclopropane-1-carboxylic acid; *t*-Zea = 10 µM *trans*-zeatin; IAA = 50 nM indole-3-acetic acid; ET: ethylene reporter (*EBSn:mCherry-PTS1*); CK: cytokinin reporter (*TCSv2:3xmVenus-N7*)^49^; AUX: auxin reporter (*DR5v2:3xSV40-3xYPet*); BF: bright field; DZ: differentiation zone; EZ: elongation zone; TZ: transition zone; RAM: root apical meristem. (B) Updated Arabidopsis hormone signaling pathway interaction model based on *ACE* and *TCSv2* activities and phenotypic responses of *ahk3 cre1* roots to exogenous application of hormones. Based on *TCSv2* and *ACE* reporters, IAA and ACC induce *TCSv2* activity. The *ahk3 cre1* mutation prominently reduces ACC- and IAA-triggered upregulation of *TCSv2*, suggesting that ethylene and auxin work upstream or at the level of cytokinin receptors.

**Supplementary Figure S5.** Detailed view of TAA1-dependent responses to exogenous hormone treatments in signaling pathway crosstalk in Arabidopsis roots. (A) Equatorial Z-stack of *ACE* activity in longitudinal sections of the same Arabidopsis roots displayed in Figure 4A. Scale bar applies to all images within the panel. ACC = 0.2 µM 1-aminocyclopropane-1-carboxylic acid; *t*-Zea = 10 µM *trans*-zeatin; IAA = 50 nM indole-3-acetic acid; ET: ethylene reporter (*EBSn:mCherry-PTS1*); AUX: auxin reporter (*DR5v2:3xSV40-3xYPet*); BF: bright field; DZ: differentiation zone; EZ: elongation zone; TZ: transition zone; RAM: root apical meristem. Root zone boundaries are approximate. (B) Updated Arabidopsis hormone signaling pathway interaction model based on *ACE* activity and phenotypic responses of *wei8* roots to exogenous application of hormones. *wei8* mutant shows normal ethylene signaling, but its ACC-induced auxin activity is suppressed. Given that *wei8* shows mild insensitivity to ACC (Figure 4B), the results suggest that ethylene-decreased root length involves TAA1-mediated increase in auxin biosynthesis in response to ACC (supported by Supplementary Figure S8). Furthermore, *wei8* also suppresses *t*-Zea-mediated increase in auxin signaling, consistent with exogenously supplied cytokinin leading to a boost in TAA1-mediated auxin biosynthesis (supported by Supplementary Figure S8).

**Supplementary Figure S6.** High endogenous concentration of auxin in *sur2* sensitizes the Arabidopsis root to the perception of ethylene and cytokinin, but it does not translate into prominently higher basal *EBSn* activity in the roots of three-day-old etiolated seedlings. (A) Half-root Z-stacks of an auxin overproducing mutant *sur2* show stronger basal activity of auxin and ethylene reporters. Moreover, high auxin in *sur2* increases the sensitivity of the root to all exogenously applied hormones tested. Arrowheads point at root sections where reporter expression differs between treatments or genotypes. (B) Equatorial Z-stack of *ACE* activity in longitudinal sections of the same Arabidopsis roots displayed in Figure 6A. Scale bar applies to all images within the panel. (C) *sur2* roots show stronger sensitivity to exogenous cytokinin and mildly, yet significantly stronger sensitivity to exogenous IAA. Despite the ethylene reporter in *sur2* roots being more sensitive than WT to exogenous ACC, *sur2* relative root length is similar to that of wild type (WT). Quantitative comparison and statistical assessment of hormone-specific inhibitory effect on growth between WT and mutants are available in Supplementary Figure S3. All microscopy images were acquired in the same experimental session. Half root Z-stack (A) confocal microscopy images show stacked pictures from the equatorial plane at the quiescent center to the cortical plane of the lateral root cap, while those labelled as “Equatorial” (B) show stacked pictures immediately surrounding the quiescent center. Scale bar applies to all images within the panel. ACC = 0.2 µM 1-aminocyclopropane-1-carboxylic acid; *t*-Zea = 10 µM *trans*-zeatin; IAA = 50 nM indole-3-acetic acid. ET: ethylene reporter (*EBSn:mCherry-PTS1*); AUX: auxin reporter (*DR5v2:3xSV40-3xYPet*); BF: Bright field. DZ: differentiation zone; EZ: elongation zone; TZ: transition zone; RAM: root apical meristem. Root zone boundaries are approximate.

**Supplementary Figure S7.** Detailed view of *EIN2-* and *AUX1*-dependent responses to exogenous hormone treatments in the signaling pathway crosstalk in Arabidopsis roots. (A) Equatorial Z-stack of *ACE* activity in longitudinal sections of the same Arabidopsis roots displayed in Figure 5A and C. (B) Updated Arabidopsis hormone signaling interaction model based on *ACE* activity and phenotypic responses of *ein2 ACE* roots to exogenous application of hormones. IAA-dependent induction of *EBSn* activity is suppressed in *ein2*, suggesting that auxin signaling can also operate upstream of ethylene inducing its signaling activity in an *EIN2*-dependent manner, rather than supporting the scenario where a subset of auxin-inducible genes trans-activate ethylene-inducible genes. Furthermore, cytokinin-dependent induction of ethylene and auxin signaling is also suppressed in *ein2*, suggesting that in WT roots, cytokinin stimulates ethylene biosynthesis or ethylene perception, hence leading to the increase of *EBSn* activity and a boost of ethylene-dependent auxin biosynthesis, rather than cytokinin directly stimulating auxin biosynthesis. (C) Equatorial Z-stack of *ACE* activity in longitudinal sections of the same Arabidopsis roots displayed in Figure 5E. (D) Updated Arabidopsis hormone signaling pathway interaction model based on *ACE* activity and phenotypic responses of *aux1* roots to exogenous application of hormones. Despite having normal ethylene signaling and fully functional auxin biosynthesis machinery, *aux1* roots fail to increase auxin signaling upon ACC or *t*-Zea supplementation, suggesting that ethylene- or cytokinin-induced local auxin biosynthesis requires normal auxin import from source cells (where TAA1 reporter accumulates; Supplementary Figure S8) to sink cells (showing *DR5v2* maxima in WT plants). However, a group of cells surrounding the quiescent center show increased auxin activity upon exposure to exogenous hormones, suggesting that increased auxin might be channeled towards the root tip through other transportation paths. Scale bar applies to all images within the panel. ACC = 0.2 µM 1-aminocyclopropane-1-carboxylic acid; *t*-Zea = 10 µM *trans*-zeatin; IAA = 50 nM indole-3-acetic acid; ET: ethylene reporter (*EBSn:mCherry-PTS1*); AUX: auxin reporter (*DR5v2:3xSV40-3xYPet*); BF: bright field; DZ: differentiation zone; EZ: elongation zone; TZ: transition zone; RAM: root apical meristem. Root zone boundaries are approximate.

**Supplementary Figure S8.** TAA1 accumulation pattern in Arabidopsis roots responds differently to the application of distinct exogenous hormones in *wei8-2 TAA1p:YPet-gTAA1*. (A) Half-root Z-stacks showing that TAA1 accumulation in roots is consistently induced by all hormones relative to untreated control. (B) Equatorial section of the roots displayed in panel A. While *t*-Zea induces TAA1 accumulation mainly in the vasculature of the primary root, ACC and IAA induce TAA1 accumulation also in the EPI and LRC, with ACC triggering stronger TAA1 accumulation in the TZ and EZ than IAA. Arrowheads point at specific cell types where reporter expression differs between treatments. All microscopy images were acquired in the same experimental session. Half root Z-stack (A) confocal microscopy images show stacked pictures from the equatorial plane at the quiescent center to the cortical plane of the lateral root cap, while those labelled as “Equatorial” (B) show stacked pictures immediately surrounding the quiescent center. Scale bar applies to all images within the figure. ACC = 0.2 µM 1-aminocyclopropane-1-carboxylic acid; *t*-Zea = 10 µM *trans*-zeatin; IAA = 50 nM indole-3-acetic acid; TAA1: TRYPTOPHAN AMINOTRANSFERASE OF ARABIDOPSIS1; LRC: lateral root cap; EPI: epidermis; TZ: transition zone; DZ: differentiation zone; RAM: root apical meristem. BF: bright field. Root zone boundaries are approximate.

**Supplementary Figure S9.** Constitutively active ethylene signaling in *ctr1* translates into more sensitized roots to exogenous hormones in three-day-old etiolated Arabidopsis seedlings. (A) Half-root Z-stacks showing that the disruption of the negative regulator *CTR1*, which exhibits the phenotype of ethylene-treated plants in the absence of ethylene, leads to strong *EBSn* reporter activity, higher basal auxin reporter activity in control conditions, and stronger induction of auxin reporter in response to exogenously applied hormones. (B) Equatorial sections of the roots displayed in panel A. (C) Seedling phenotypes in the presence or absence of hormone supplementation. Despite obvious differences in reporters’ activity in response to exogenously applied hormones, *ctr1* shows very mild organ size changes in response to these growth regulators, suggesting a near saturation of the physiological and developmental responses to hormonal signals. Quantitative comparison and statistical assessment of hormone-specific inhibitory effect on growth between WT and mutants are available in Supplementary Figure S3. All microscopy images were acquired in the same experimental session. Half root Z-stack (A) confocal microscopy images show stacked pictures from the equatorial plane at the quiescent center to the cortical plane of the lateral root cap, while those labelled as “Equatorial” (B) show stacked pictures immediately surrounding the quiescent center. Scale bar applies to all images within the panel. ACC = 0.2 µM 1-aminocyclopropane-1-carboxylic acid; *t*-Zea = 10 µM *trans*-zeatin; IAA = 50 nM indole-3-acetic acid. ET: ethylene reporter (*EBSn:mCherry-PTS1*); AUX: auxin reporter (*DR5v2:3xSV40-3xYPet*); BF: bright field. Root zone boundaries are approximate.

**Supplementary Figure S10.** Expression of the *ACE* reporter in leaves of four-week-old Arabidopsis rosettes grown under long-day conditions. (A) *ACE* expression at the region of the leaf marked on the left-hand side cartoon (Top row: center of the leaf blade near the vasculature, Middle row: edge of the leaf, Bottom row: tip of the leaf) in adult cotyledons (left column), first true leaf of the rosette (middle column), and the youngest fully expanded leaf (right column). (B) Digital magnification of images displayed in panel A showing peroxisome-targeted ethylene reporter’s activity (arrows) between chloroplast autofluorescence activity. Confocal microscopy images show stacked pictures from mesophyll to epidermis cell planes. All microscopy images were acquired in the same experimental session. Confocal microscopy images show stacked pictures spanning the full depth of leaf epidermal cells, extending into the palisade parenchyma. Scale bar applies to all images within the panel. ET: ethylene reporter (*EBSn:mCherry-PTS1*); CK: cytokinin reporter (*TCSn:MTS-mTagBFP\**); AUX: auxin reporter (*DR5v2:3xSV40-3xYPet*); mTagBFP*: Hybrid version of mTagBFP that contains mTagBFP2-like amino acid substitution in mTagBFP chromophore moiety but does not include mTagBFP2 N-terminus; BF: bright field.

**Supplementary Figure S11.** *ACE* activity in Arabidopsis embryos extracted from fully expanded siliques. The auxin reporter is active in the tips of the cotyledons, the root, and the vasculature, but weak-to-negligible ethylene reporter activity is observed. Strong background signal in the blue channel impairs the assessment of the cytokinin reporter. Fluorescent microscopy images show stacked pictures from equatorial to cortical planes. Embryos were extracted by gently tapping on seeds embedded in a 15% glycerol solution using a microscopy glass and coverslip. ET: ethylene reporter (*EBSn:mCherry-PTS1*); CK: cytokinin reporter (*TCSn:MTS-mTagBFP\**); AUX: auxin reporter (*DR5v2:3xSV40-3xYPet*); mTagBFP*: Hybrid version of mTagBFP that contains mTagBFP2-like amino acid substitution in mTagBFP chromophore moiety but does not include mTagBFP2 N-terminus; BF: bright field.

**Supplementary Figure S12.** *ACE* expression in reproductive organs of Arabidopsis grown under long day conditions and its comparison to that of the cytokinin reporter, *TCSv2:3xmVenus-N7*^49^. The strong cytokinin activity detected in (A) sepals and (B) petal vasculature, (C) stamens, and (D) pistils by *TCSv2* is barely detected by *ACE*-encoded *TCSn*-driven cytokinin reporter due to the strong background signal. Conversely, fluorescence detected in *ACE*-encoded cytokinin and auxin reporters in (E) the funiculus (black arrow) connecting the seed and the replum was not detected in *TCSv2* cytokinin reporter, suggesting potential excitation of YPet in the 395–415 nm window and fluorescent emission in the 435–485 nm window *in planta*. Similarly, fluorescence activity patterns of ethylene reporter are indistinguishable from background fluorescence of wild type (WT). All images were acquired in the same experimental session. Images whose contrast or brightness balance was modified after image acquisition for the purpose of better displaying fluorescent patterns are labelled as “Enh. Bri/Con”. Fluorescent microscopy images are single-plane pictures. All scale bars represent 0.5 mm and are applicable to all images within the fluorescence or bright field group for a given organ. Bkgd: background fluorescence; ET: ethylene reporter (*EBSn:mCherry-PTS1*); CK: cytokinin reporter (*TCSn:MTS-mTagBFP\** (*ACE*) or *TCSv2:3xmVenus-N7* (*TCSv2*^49^); AUX: auxin reporter (*DR5v2:3xSV40-3xYPet*); mTagBFP*: Hybrid version of mTagBFP that contains mTagBFP2-like amino acid substitution in mTagBFP chromophore moiety but does not include mTagBFP2 N-terminus; BF: bright field.

**Supplementary Figure S13.** Exogenously applied hormones do not trigger prominent changes in peroxisomal localization or abundance in three-day-old etiolated Arabidopsis seedlings. (A) *ACE* reporter activity on unsupplemented or hormone-supplemented media in hooks and root tips. (B) [*35Sp:NeonGreen-PTS^PEX^*^26^] *–* [*UBQ10p:mRuby3-PTS1*]^88^ dual peroxisomal reporter activity on unsupplemented or hormone-supplemented media in hooks and root tips of seedlings grown alongside *ACE* seedlings shown in (A) under the same growth conditions. (C) Background fluorescence in wild-type (WT) seedlings grown alongside *ACE* and peroxisomal reporters. (D) Integrated single-cell relative expression of *UBQ10* in Arabidopsis roots. Modified from Root Cell Atlas (https://rootcellatlas.org/) output. All microscopy images were acquired in the same experimental session. Confocal microscopy images show stacked pictures from the equatorial plane (at the widest section of the hook or at the quiescent center of the root) to the cortical plane (surface of the organ). Scale bar applies to all images within the panel. ACC = 10 µM 1-aminocyclopropane-1-carboxylic acid; *t*-Zea = 10 µM *trans*-zeatin; IAA = 1 µM indole-3-acetic acid. ET: ethylene reporter (*EBSn:mCherry-PTS1*); CK: cytokinin reporter (*TCSn:MTS-mTagBFP\**); AUX: auxin reporter (*DR5v2:3xSV40-3xYPet*); mTagBFP*: Hybrid version of mTagBFP that contains mTagBFP2-like amino acid substitution in mTagBFP chromophore moiety but does not include mTagBFP2 N-terminus; BF: bright field.

**Supplementary Figure S14. Comparison of auxin reporters activity contained in *ACE* (*DR5v2*) vs *ACE_DD2_* (*DR5_DR5v2*) activity in three-day-old etiolated Arabidopsis seedlings.** (A) Schematic representation of the reporters, highlighting that the only difference between *ACE* and *ACEDD2* is the synthetic promoter driving auxin reporter. (B) Fluorescent images of auxin reporter activity in three-day-old etiolated seedlings upon transfer onto control (AT) or IAA-supplemented plates. All images within the panel were acquired in the same experimental session. (C) Confocal images of auxin reporter activity in seedlings presented in (B). All images within the panel were acquired in the same experimental session. Seedlings were grown for 2.5 days in the dark on horizontal control plates (AT + 6 g L^-1^ agar) and transferred onto vertical control (AT + 8 g L^-1^) or supplemented plates with 1 µM indole-3-acetic acid (IAA). They were placed vertically and incubated for 12-16 h in the dark (12 h Thunder imaging (B), 16 h confocal (C)). Pictures in B are composite images stitched together using built-in LEICA software, or manually adjusted when the software failed to accurately assemble them (*ACE_DD2_ L23*). Confocal microscopy pictures in C show stacked images from equatorial (at the quiescent center) to cortical (surface of the root) planes. AUX: auxin reporter *DR5v2: 3xSV40-3xYPet* (*ACE*) or *DR5_DR5v2: 3xSV40-3xYPet* (*ACE_DD2_*); BF: bright field.

**Supplementary Figure S15.** Comparative study of *ACE* and *ACE_DD2_* reporter expression in apical hooks and root tips of three-day-old etiolated Arabidopsis seedlings. Two independent lines per construct are displayed. All microscopy images were acquired in the same experimental session. Confocal microscopy images show stacked pictures from the equatorial to cortical planes. Seeds were germinated in the dark for three days in horizontal control (AT + 6 g L^-1^ agar) plates or supplemented with individual hormones. ACC = 0.2 µM 1-aminocyclopropane-1-carboxylic acid; *t*-Zea = 50 µM *trans*-zeatin. Higher *t*-Zea concentration than in previous assays aimed to achieve greater induction of the cytokinin reporter, but no prominent differences were observed compared to 10 µM; IAA = 50 nM indole-3-acetic acid; ET: ethylene reporter (*EBSn:mCherry-PTS1*); CK: cytokinin reporter (*TCSn:MTS-mTagBFP\**); AUX: auxin reporter *DR5v2:3xSV40-3xYPet* (*ACE*) or *DR5_DR5v2:3xSV40-3xYPet* (*ACE_DD2_*); mTagBFP*: Hybrid version of mTagBFP that contains mTagBFP2-like amino acid substitution in mTagBFP chromophore moiety but does not include mTagBFP2 N-terminus; BF: bright field.

**Supplementary Figure S16.** Schematic representation of *ACE2* variants and their nomenclature. The three-digit code after *ACE2* (*ACE2-<u>Mxy</u>*) reads as follows: (A) *M* refers to the selectable marker; (B) *x* refers to the spatial arrangement (*A*=aggregated, *S*=segregated); and (C) *y* indicates whether *mTagBFP2* is encoded by a single (*ACE2-Mx1*) or triple tandem (*ACE2-Mx3*) fluorescent protein. LB: left T-DNA border; RB: right T-DNA border; *PTS1: Peroxisomal Targeting Signal 1*. *MTS: Mitochondrial Targeting Signal*; *NLS: Nuclear Localization Signal*. *N7v1-N7v3* refers to three sequence-divergent coding DNA sequences for *N7 NLS*^89^. *CVMVp*: promoter from Cassava vein mosaic virus^63,65^; *tHSP18.2*: *HEAT SHOCK PROTEIN 18.2* terminator^116^.

**Supplementary Figure S17.** Characterization of the *ACE2* reporter variants in Arabidopsis root tips. (A) Half-root Z-stack images of two independent insertional lines of *ACE2* reporter in its two spatial arrangements. T1 seedlings show strong activity upon exogenous hormone supplementation and improved delivery to the targeted organelles relative to *ACE*. (B) Single-plane images of the whole primary root and root tip of two lines displayed in (C). Details of the root tip were acquired with increased optical magnification of the regions marked with a white box. (D) Half-root Z-stack images of two independent insertional events of *ACE2-3Sy* show that triple mTagBFP2 does not improve the detection of basal reporter activity compared to the single copy version under control conditions in T1 seedlings. All microscopy images were acquired in the same experimental session. Seeds were germinated for three days in the dark on AT plates (6 g L^-1^ agar) supplemented with 10 µg mL^-1^ phosphinothricin and 300 µg mL^-1^ timentin and grown for four additional days under continuous light. Resistant plants (with healthy cotyledons as opposed to those that bleached) were transferred to sugar-less AT (8 g L^-1^ agar) plates supplemented with 300 µg mL^-1^ timentin as control, or adding all three hormones (10 µM ACC + 30 µM *t*-Zea + 10 µM NAA) replacing IAA with the more photostable synthetic auxin, NAA, for 16 h prior to image acquisition. ET: ethylene reporter (*EBSn:MTS-mCherry*); CK: cytokinin reporter (*TCSv2:mTagBFP2-N7*); AUX: auxin reporter (*DR5v2:YPet-PTS1*); BF: bright field.

**Supplementary Figure S18.** Characterization of the second-generation *ACE* reporter, *ACE2-3A1*, in stable Arabidopsis transgenic lines. (A) *ACE2-3A1* construct structure. Fragments not at scale. (B) Basal reporter activity and hormone inducibility of *ACE2-3A1* compared to the first-generation ACE reporter in hooks (top) and roots (bottom) of Arabidopsis. Three-day-old etiolated seedlings were imaged under endogenous hormone concentrations (AT) and following supplementation with ACC, trans-zeatin, or IAA to selectively activate ethylene, cytokinin, and auxin signaling pathways, respectively. Comparison of reporter generations showed consistent patterns of activity and induction, with T5 generation being presented here. Circular schematic shows that the expected subcellular localization of the fluorescent proteins is nuclear for all three reporters (Mt: mitochondria, Px: peroxisomes, Nu: nucleus). All microscopy images were acquired using identical acquisition settings for direct comparison between reporter generations. Confocal microscopy images show stacked pictures from equatorial to cortical planes (half-root Z-stacks). L3.5 and L4.3 refer to two independent single-insertion transgenic lines of *ACE2-3A1* reporter. Scale bar applies to all images within the panel. Images were digitally enhanced to facilitate detection of low-expressing cells and the same settings were applied to all images within a channel across treatments. LB: left T-DNA border; RB: right T-DNA border; NLS: Nuclear Localization Signal (N7); ACC = 10 µM 1-aminocyclopropane-1-carboxylic acid; *t*-Zea = 10 µM *trans*-zeatin; IAA = 1 µM indole-3-acetic acid; ET: ethylene reporter (*EBSn*); CK: cytokinin reporter (*TCSv2*); AUX: auxin reporter (*DR5v2*); BF: bright field.

**Supplementary Figure S19.** Logic to build a nonuple reporter to monitor phytohormone concentrations *in vivo* (*aka*, the hormometer). Combining hormone-specific promoters responsive to all nine major non-peptidic phytohormone classes does not fully resolve the constraints of simultaneous *in vivo* detection imposed by spectral limitations. (A) Three fluorescence windows are robustly separable in plants: blue (e.g., mTagBFP2), green–yellow (e.g., YPet), and red (e.g., mCherry). This limitation restricts conventional multiplexing strategies. However, spatial segregation of fluorescent proteins through subcellular targeting offers an orthogonal dimension for multiplexing (B). By combining three spectrally distinct fluorophores (mTagBFP2, YPet, and mCherry) with targeting to the nucleus, peroxisomes, and mitochondria, and driving each with distinct hormone-responsive promoters, it is theoretically feasible to generate a nonuple synthetic reporter capable of monitoring nine hormone classes in parallel.

**Supplementary Figure S20. Cytokinin-inducible *TCSn* promoter subcloning strategy.** (A) GoldenBraid (GB) domestication of the *TCSn:35SminPro0* promoter. Oligos F1 and R1 (Supplementary Table S1A) were designed to amplify the *TCSn:35SminPro0* sequence from *pUC18_TCSn:35SminPro0:LUC:NOSter*^48^ and introduce the DNA elements needed for its subcloning into *pUPD2* entry vector (with the GB code for a proximo-distal promoter (A1-A2) shown in purple, the *pUPD2* cloning overhang marked in dark blue, and BsmBI recognition sites indicated in cyan) in a similar fashion as previously described^51^. The amplification product included the 5’ end of the *35SminPro0* sequence (dark grey) due to the difficulty of designing a reverse oligo over the repetitive *TCSn* promoter (yellow). (B) PCR and GB cloning of the *TCSn:35SminPro0* promoter. The amplification of the *TCSn:35SminPro0* promoter was done using iPROOF High-Fidelity DNA polymerase Kit (Bio-Rad Laboratories, 172-5302) with the following conditions: 94^ᴼ^C – 2min, (94^ᴼ^C – 30sec, 56^ᴼ^C – 30sec, 72^ᴼ^C – 2min)x35, 72^ᴼ^C – 10min, 10^ᴼ^C – ∞min. For subcloning the amplified promoter fragment into the *pUPD2* entry vector, previously described conditions^51^ were used. Briefly: (37 ^ᴼ^C – 4 min; 16 ^ᴼ^C – 5 min)x25, 55 ^ᴼ^C – 20 min, 80 ^ᴼ^C – 20 min using BsmBI (NEB CutSmart R05580L) restriction enzyme and T4 DNA ligase (NEB M0202L). (C,D). *minPro*-less oligo design and the resulting PCR products. Three additional oligos (F2, R2 and R3, Supplementary Table S1A) were created for the re-amplification of the *pUPD2_TCSn:35SminPro0* fragment and subsequent recloning of it without the *35SminPro0*. The R3 oligo was designed to include the AcuI restriction site (red) 14 bp downstream of the 3’-end of *TCSn* sequence. Together with F1, the two oligos amplified PCR product 1 (336 bp long). Oligos F2 and R2 are overlapping oligos designed to complement the sticky end of PCR product 1 while adding the *pUPD2* cloning overhang to complete a proximo-distal promoter DNA element (A1-A2); they amplified PCR product 2 (45 bp long). A random sequence of 14 bp was included between the AcuI recognition site and the 2 bp long *TCSn* sequence (sticky end). Same PCR conditions as in panel B were used. (E) AcuI digestion and ligation. This step was designed to generate a *minPro*-less *TCSn* sequence flanked by the *pUPD2* cloning overhangs. For that, the AcuI restriction enzyme cut 16 bp away from the recognition site and left 2 bp sticky ends in both PCR products, 1 and 2. When ligated by T4 DNA ligase, PCR product 2 filled the void made by AcuI in PCR product 1, regenerating the *TCSn* promoter while removing the *35SminPro0* sequence. The cloning conditions for this reaction were (37 ^ᴼ^C – 4 min; 16 ^ᴼ^C – 5 min)x25, 37 ^ᴼ^C – 20 min, 65 ^ᴼ^C – 20 min. **F.** *minPro*-less *TCSn* GB cloning into *pUPD2* entry clone. Following a similar GB assembly strategy as mentioned in panel B, the *minPro*-less *pUPD2_TCSn* clone was generated (Supplementary Table S1A).

**Supplementary Figure S21. Oligo design strategies for auxin *DR5*-related and *fullProCVMV* promoters. (**A) Oligo design for the individual auxin-inducible *DR5_×10_* and *DR5v2_×10_* proximo-distal (A1-A2) promoters. F1 and F2 oligos (Supplementary Table S1A) were designed over the 5’-multiple cloning site (MSC) region and the first *DR5* or *DR5v2* repeat, respectively, to provide sequence specificity while ensuring primer annealing to the first repeat of these repetitive promoters. Additionally, the oligos are flanked by a 5’-A1 GB code (purple), a *pUPD2* cloning overhang (dark blue), and a BsmBI recognition site (cyan) for their subcloning into *pUPD2* entry vector. The reverse oligo (R1) was made complementary to the common 3’-MCS region and was flanked by the same cloning elements as F1 and F2 but adding the 3’-A2 GB code instead. (B) Oligo design for the individual auxin-inducible *DR5_×10_:DR5v2_×10_* proximo-distal (A1-A2) promoter. Two additional oligos were designed (F3 and R2, Supplementary Table S1A) to amplify DR5_×10_ and DR5v2_×10_ as distal (A1) and proximal (A2) promoters, respectively, and allow their assembly during their subcloning into *pUPD2* entry vector to create a final combined *DR5_×10_:DR5v2_×10_* proximo-distal (A1-A2) promoter. For that, the 5’-A1 GB code in F2 oligo was changed to the 5’-A2 GB code (highlighted in green) to create F3. Likewise, the 3’-A2 GB code in R1 oligo was switched to the 3’-A1 GB code (highlighted in green) to create R2. The template used for amplifying *DR5_×10_*, *DR5v2_×10_*, and *DR5_×10_:DR5v2_×10_* proximo-distal promoters was *pGREEN II_DR5×10-dTomato / DR5v2-3nGFP*^47^. The PCR products obtained with each oligo pair combination are displayed on the right side of the figure. (C) Oligo design for subcloning the *Cassava vein mosaic virus* (*CVMV*) promoter. The full-length *CVMV* promoter was split into two independent GBlocks to pass the quality thresholds set by the vendor (Integrative DNA Technologies) for its commercial synthesis. For that, a *CVMV* 480 bp fragment was included in the GBlock_A, and the remaining *CVMV* 318 bp fragment was synthesized as the GBlock_B. Four oligos (F1, F2, R1, and R2, Supplementary Table S1B) were designed to amplify both fragments and allow their scar-less assembly into the *pUPD2* entry clone. The oligos F1 and R1 harbor the GB codes (A1 and B1, respectively) shown in purple, the *pUPD2* cloning overhangs displayed in dark blue, and the BsmBI recognition sites marked in cyan, enabling the fragment subcloning into the entry vector. Oligos F2 and R2 are flanked by the BsmBI recognition sites (cyan) to generate compatible 4 bp sticky ends. During the regular GB assembly conditions^98^, the T4 DNA ligase will combine the two PCR fragments with the vector, thus regenerating the full-length of *fullProCVMV* promoter. PCR conditions employed herein are the same as described in Supplementary Figure S20.

## Abbreviations

SCF^TIR1/AFB^: SKP1–CULLIN–F-BOX complex with TRANSPORT INHIBITOR RESPONSE1/AUXIN SIGNALING F-BOX proteins
AUX/IAA: AUXIN/INDOLE-3-ACETIC ACID PROTEINS
ARF: AUXIN RESPONSE FACTOR
AHK: ARABIDOPSIS HISTIDINE KINASE
AHP: ARABIDOPSIS HISTIDINE-CONTAINING PHOSPHOTRANSFER PROTEIN
ARR: ARABIDOPSIS RESPONSE REGULATOR
EIN2/EIN3: ETHYLENE INSENSITIVE2/3
TAA1: TRYPTOPHAN AMINOTRANSFERASE OF ARABIDOPSIS1
AUX1: AUXIN RESISTANT1
LRC: lateral Root Cap
EPI: epidermis
COR: cortex
END: endodermis
PER: pericycle
STE: stele
QC: quiescent center
CLM: columella
SCN: stem cell niche
DZ: differentiation Zone
EZ: elongation zone
TZ: transition zone
RAM: root apical meristem
ARG, CRG, ERG: auxin-, cytokinin-, ethylene-regulated genes

## Acknowledgements

The work in the Alonso-Stepanova lab is supported by National Science Foundation grants 2503590, 1650139, and 1444561 to JMA and ANS, and by grant 1750006 to ANS, and Research Capacity Fund (HATCH) project awards 7005468 and 7005482 from the U.S. Department of Agriculture’s National Institute of Food and Agriculture to JMA and ANS, respectively. KV is a recipient of a Graduate Research Fellowship Program award 202335657. AEY is a recipient of the Genetics and Genomics Scholars fellowship from NC State University and of Molecular Biotechnology Training Program grant from the National Institutes of Health. AGD is the recipient of Coordenação de Aperfeiçoamento de Pessoal de Nível Superior - Brasil (CAPES) – Finance Code 001 / CAPES-PrInt Program fellowship and DSM research fellow of CNPq (309743/2021-4). S.B. was supported by EU-Marie Skłodowska-Curie grant agreement No. 101007738-EVOfruland.

We thank Dr. Bonnie Bartel (Rice University, USA) for sharing the [*35S:NeonGreen-PTS^PEX^*^26^] – [*UBQ10:mRuby3-PTS1*] peroxisomal dual marker line, Dr. Joseph Kieber (University of North Carolina at Chapel Hill, USA) for providing the *ahk3-3 cre1-12* mutant, Dr. Maya Bar (Volcani Center, Israel) for the Arabidopsis *TCSv2* reporter line, Dr. Jan Petrasek for valuable discussions and input during the development of this work, and undergraduates Kirby Morris and Damla Ayrilmaz for technical assistance. We are also grateful to Drs. Miguel Flores Vergara, Robert Franks, Imara Perera, and Marcela Rojas-Pierce for sharing reagents and equipment. We would also like to thank NC State University’s Genomic Sciences Laboratory for Sanger sequencing services and the Cellular and Molecular Imaging Facility for providing user training and access to their equipment. We thank members of the Alonso-Stepanova laboratory and the INTRINSyC Plant Biology community at North Carolina State University for helpful discussions and feedback on the manuscript. Large language models were used to assist with language editing and to improve clarity. The authors thoroughly reviewed and edited the manuscript and take full responsibility for its content.

## Author information

### Contributions

T.A.-I., A.N.S. and J.M.A. conceived the project. A.N.S., J.M.A., M.F. and J.P.F.-M. designed the research. J.P.F.-M. led the generation and validation of GoldenBraid-compatible library with the contribution of C.X. and M.F., assembled *ACE*, and transformed it into Arabidopsis. M.F. led the isolation, introgression, and characterization of *ACE* lines in WT and mutant backgrounds with the contribution of A.B. and A.G.D.; M.F. led the assembly of *ACE2* with the contribution of A.G.D., A.N., S.B., H.H., J.J., K.V. and K.M.; M.F and A.G.D transformed *ACE2* in Arabidopsis; M.F. and J.S.T. characterized *ACE2* activity. D.S.M., M.M.K., T.J.A.-I., A.N.S. and J.M.A. supervised the research. All authors discussed the results and contributed to the final manuscript.

## Ethics declarations

### Code availability

All scripts were deposited at Zenodo:

- Arabidopsis codon-optimized CDSs, DOI: 10.5281/zenodo.21797666
- *35Sp*-derived minimal promoters (*minPro*), DOI: 10.5281/zenodo.21797744
- *35Ster*-derived terminators (*ter*), DOI: 10.5281/zenodo.21797824

### Data availability

DNA clones will be deposited in Addgene and the seeds of the *ACE/ACE2* sensors in the ABRC stock center

