## Supplemental Figures for "An Integrated Spatially Resolved Mechanistic Model of Hierarchical Auxin–Cytokinin–Ethylene Crosstalk Underlying Root Growth Inhibition in Arabidopsis"

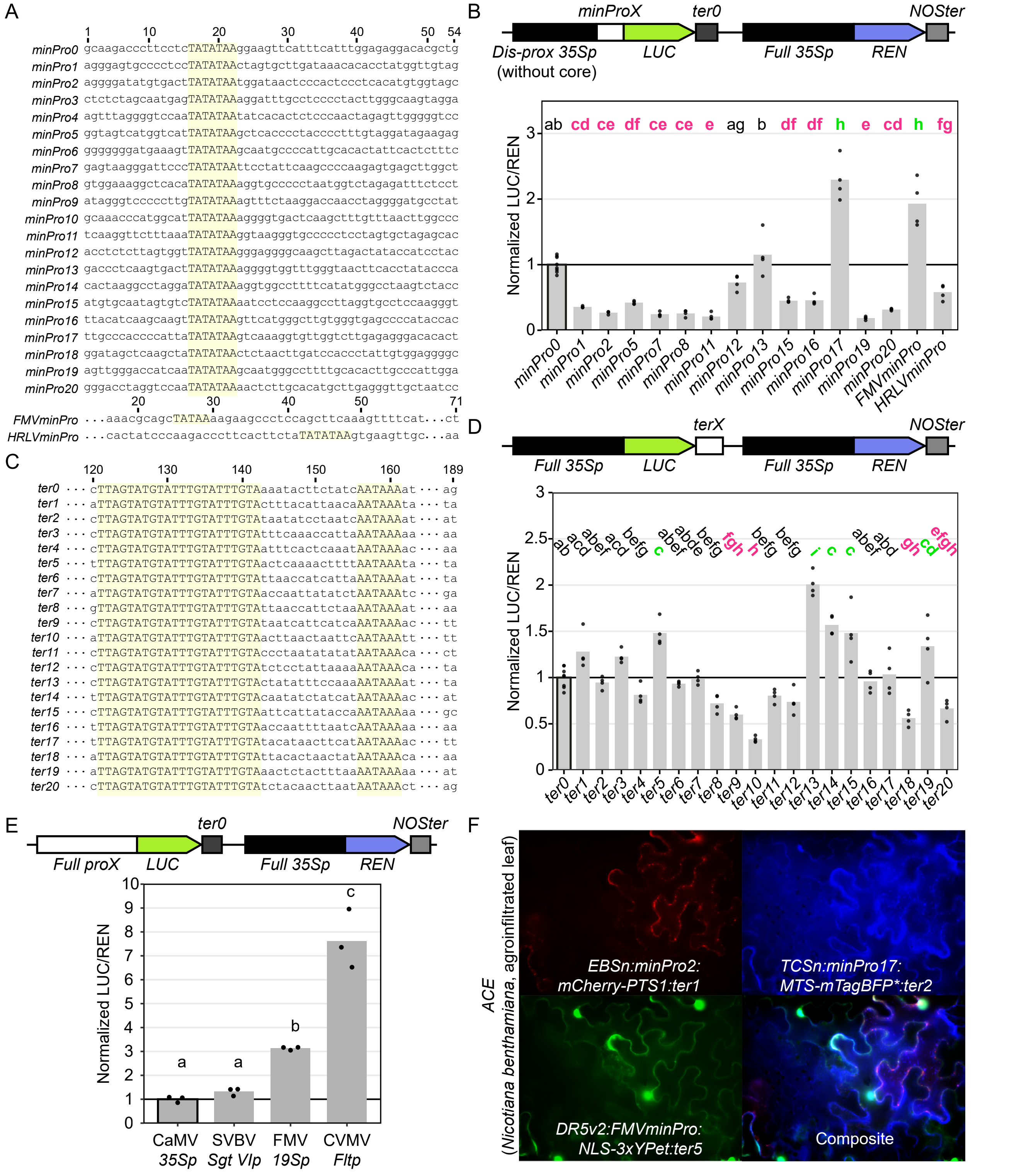


**Supplementary Figure S1. Collection of minimal promoters (*minPro*) and terminators (*ter*) assessed for transcriptional activity in transient expression assays in agroinfiltrated *Nicotiana benthamiana* leaves.**

(A) DNA sequence comparison of 22 minimal promoters (twenty of which were derived from CaMV *35Sp* plus those derived from FMV DxS strain subgenomic transcript VI (19S) or HRLV full-length transcript) relative to CaMV *35SminPro0* (*minPro0*) visualized using MEGAX[117] software without sequence alignment. The position of the *TATA* box (highlighted in yellow) is preserved in all *35Sp*-derived promoters.

(B) Construct structures and normalized activities of fifteen minimal promoters relative to *minPro0* (black-edged vertical bar; horizontal black line at y-axis=1) were assessed by transient assays in *Nicotiana benthamiana* leaves agroinfiltrated with [*DisProx35Sp:minProX:LUCIFERASE:ter0*] – [*35Sp:RENILLA:NOSter*]. Vertical bars represent the average LUC/REN ratios, and data points indicate individual pseudoreplicates (n=4). *DisProx35Sp* refers to CaMV *minPro0*-less *35Sp*. For visual reference, significant differences between a given *minProX* and *minPro0* are highlighted in bold letters, with magenta and green indicating lower or higher activity, respectively.

(C) DNA sequence comparison of the twenty CaMV *35Ster*-derived terminators compared to *35Ster0* (*ter0*) visualized using MEGAX[117] without sequence alignment. The polyadenylation signals are highlighted in yellow and their position is preserved in all terminators.

(D) Construct structures and normalized activities of twenty terminators relative to *ter0* (black-edged vertical bar; horizontal black line at y-axis=1) assessed by transient assays in Nicotiana benthamiana leaves agroinfiltrated with [*35Sp:LUCIFERASE:terX*] – [*35Sp:RENILLA:NOSter*]. Vertical bars represent the average LUC/REN ratios and data points indicate individual pseudoreplicates (n=4). For visual reference, significant differences between a given *terX* and *ter0* are highlighted in bold letters, with magenta and green indicating lower or higher activity, respectively.

(E) Construct structures and normalized activities of three viral-sourced promoters relative to CaMV *35Sp* (black-edged vertical bar; horizontal black line at y-axis=1) were assessed by transient assays in *Nicotiana benthamiana* leaves agroinfiltrated with [*Full_proX:LUCIFERASE:ter0*] – [*35Sp:RENILLA:NOSter*]. *Full_proX*: full promoters from the Strawberry vein banding virus (SVBV; 988 bp) Sgt VI[61,62], FMV M3 strain (416 bp) 19S (Sgt VI)[63,64], or Cassava vein mosaic virus (CVMV; 2895 bp) Flt[63,65] (Sgt: subgenomic transcript; Flt: full-length transcript; VI: sixth open reading frame). Vertical bars represent the average LUC/REN ratio, and data points indicate individual pseudoreplicates (n=3) representing technical variability. Different letters denote statistically significant differences between means (one-way ANOVA, α=0.05).

(F) Transient expression of *ACE* reporter in N. benthamiana leaves upon hormone infiltration.

ACC = 10 μM 1-aminocyclopropane-1-carboxylic acid, *t*-Zea = 10 μM *trans*-zeatin, IAA = 0.1 μM indole-3-acetic acid. *PTS1: Peroxisomal Targeting Signal 1* (N-KSRM-C). *MTS: Mitochondrial Targeting Signal* (MitCox4); *NLS: Nuclear Localization Signal* (3xSV40). mTagBFP*: Hybrid version of mTagBFP that contains mTagBFP2-like amino acid substitution in mTagBFP chromophore moiety but does not include the mTagBFP2 N-terminus.

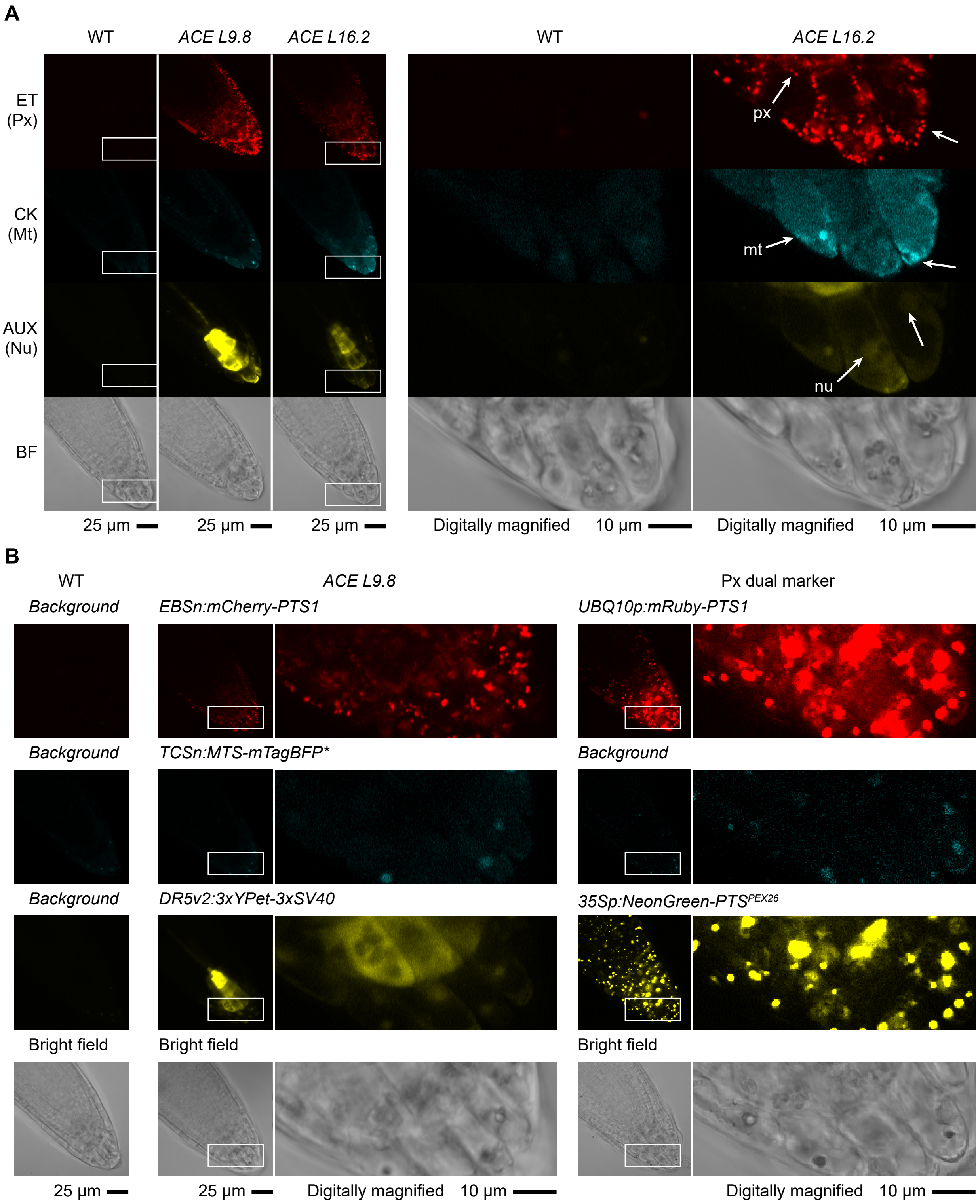


**Supplementary Figure S2. Subcellular localization of the *ACE* reporter in Arabidopsis root tip cells.**
(A) Two independent insertion lines of the *ACE* reporter (*L9.8* and *L16.2*) reveal subcellular localization of mCherry and mTagBFP* consistent with peroxisomes (Px) and mitochondria (Mt), respectively, whereas YPet signal is not confined to the nucleus (Nu).
(B) Comparison of mCherry localization in the *ACE* reporter with the previously described dual peroxisomal marker [*35Sp:NeonGreen-PTS$^{PEX26}$*] - [*UBQ10p:mRuby3-PTS1*][88] imaged in separate roots supports peroxisomal targeting of mCherry.
White boxes represent the area of the root that is magnified within the same panel for a given genotype and reporter. Three-day-old etiolated seedlings were germinated on unsupplemented media. All microscopy images were acquired in the same experimental session. Confocal microscopy images show stacked pictures from the equatorial plane at the quiescent center to the cortical plane of the lateral root cap. Scale bar applies to all images within the panel. ET: ethylene reporter (*EBSn:mCherry-PTS1*); CK: cytokinin reporter (*TCSn:MTS-mTagBFP**); AUX: auxin reporter (*DR5v2:3xSV40-3xYPet*); mTagBFP*: Hybrid version of mTagBFP that contains mTagBFP2-like amino acid substitution in mTagBFP chromophore moiety but does not include mTagBFP2 N-terminus; BF: bright field; WT: wild-type.

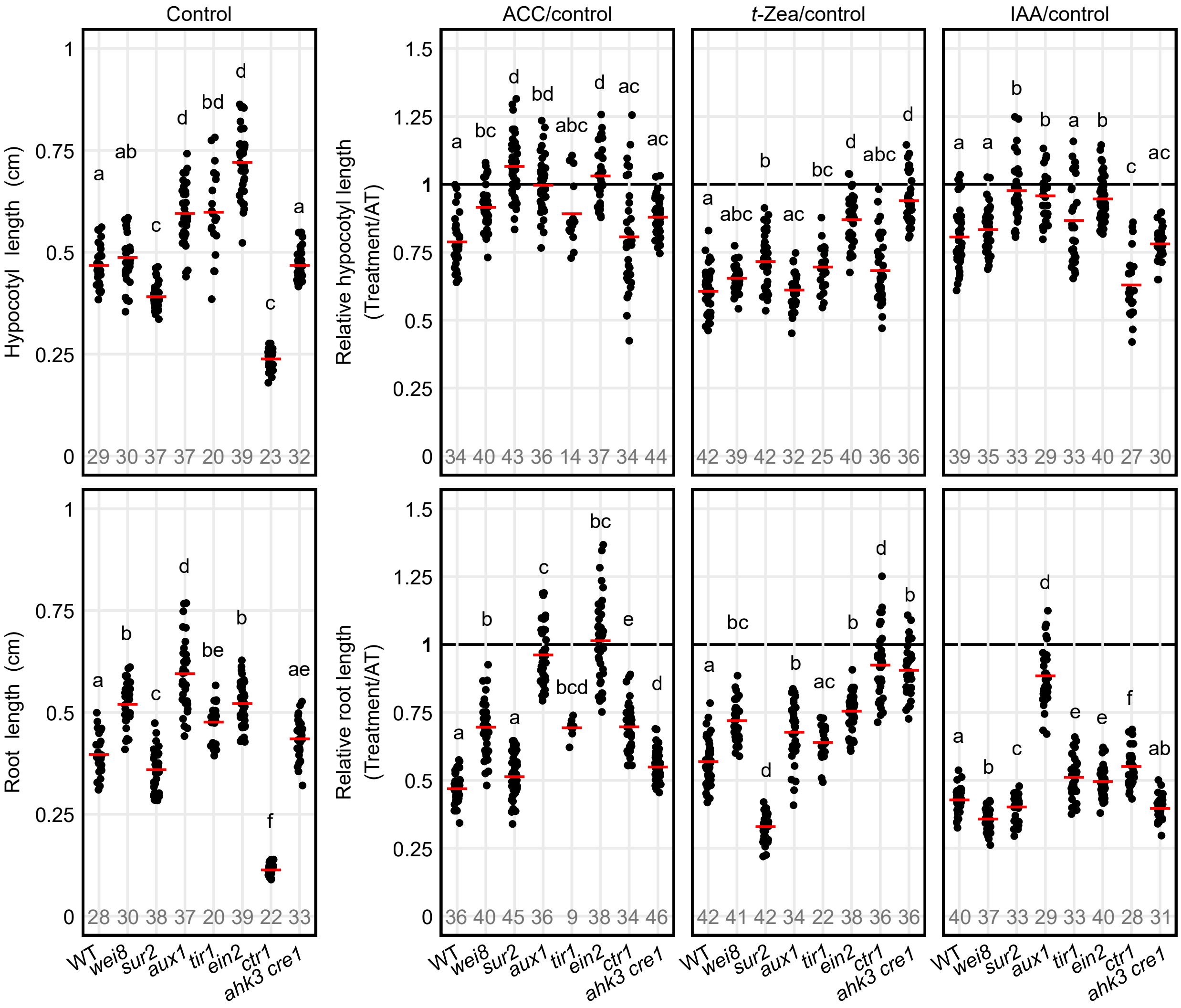


**Supplementary Figure S3. Quantitative analysis of relative organ sizes in three-day-old etiolated wild-type (WT) and mutant seedlings of Arabidopsis defective in ethylene, cytokinin, or auxin biosynthesis, perception, signaling, or transport in response to exogenous hormone treatments.**

All genotype x treatment combinations were grown in the same experimental session. Black dots represent independent organ measurements of individual seedlings and horizontal red bars across them represent the mean values. Numbers at the bottom of the plot (in grey) represent the number of biological replicates (n) for a given organ and treatment. Different letters indicate statistically significant differences among genotypes for a given organ within each treatment. Multiple comparison analyses were performed as described in the methods section under Image acquisition and data analysis.

ACC = 0.2 μM 1-aminocyclopropane-1-carboxylic acid; *t*-Zea = 10 μM *trans*-zeatin; IAA = 50 nM indole-3-acetic acid.

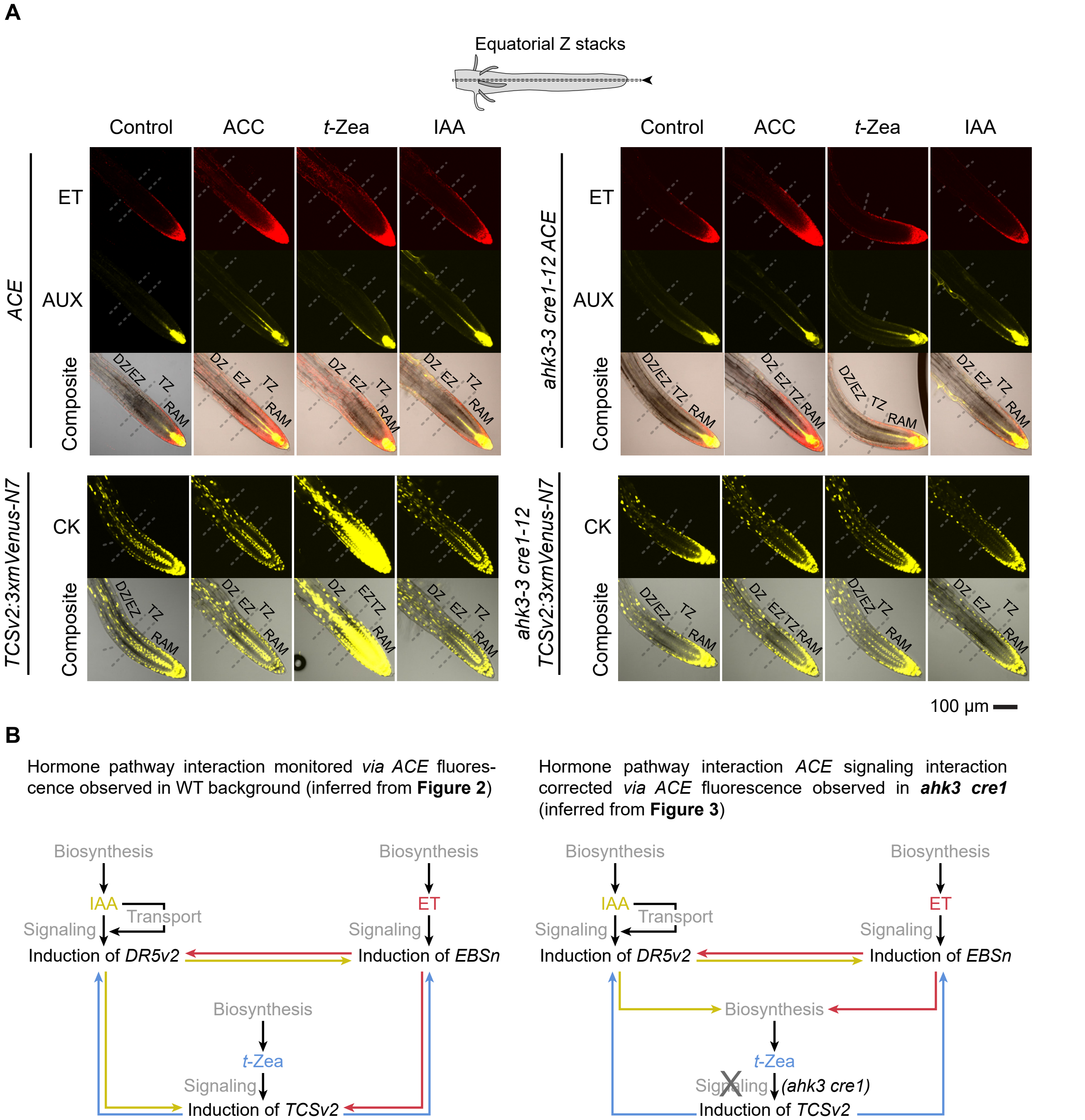


**Supplementary Figure S4. Detailed view of AHK3- and CRE1-dependent responses to exogenous hormone treatments in signaling pathway crosstalk in Arabidopsis roots.**

(A) Equatorial Z-stack of *ACE* and *TCSv2* activities in longitudinal sections of the same Arabidopsis roots displayed in Figure 3A. Scale bar applies to all images within the panel. ACC = 0.2 µM 1-aminocyclopropane-1-carboxylic acid; *t*-Zea = 10 µM *trans*-zeatin; IAA = 50 nM indole-3-acetic acid; ET: ethylene reporter (*EBSn:mCherry-PTS1*); CK: cytokinin reporter (*TCSv2:3xmVenus-N7*)49; AUX: auxin reporter (*DR5v2:3xSV40-3xYPet*); BF: bright field; DZ: differentiation zone; EZ: elongation zone; TZ: transition zone; RAM: root apical meristem.

(B) Updated Arabidopsis hormone signaling pathway interaction model based on *ACE* and *TCSv2* activities and phenotypic responses of *ahk3 cre1* roots to exogenous application of hormones. Based on *TCSv2* and *ACE* reporters, IAA and ACC induce *TCSv2* activity. The *ahk3 cre1* mutation prominently reduces ACC- and IAA-triggered upregulation of *TCSv2*, suggesting that ethylene and auxin work upstream or at the level of cytokinin receptors.

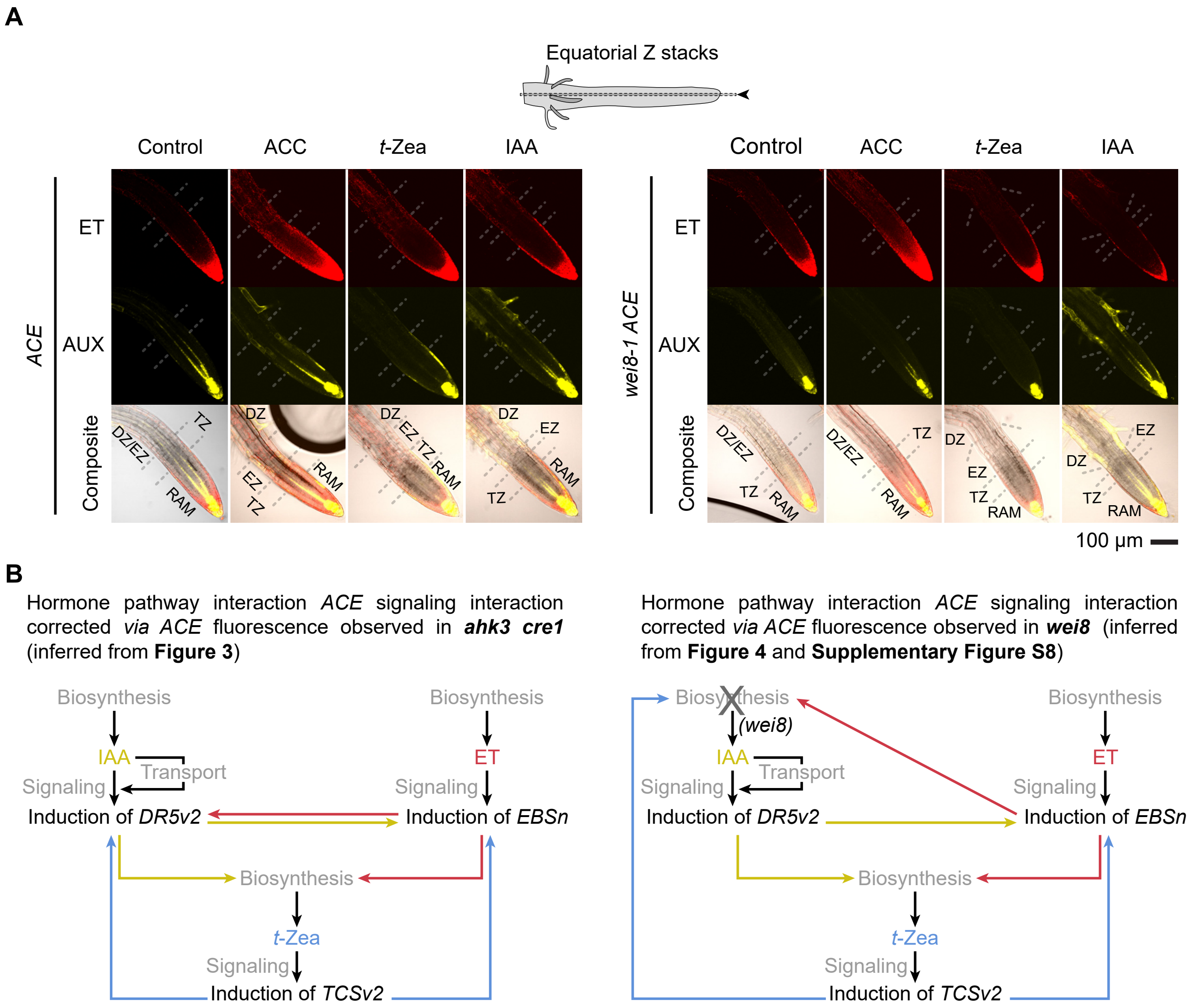


**Supplementary Figure S5. Detailed view of TAA1-dependent responses to exogenous hormone treatments in signaling pathway crosstalk in Arabidopsis roots.**

(A) Equatorial Z-stack of *ACE* activity in longitudinal sections of the same Arabidopsis roots displayed in Figure 4A. Scale bar applies to all images within the panel. ACC = 0.2 µM 1-aminocyclopropane-1-carboxylic acid; *t*-Zea = 10 µM *trans*-zeatin; IAA = 50 nM indole-3-acetic acid; ET: ethylene reporter (*EBSn:mCherry-PTS1*); AUX: auxin reporter (*DR5v2:3xSV40-3xYPet*); BF: bright field; DZ: differentiation zone; EZ: elongation zone; TZ: transition zone; RAM: root apical meristem. Root zone boundaries are approximate.

(B) Updated Arabidopsis hormone signaling pathway interaction model based on *ACE* activity and phenotypic responses of *wei8* roots to exogenous application of hormones. *wei8* mutant shows normal ethylene signaling, but its ACC-induced auxin activity is suppressed. Given that *wei8* shows mild insensitivity to ACC (Figure 4B), the results suggest that ethylene-decreased root length involves TAA1-mediated increase in auxin biosynthesis in response to ACC (supported by Supplementary Figure S8). Furthermore, *wei8* also suppresses *t*-Zea-mediated increase in auxin signaling, consistent with exogenously supplied cytokinin leading to a boost in TAA1-mediated auxin biosynthesis (supported by Supplementary Figure S8).

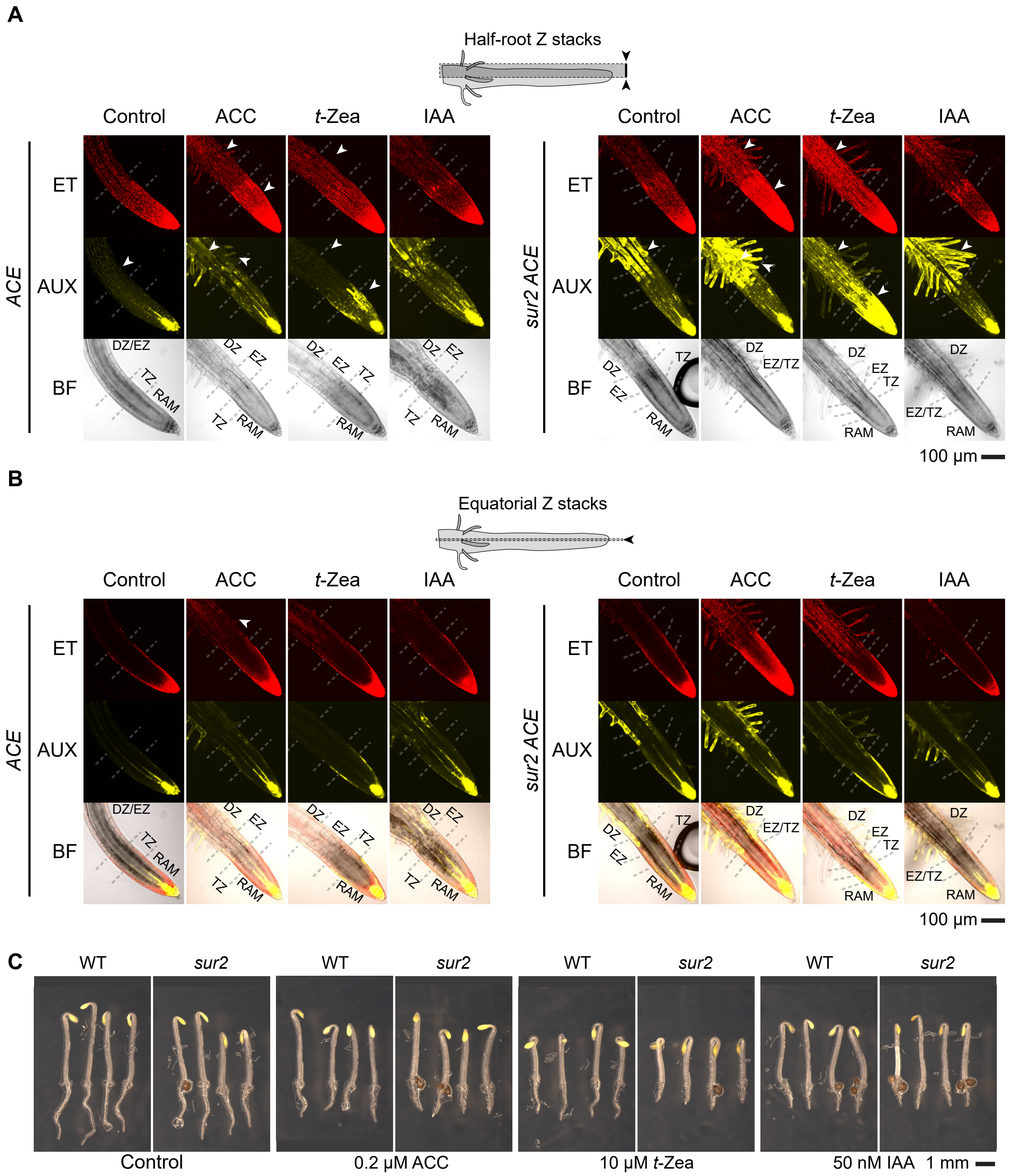


**Supplementary Figure S6. High endogenous concentration of auxin in *sur2* sensitizes the Arabidopsis root to the perception of ethylene and cytokinin, but it does not translate into prominently higher basal *EBSn* activity in the roots of three-day-old etiolated seedlings.**

(A) Half-root Z-stacks of an auxin overproducing mutant *sur2* show stronger basal activity of auxin and ethylene reporters. Moreover, high auxin in *sur2* increases the sensitivity of the root to all exogenously applied hormones tested. Arrowheads point at root sections where reporter expression differs between treatments or genotypes.

(B) Equatorial Z-stack of *ACE* activity in longitudinal sections of the same Arabidopsis roots displayed in Figure 6A. Scale bar applies to all images within the panel.

(C) *sur2* roots show stronger sensitivity to exogenous cytokinin and mildly, yet significantly stronger sensitivity to exogenous IAA. Despite the ethylene reporter in *sur2* roots being more sensitive than WT to exogenous ACC, *sur2* relative root length is similar to that of wild type (WT). Quantitative comparison and statistical assessment of hormone-specific inhibitory effect on growth between WT and mutants are available in Supplementary Figure S3.

All microscopy images were acquired in the same experimental session. Half root Z-stack (A) confocal microscopy images show stacked pictures from the equatorial plane at the quiescent center to the cortical plane of the lateral root cap, while those labelled as “Equatorial” (B) show stacked pictures immediately surrounding the quiescent center. Scale bar applies to all images within the panel. ACC = 0.2 µM 1-aminocyclopropane-1-carboxylic acid; *t*-Zea = 10 µM *trans*-zeatin; IAA = 50 nM indole-3-acetic acid. ET: ethylene reporter (*EBSn:mCherry-PTS1*); AUX: auxin reporter (*DR5v2:3xSV40-3xYPet*); BF: Bright field. DZ: differentiation zone; EZ: elongation zone; TZ: transition zone; RAM: root apical meristem. Root zone boundaries are approximate.

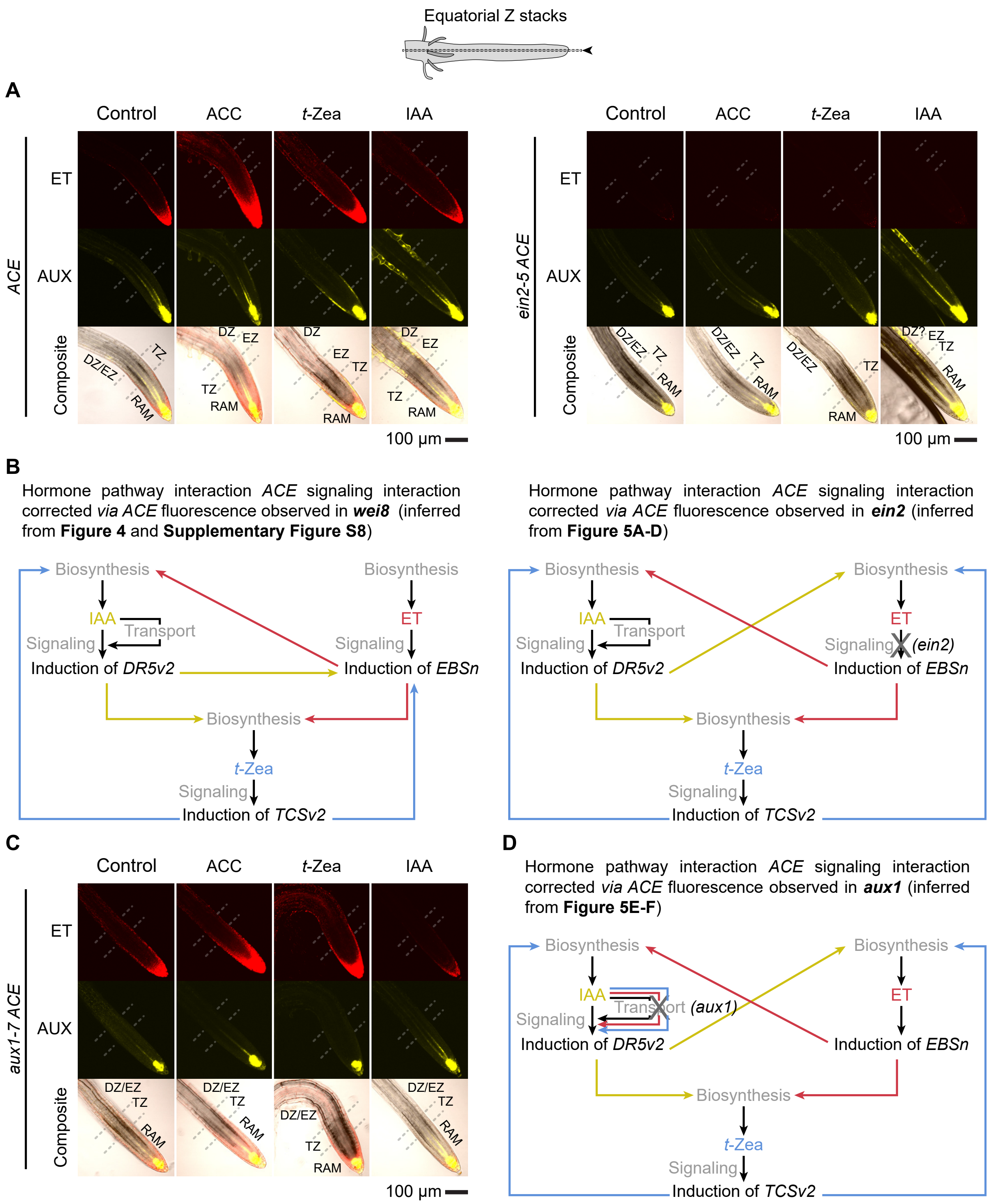


**Supplementary Figure S7. Detailed view of EIN2- and AUX1-dependent responses to exogenous hormone treatments in the signaling pathway crosstalk in Arabidopsis roots.**

(A) Equatorial Z-stack of *ACE* activity in longitudinal sections of the same Arabidopsis roots displayed in Figure 5A and C.

(B) Updated Arabidopsis hormone signaling interaction model based on *ACE* activity and phenotypic responses of *ein2 ACE* roots to exogenous application of hormones. IAA-dependent induction of *EBSn* activity is suppressed in *ein2*, suggesting that auxin signaling can also operate upstream of ethylene inducing its signaling activity in an EIN2-dependent manner, rather than supporting the scenario where a subset of auxin-inducible genes trans-activate ethylene-inducible genes. Furthermore, cytokinin-dependent induction of ethylene and auxin signaling is also suppressed in *ein2*, suggesting that in WT roots, cytokinin stimulates ethylene biosynthesis or ethylene perception, hence leading to the increase of *EBSn* activity and a boost of ethylene-dependent auxin biosynthesis, rather than cytokinin directly stimulating auxin biosynthesis.

(C) Equatorial Z-stack of *ACE* activity in longitudinal sections of the same Arabidopsis roots displayed in Figure 5E.

(D) Updated Arabidopsis hormone signaling pathway interaction model based on *ACE* activity and phenotypic responses of *aux1* roots to exogenous application of hormones. Despite having normal ethylene signaling and fully functional auxin biosynthesis machinery, *aux1* roots fail to increase auxin signaling upon ACC or *t*-Zea supplementation, suggesting that ethylene- or cytokinin-induced local auxin biosynthesis requires normal auxin import from source cells (where TAA1 reporter accumulates; Supplementary Figure S8) to sink cells (showing *DR5v2* maxima in WT plants). However, a group of cells surrounding the quiescent center show increased auxin activity upon exposure to exogenous hormones, suggesting that increased auxin might be channeled towards the root tip through other transportation paths.

Scale bar applies to all images within the panel. ACC = 0.2 μM 1-aminocyclopropane-1-carboxylic acid; *t*-Zea = 10 μM *trans*-zeatin; IAA = 50 nM indole-3-acetic acid; ET: ethylene reporter (*EBSn:mCherry-PTS1*); AUX: auxin reporter (*DR5v2:3xSV40-3xYPet*); BF: bright field; DZ: differentiation zone; EZ: elongation zone; TZ: transition zone; RAM: root apical meristem. Root zone boundaries are approximate.

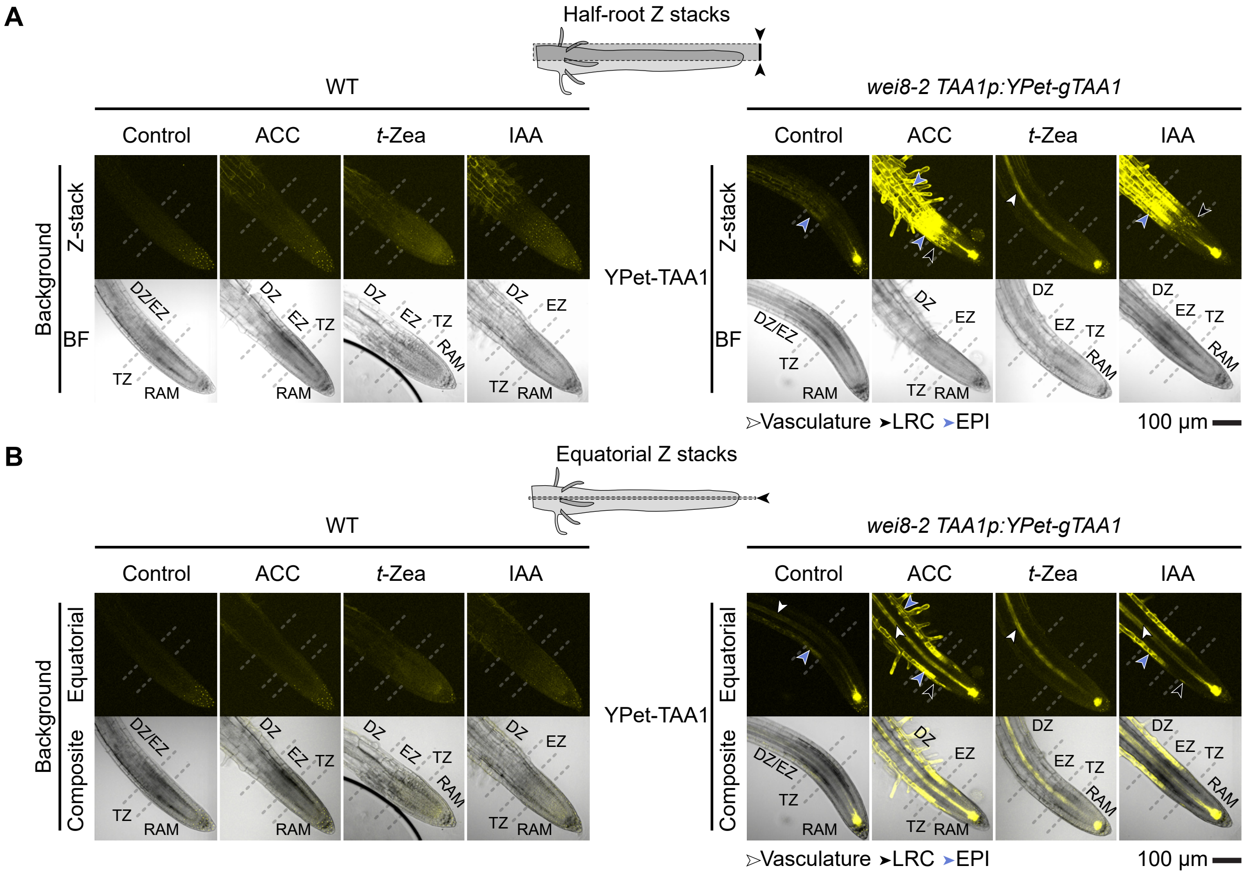


**Supplementary Figure S8. TAA1 accumulation pattern in Arabidopsis roots responds differently to the application of distinct exogenous hormones in *wei8-2 TAA1p:YPet-gTAA1*.**

(A) Half-root Z-stacks showing that TAA1 accumulation in roots is consistently induced by all hormones relative to untreated control.

(B) Equatorial section of the roots displayed in panel A. While *t*-Zea induces TAA1 accumulation mainly in the vasculature of the primary root, ACC and IAA induce TAA1 accumulation also in the EPI and LRC, with ACC triggering stronger TAA1 accumulation in the TZ and EZ than IAA. Arrowheads point at specific cell types where reporter expression differs between treatments.

All microscopy images were acquired in the same experimental session. Half root Z-stack (A) confocal microscopy images show stacked pictures from the equatorial plane at the quiescent center to the cortical plane of the lateral root cap, while those labelled as “Equatorial” (B) show stacked pictures immediately surrounding the quiescent center. Scale bar applies to all images within the figure. ACC = 0.2 µM 1-aminocyclopropane-1-carboxylic acid; *t*-Zea = 10 µM *trans*-zeatin; IAA = 50 nM indole-3-acetic acid; *TAA1: TRYPTOPHAN AMINOTRANSFERASE OF ARABIDOPSIS1*; LRC: lateral root cap; EPI: epidermis; TZ: transition zone; DZ: differentiation zone; RAM: root apical meristem. BF: bright field. Root zone boundaries are approximate.

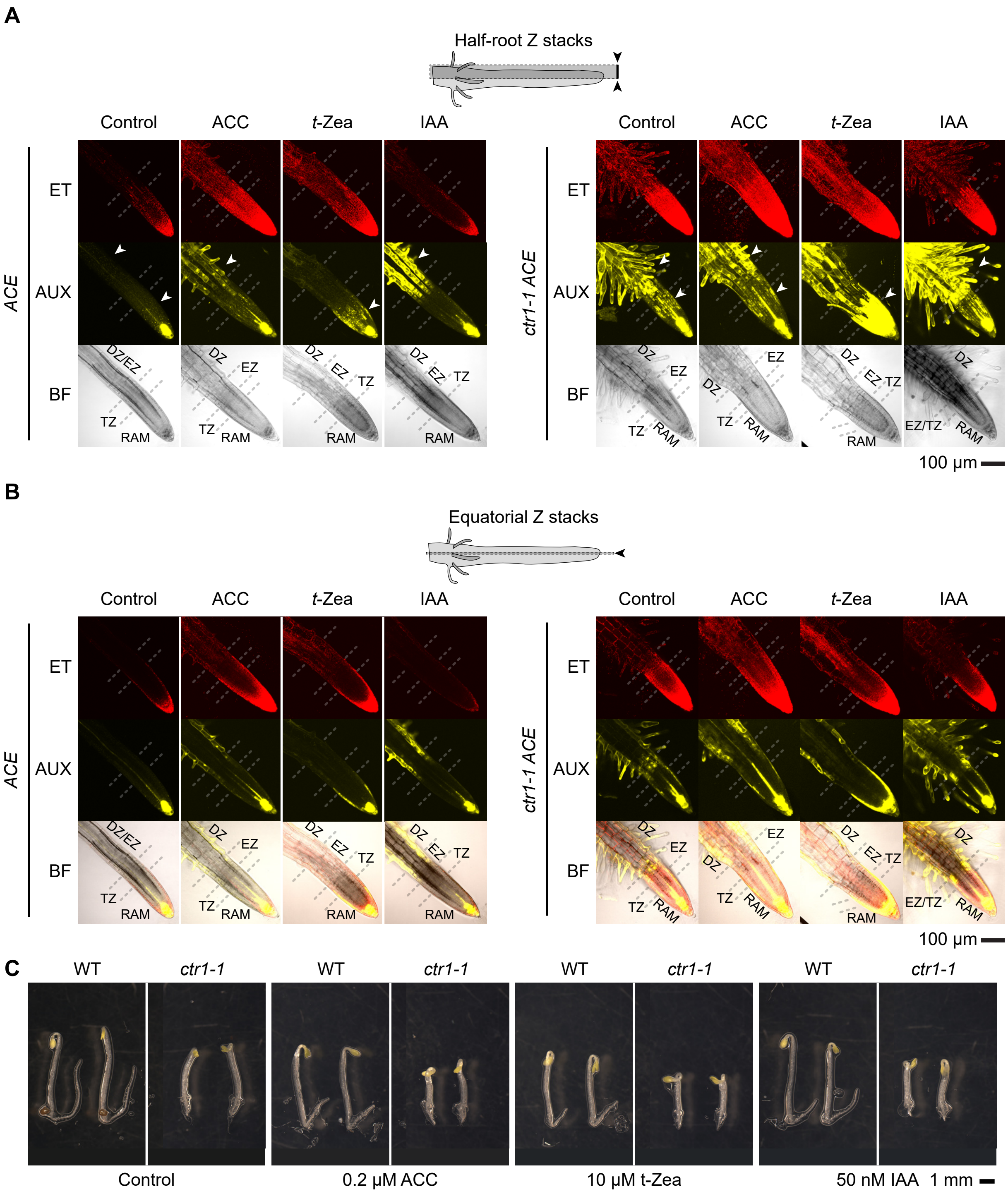


**Supplementary Figure S9. Constitutively active ethylene signaling in *ctr1* translates into more sensitized roots to exogenous hormones in three-day-old etiolated Arabidopsis seedlings.**

(A) Half-root Z-stacks showing that the disruption of the negative regulator CTR1, which exhibits the phenotype of ethylene-treated plants in the absence of ethylene, leads to strong *EBSn* reporter activity, higher basal auxin reporter activity in control conditions, and stronger induction of auxin reporter in response to exogenously applied hormones.

(B) Equatorial sections of the roots displayed in panel A.

(C) Seedling phenotypes in the presence or absence of hormone supplementation. Despite obvious differences in reporters' activity in response to exogenously applied hormones, *ctr1* shows very mild organ size changes in response to these growth regulators, suggesting a near saturation of the physiological and developmental responses to hormonal signals. Quantitative comparison and statistical assessment of hormone-specific inhibitory effect on growth between WT and mutants are available in Supplementary Figure S3.

All microscopy images were acquired in the same experimental session. Half root Z-stack (A) confocal microscopy images show stacked pictures from the equatorial plane at the quiescent center to the cortical plane of the lateral root cap, while those labelled as "Equatorial" (B) show stacked pictures immediately surrounding the quiescent center. Scale bar applies to all images within the panel. ACC = 0.2 µM 1-aminocyclopropane-1-carboxylic acid; *t*-Zea = 10 µM *trans*-zeatin; IAA = 50 nM indole-3-acetic acid. ET: ethylene reporter (*EBSn:mCherry-PTS1*); AUX: auxin reporter (*DR5v2:3xSV40-3xYPet*); BF: bright field. Root zone boundaries are approximate.

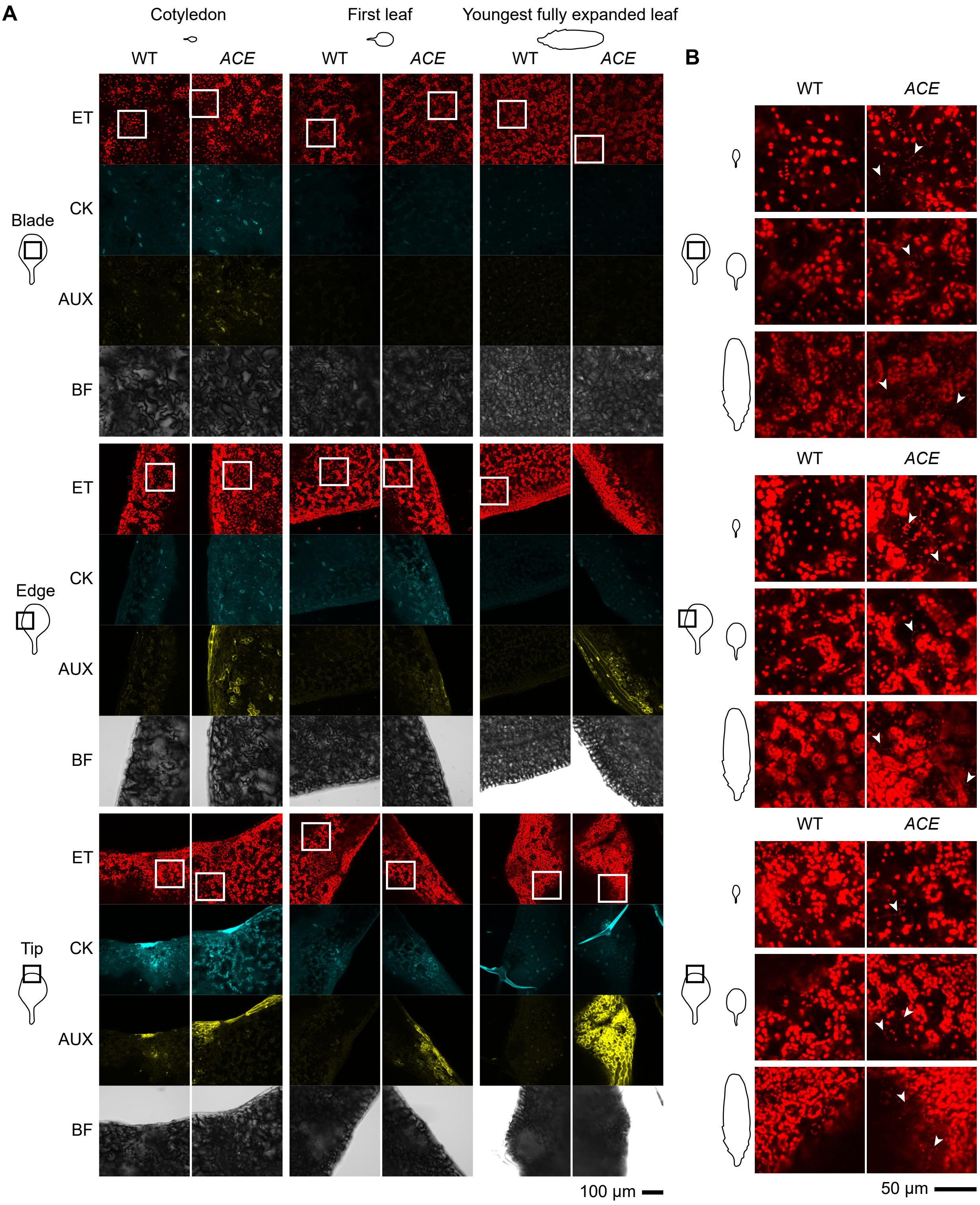


**Supplementary Figure S10. Expression of the *ACE* reporter in leaves of four-week-old Arabidopsis rosettes grown under long-day conditions.**

(A) *ACE* expression at the region of the leaf marked on the left-hand side cartoon (Top row: center of the leaf blade near the vasculature, Middle row: edge of the leaf, Bottom row: tip of the leaf) in adult cotyledons (left column), first true leaf of the rosette (middle column), and the youngest fully expanded leaf (right column).

(B) Digital magnification of images displayed in panel A showing peroxisome-targeted ethylene reporter's activity (arrows) between chloroplast autofluorescence activity. Confocal microscopy images show stacked pictures from mesophyll to epidermis cell planes.

All microscopy images were acquired in the same experimental session. Confocal microscopy images show stacked pictures spanning the full depth of leaf epidermal cells, extending into the palisade parenchyma. Scale bar applies to all images within the panel. ET: ethylene reporter (*EBSn:mCherry-PTS1*); CK: cytokinin reporter (*TCSn:MTS-mTagBFP**); AUX: auxin reporter (*DR5v2:3xSV40-3xYPet*); mTagBFP*: Hybrid version of mTagBFP that contains mTagBFP2-like amino acid substitution in mTagBFP chromophore moiety but does not include mTagBFP2 N-terminus; BF: bright field.

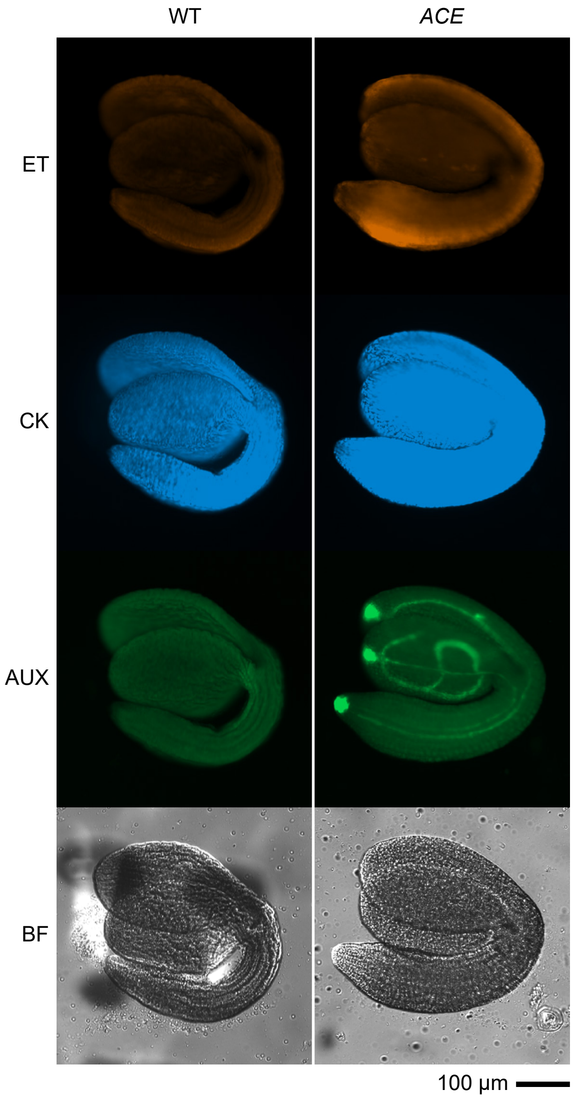


**Supplementary Figure S11.** ***ACE*** **activity in Arabidopsis embryos extracted from fully expanded siliques.**
The auxin reporter is active in the tips of the cotyledons, the root, and the vasculature, but weak-to-negligible ethylene reporter activity is observed. Strong background signal in the blue channel impairs the assessment of the cytokinin reporter.
Fluorescent microscopy images show stacked pictures from equatorial to cortical planes. Embryos were extracted by gently tapping on seeds embedded in a 15% glycerol solution using a microscopy glass and coverslip. ET: ethylene reporter (*EBSn:mCherry-PTS1*); CK: cytokinin reporter (*TCSn:MTS-mTagBFP**); AUX: auxin reporter (*DR5v2:3xSV40-3xYPet*); mTagBFP*: Hybrid version of mTagBFP that contains mTagBFP2-like amino acid substitution in mTagBFP chromophore moiety but does not include mTagBFP2 N-terminus; BF: bright field.

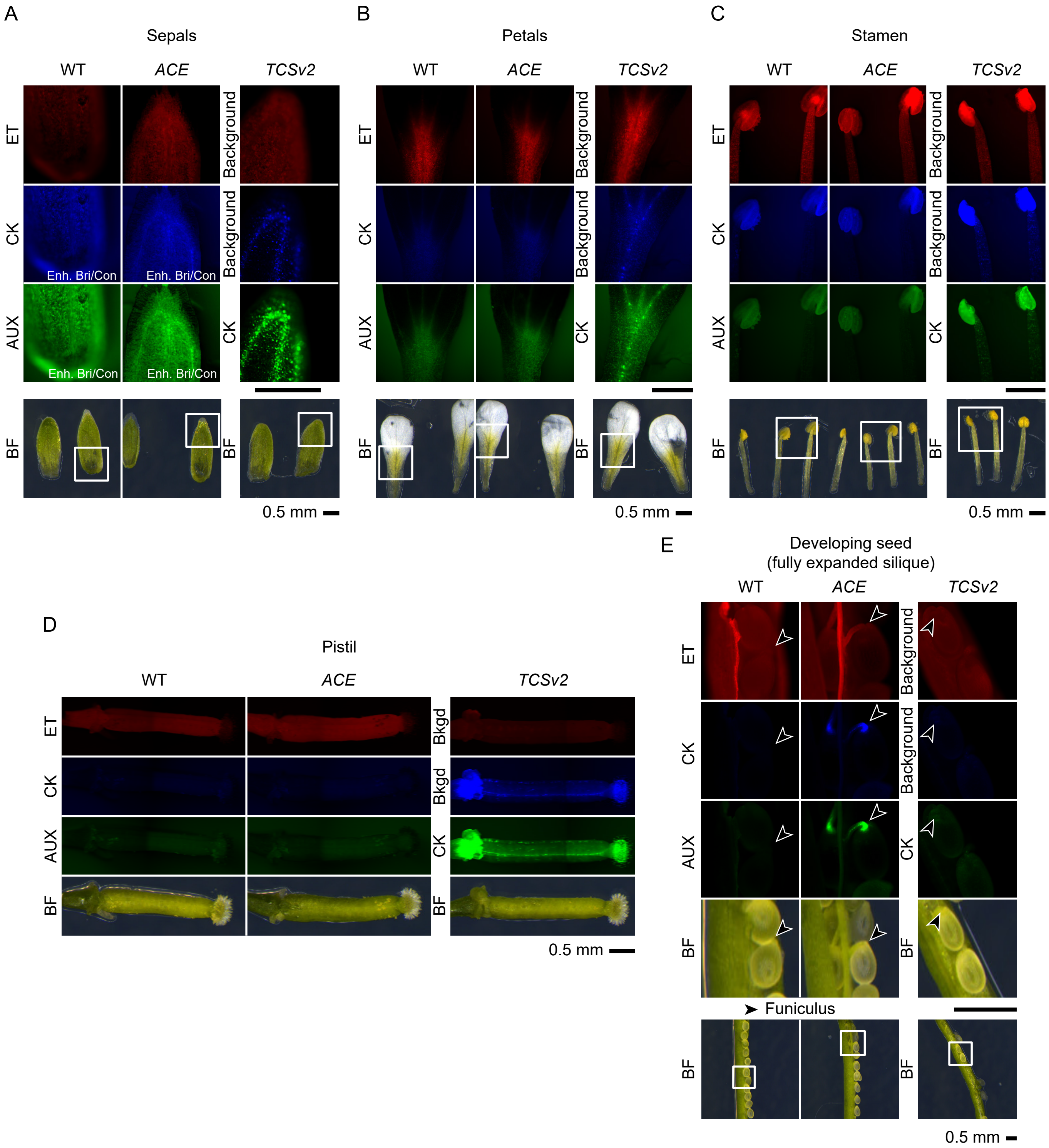


**Supplementary Figure S12.** ***ACE*** **expression in reproductive organs of Arabidopsis grown under long day conditions and its comparison to that of the cytokinin reporter,** ***TCSv2:3xmVenus-N7*****[49].**

The strong cytokinin activity detected in (A) sepals and (B) petal vasculature, (C) stamens, and (D) pistils by *TCSv2* is barely detected by *ACE*-encoded *TCSn*-driven cytokinin reporter due to the strong background signal. Conversely, fluorescence detected in *ACE*-encoded cytokinin and auxin reporters in (E) the funiculus (black arrow) connecting the seed and the replum was not detected in *TCSv2* cytokinin reporter, suggesting potential excitation of YPet in the 395–415 nm window and fluorescent emission in the 435–485 nm window *in planta*. Similarly, fluorescence activity patterns of ethylene reporter are indistinguishable from background fluorescence of wild type (WT).

All images were acquired in the same experimental session. Images whose contrast or brightness balance was modified after image acquisition for the purpose of better displaying fluorescent patterns are labelled as “Enh. Bri/Con”. Fluorescent microscopy images are single-plane pictures. All scale bars represent 0.5 mm and are applicable to all images within the fluorescence or bright field group for a given organ. Bkgd: background fluorescence; ET: ethylene reporter (*EBSn:mCherry-PTS1*); CK: cytokinin reporter (*TCSn:MTS-mTagBFP** (*ACE*) or *TCSv2:3xmVenus-N7* (*TCSv2*[49]); AUX: auxin reporter (DR5v2:3xSV40-3xYPet); mTagBFP*: Hybrid version of mTagBFP that contains mTagBFP2-like amino acid substitution in mTagBFP chromophore moiety but does not include mTagBFP2 N-terminus; BF: bright field.

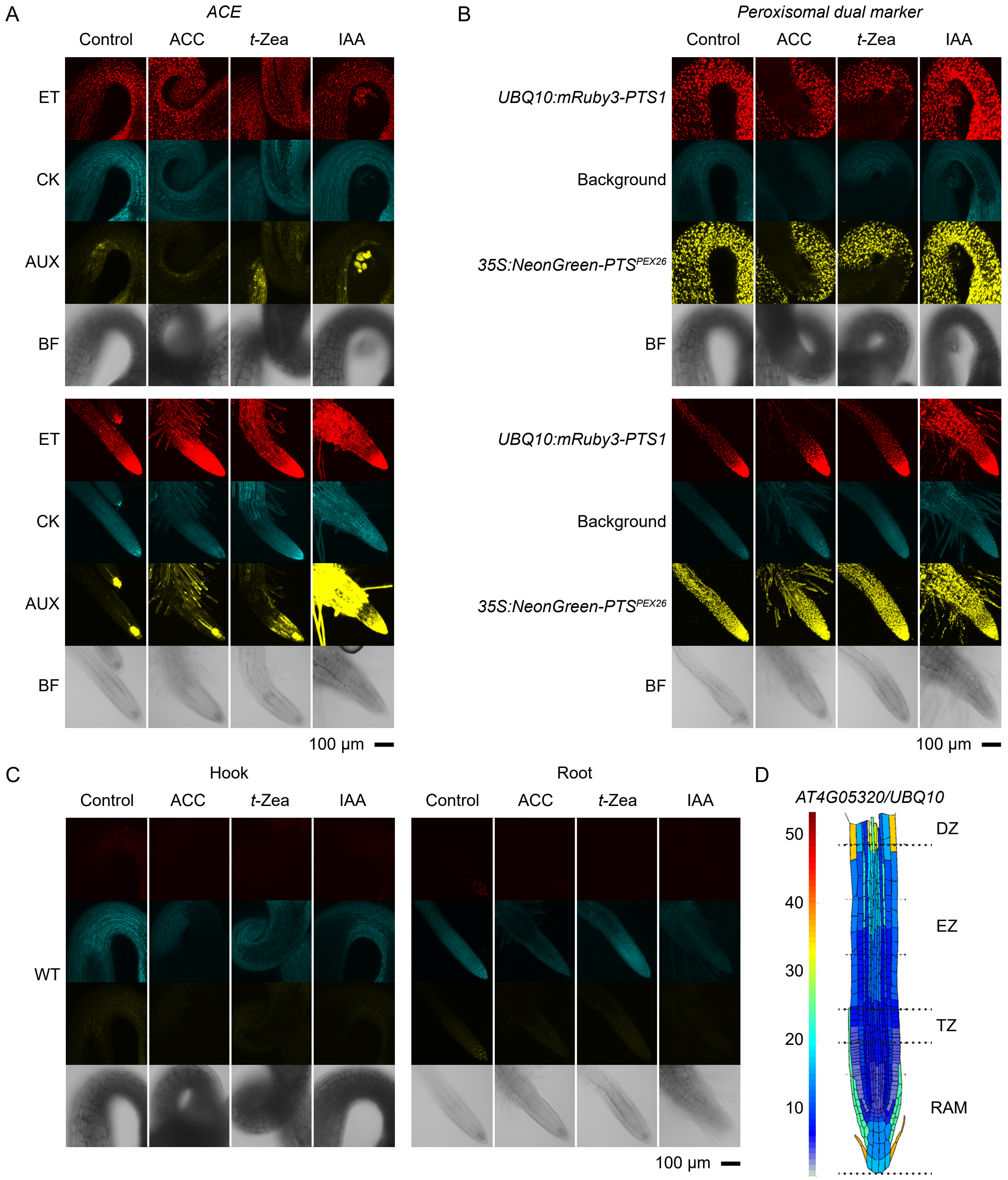


**Supplementary Figure S13. Exogenously applied hormones do not trigger prominent changes in peroxisomal localization or abundance in three-day-old etiolated Arabidopsis seedlings.**

(A) *ACE* reporter activity on unsupplemented or hormone-supplemented media in hooks and root tips.

(B) [*35Sp:NeonGreen-PTS$^{PEX26}$*] - [*UBQ10p:mRuby3-PTS1*][88] dual peroxisomal reporter activity on unsupplemented or hormone-supplemented media in hooks and root tips of seedlings grown alongside *ACE* seedlings shown in (A) under the same growth conditions.

(C) Background fluorescence in wild-type (WT) seedlings grown alongside *ACE* and peroxisomal reporters.

(D) Integrated single-cell relative expression of *UBQ10* in Arabidopsis roots. Modified from Root Cell Atlas (https://rootcellatlas.org/) output.

All microscopy images were acquired in the same experimental session. Confocal microscopy images show stacked pictures from the equatorial plane (at the widest section of the hook or at the quiescent center of the root) to the cortical plane (surface of the organ). Scale bar applies to all images within the panel. ACC = 10 μM 1-aminocyclopropane-1-carboxylic acid; *t*-Zea = 10 μM *trans*-zeatin; IAA = 1 μM indole-3-acetic acid. ET: ethylene reporter (*EBSn:mCherry-PTS1*); CK: cytokinin reporter (*TCSn:MTS-mTagBFP**); AUX: auxin reporter (*DR5v2:3xSV40-3xYPet*); mTagBFP*: Hybrid version of mTagBFP that contains mTagBFP2-like amino acid substitution in mTagBFP chromophore moiety but does not include mTagBFP2 N-terminus; BF: bright field.

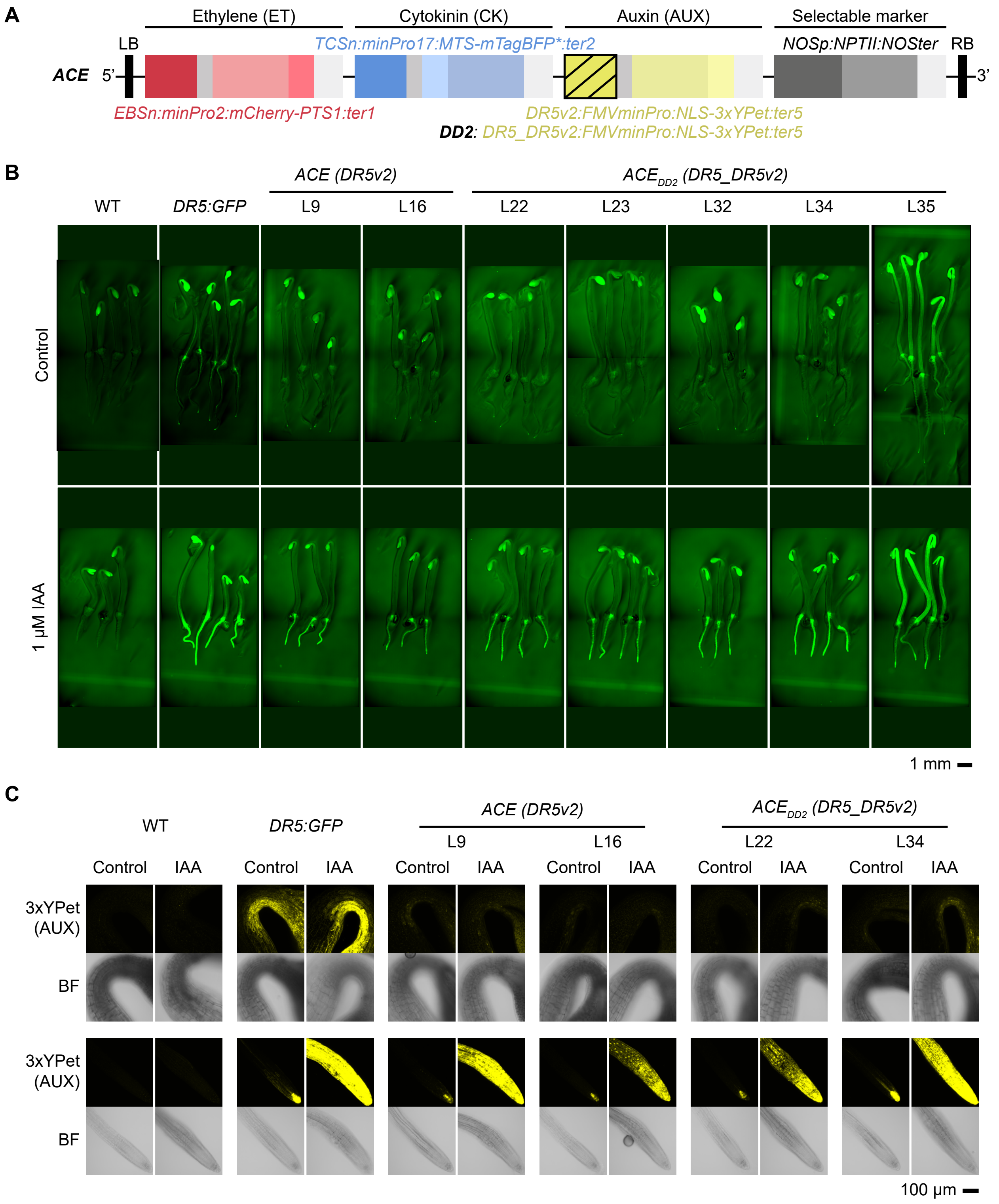


**Supplementary Figure S14. Comparison of auxin reporters activity contained in *ACE* (*DR5v2*) vs *ACE*$_{DD2}$ (*DR5_DR5v2*) activity in three-day-old etiolated Arabidopsis seedlings.**

(A) Schematic representation of the reporters, highlighting that the only difference between *ACE* and *ACE*$_{DD2}$ is the synthetic promoter driving auxin reporter.

(B) Fluorescent images of auxin reporter activity in three-day-old etiolated seedlings upon transfer onto control (AT) or IAA-supplemented plates. All images within the panel were acquired in the same experimental session.

(C) Confocal images of auxin reporter activity in seedlings presented in (B). All images within the panel were acquired in the same experimental session.

Seedlings were grown for 2.5 days in the dark on horizontal control plates (AT + 6 g L$^{-1}$ agar) and transferred onto vertical control (AT + 8 g L$^{-1}$) or supplemented plates with 1 µM indole-3-acetic acid (IAA). They were placed vertically and incubated for 12-16 h in the dark (12 h Thunder imaging (B), 16 h confocal (C)). Pictures in B are composite images stitched together using built-in LEICA software, or manually adjusted when the software failed to accurately assemble them (*ACE*$_{DD2}$ *L23*). Confocal microscopy pictures in C show stacked images from equatorial (at the quiescent center) to cortical (surface of the root) planes. AUX: auxin reporter *DR5v2:3xSV40-3xYPet* (*ACE*) or *DR5_DR5v2:3xSV40-3xYPet* (*ACE*$_{DD2}$); BF: bright field.

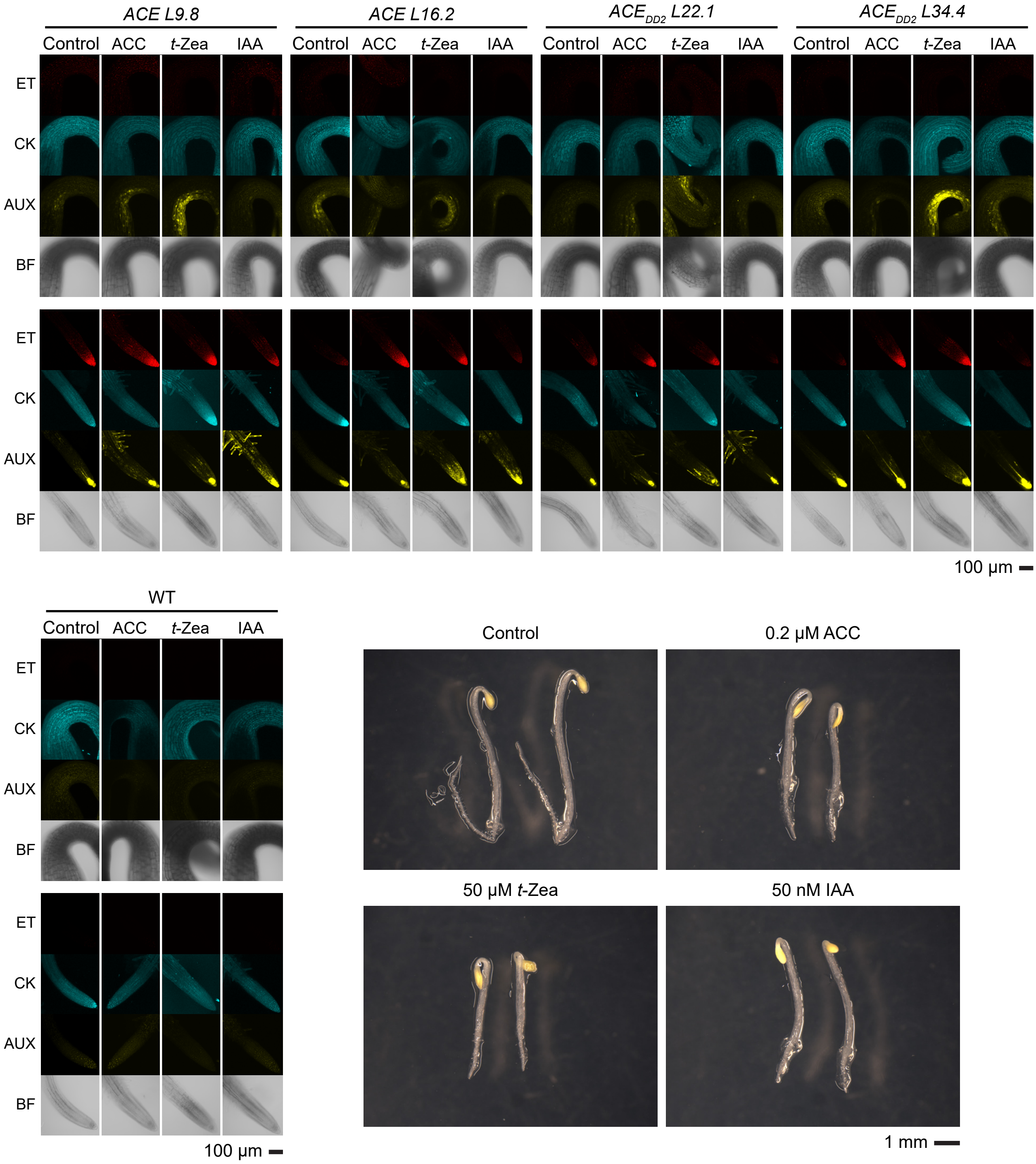


**Supplementary Figure S15. Comparative study of *ACE* and *ACE*$_{DD2}$ reporter expression in apical hooks and root tips of three-day-old etiolated Arabidopsis seedlings.**

Two independent lines per construct are displayed. All microscopy images were acquired in the same experimental session. Confocal microscopy images show stacked pictures from the equatorial to cortical planes. Seeds were germinated in the dark for three days in horizontal control (AT + 6 g $L^{-1}$ agar) plates or supplemented with individual hormones. ACC = 0.2 µM 1-aminocyclopropane-1-carboxylic acid; *t*-Zea = 50 µM *trans*-zeatin. Higher *t*-Zea concentration than in previous assays aimed to achieve greater induction of the cytokinin reporter, but no prominent differences were observed compared to 10 µM; IAA = 50 nM indole-3-acetic acid; ET: ethylene reporter (*EBSn:mCherry-PTS1*); CK: cytokinin reporter (*TCSn:MTS-mTagBFP**); AUX: auxin reporter *DR5v2:3xSV40-3xYPet* (*ACE*) or *DR5_DR5v2:3xSV40-3xYPet* (*ACE*$_{DD2}$); mTagBFP*: Hybrid version of mTagBFP that contains mTagBFP2-like amino acid substitution in mTagBFP chromophore moiety but does not include mTagBFP2 N-terminus; BF: bright field.

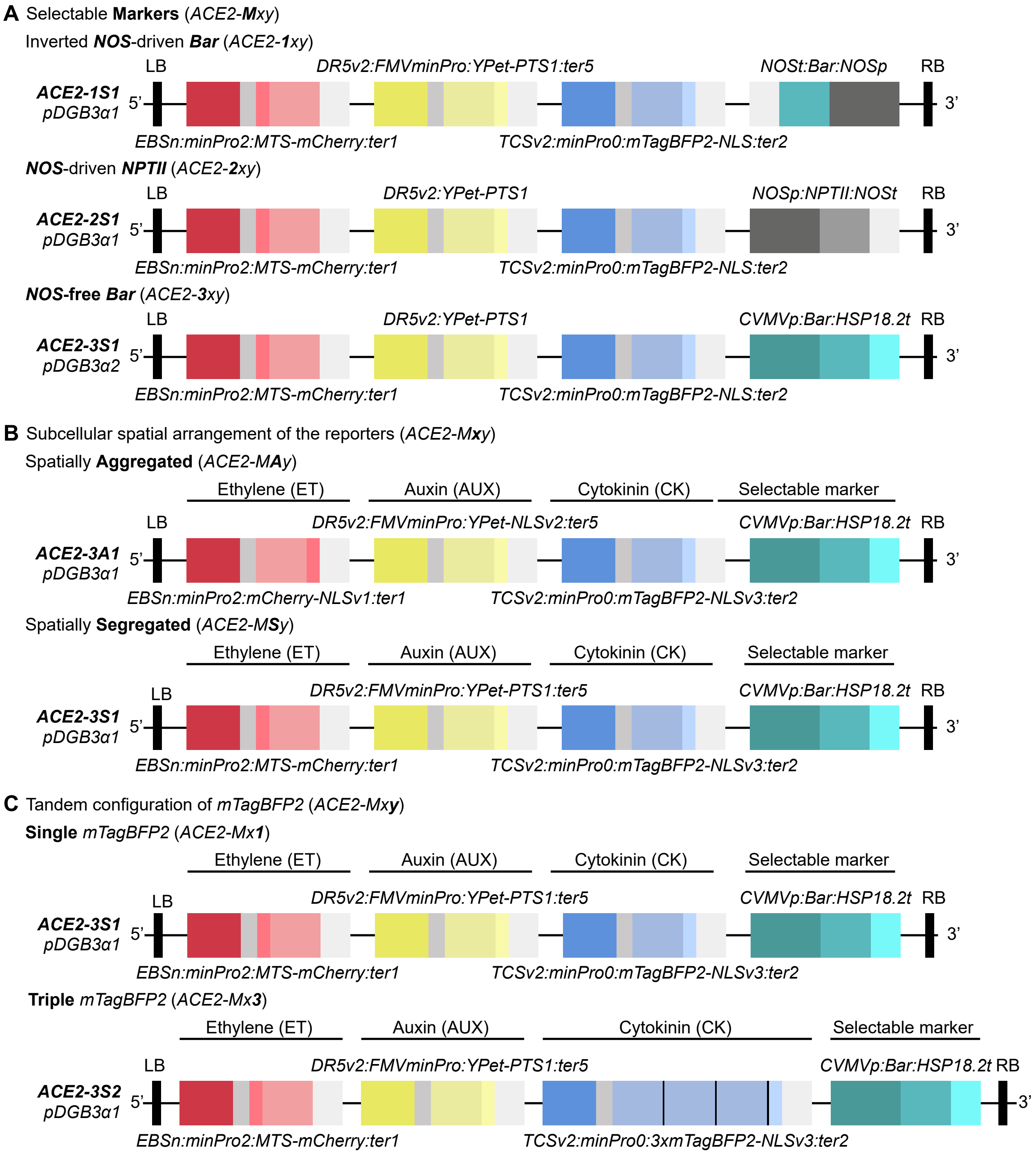


**Supplementary Figure S16. Schematic representation of *ACE2* variants and their nomenclature.**
The three-digit code after *ACE2* (*ACE2-Mxy*) reads as follows: (A) *M* refers to the selectable marker; (B) *x* refers to the spatial arrangement (*A*=aggregated, *S*=segregated); and (C) *y* indicates whether *mTagBFP2* is encoded by a single (*ACE2-Mx1*) or triple tandem (*ACE2-Mx3*) fluorescent protein. LB: left T-DNA border; RB: right T-DNA border; *PTS1: Peroxisomal Targeting Signal 1. MTS: Mitochondrial Targeting Signal; NLS: Nuclear Localization Signal*. *N7v1-N7v3* refers to three sequence-divergent coding DNA sequences for N7 NLS[89]. *CVMVp*: promoter from Cassava vein mosaic virus[63,65]; t*HSP18.2: HEAT SHOCK PROTEIN 18.2* terminator[116].

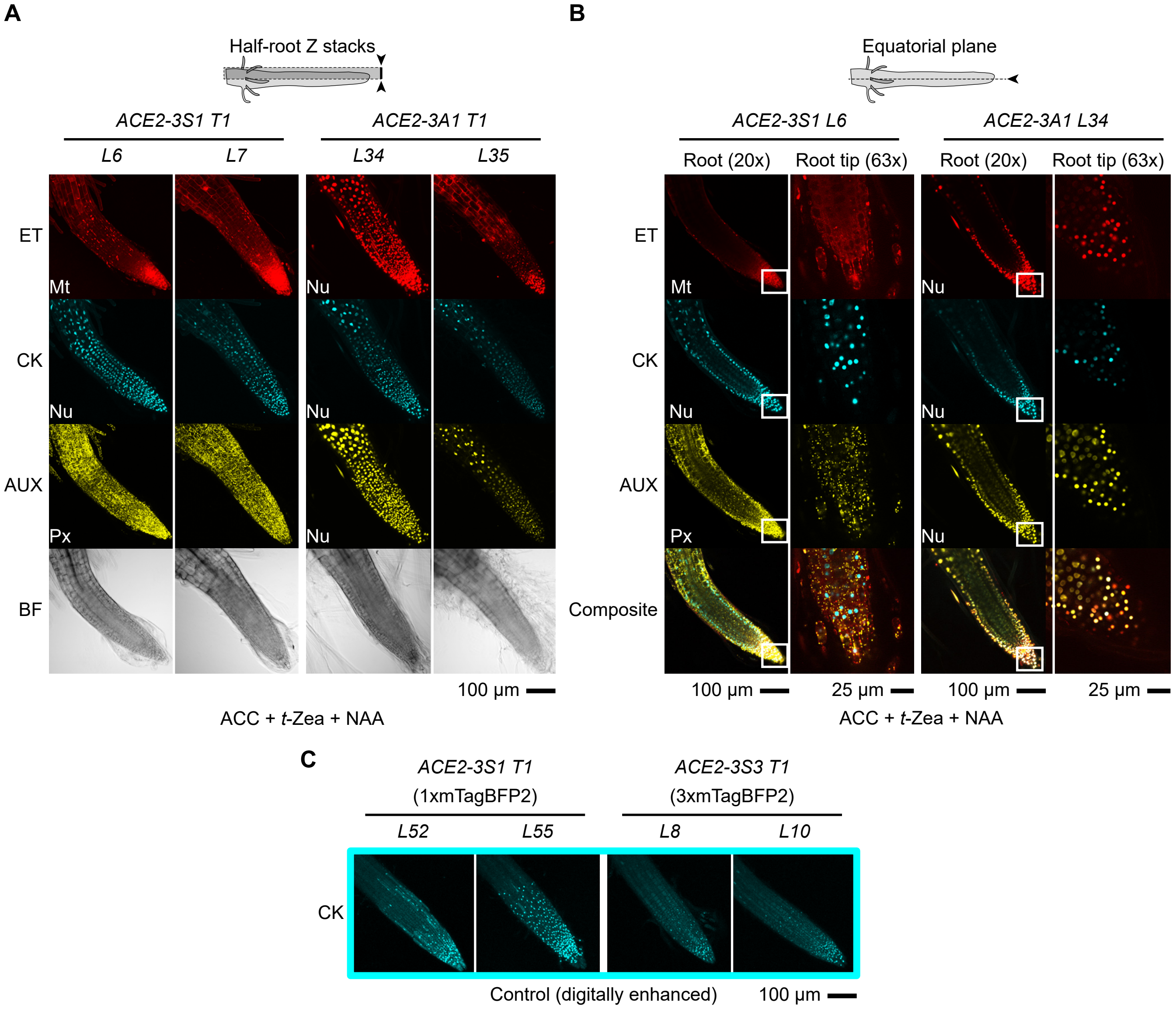


**Supplementary Figure S17. Characterization of the *ACE2* reporter variants in Arabidopsis root tips.**
(A) Half-root Z-stack images of two independent insertional lines of *ACE2* reporter in its two spatial arrangements. T1 seedlings show strong activity upon exogenous hormone supplementation and improved delivery to the targeted organelles relative to *ACE*.
(B) Single-plane images of the whole primary root and root tip of two lines displayed in (A). Details of the root tip were acquired with increased optical magnification of the regions marked with a white box.
(C) Half-root Z-stack images of two independent insertional events of *ACE2-3Sy* show that triple mTagBFP2 does not improve the detection of basal reporter activity compared to the single copy version under control conditions in T1 seedlings.
All microscopy images were acquired in the same experimental session. Seeds were germinated for three days in the dark on AT plates (6 g $L^{-1}$ agar) supplemented with 10 µg $mL^{-1}$ phosphinothricin and 300 µg $mL^{-1}$ timentin and grown for four additional days under continuous light. Resistant plants (with healthy cotyledons as opposed to those that bleached) were transferred to sugar-less AT (8 g $L^{-1}$ agar) plates supplemented with 300 µg $mL^{-1}$ timentin as control, or adding all three hormones (10 µM ACC + 30 µM *t*-Zea + 10 µM NAA) replacing IAA with the more photostable synthetic auxin, NAA, for 16 h prior to image acquisition. ET: ethylene reporter (*EBSn:MTS-mCherry*); CK: cytokinin reporter (*TCSv2:mTagB-FP2-N7*); AUX: auxin reporter (*DR5v2:YPet-PTS1*); BF: bright field.

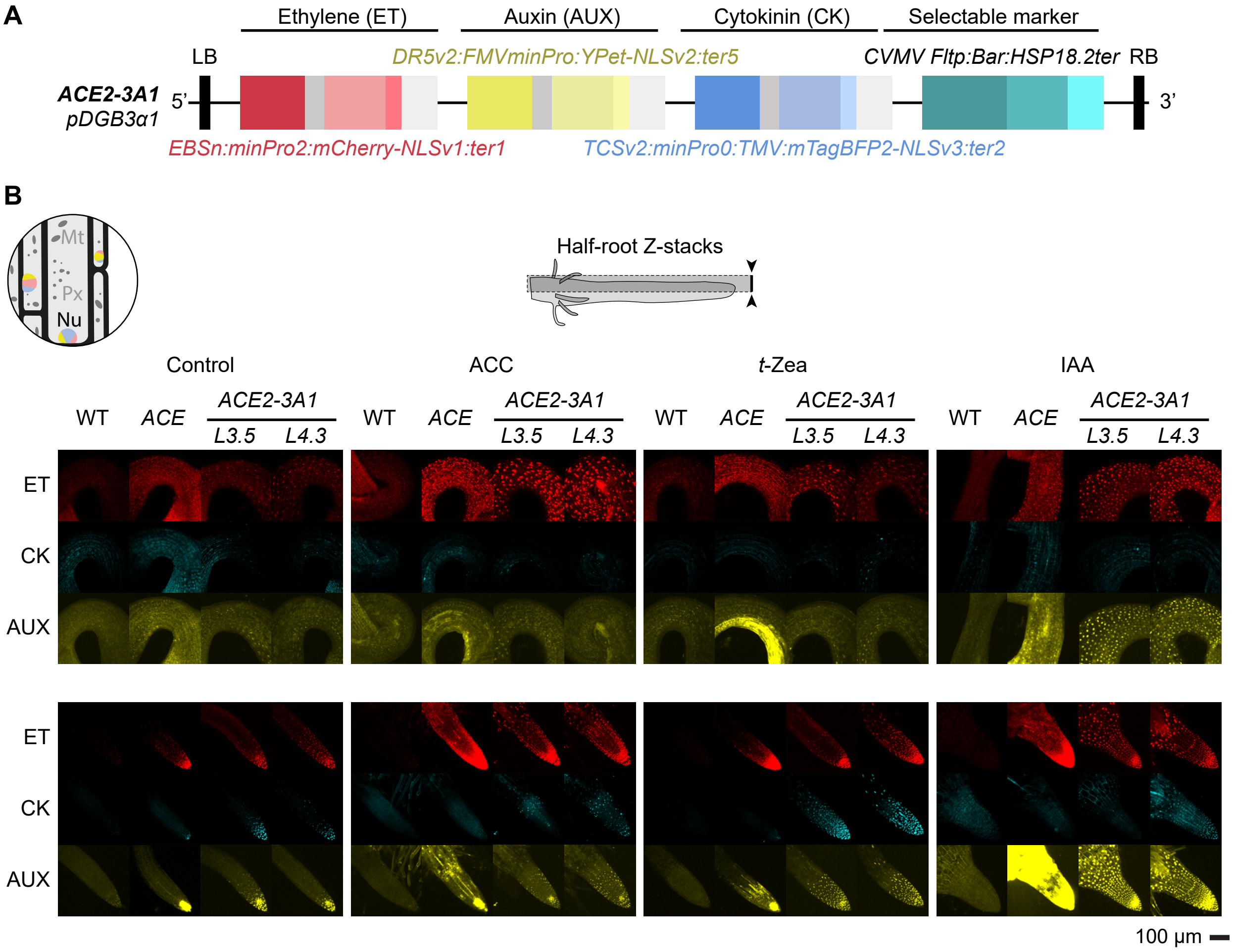


**Supplementary Figure S18. Characterization of the second-generation *ACE* reporter, *ACE2-3A1,* in stable Arabidopsis transgenic lines.**
(A) *ACE2-3A1* construct structure. Fragments not at scale.
(B) Basal reporter activity and hormone inducibility of *ACE2-3A1* compared to the first-generation *ACE* reporter in hooks (top) and roots (bottom) of Arabidopsis. Three-day-old etiolated seedlings were imaged under endogenous hormone concentrations (AT) and following supplementation with ACC, *trans*-zeatin, or IAA to selectively activate ethylene, cytokinin, and auxin signaling pathways, respectively. Comparison of reporter generations showed consistent patterns of activity and induction, with T5 generation being presented here.
Circular schematic shows that the expected subcellular localization of the fluorescent proteins is nuclear for all three reporters (Mt: mitochondria, Px: peroxisomes, Nu: nucleus). All microscopy images were acquired using identical acquisition settings for direct comparison between reporter generations. Confocal microscopy images show stacked pictures from equatorial to cortical planes (half-root Z-stacks). *L3.5* and *L4.3* refer to two independent single-insertion transgenic lines of *ACE2-3A1* reporter. Scale bar applies to all images within the panel. Images were digitally enhanced to facilitate detection of low-expressing cells and the same settings were applied to all images within a channel across treatments. LB: left T-DNA border; RB: right T-DNA border; *NLS: Nuclear Localization Signal (N7)*; ACC = 10 µM 1-aminocyclopropane-1-carboxylic acid; *t*-Zea = 10 µM *trans*-zeatin; IAA = 1 µM indole-3-acetic acid; ET: ethylene reporter (*EBSn*); CK: cytokinin reporter (*TCSv2*); AUX: auxin reporter (*DR5v2*); BF: bright field.

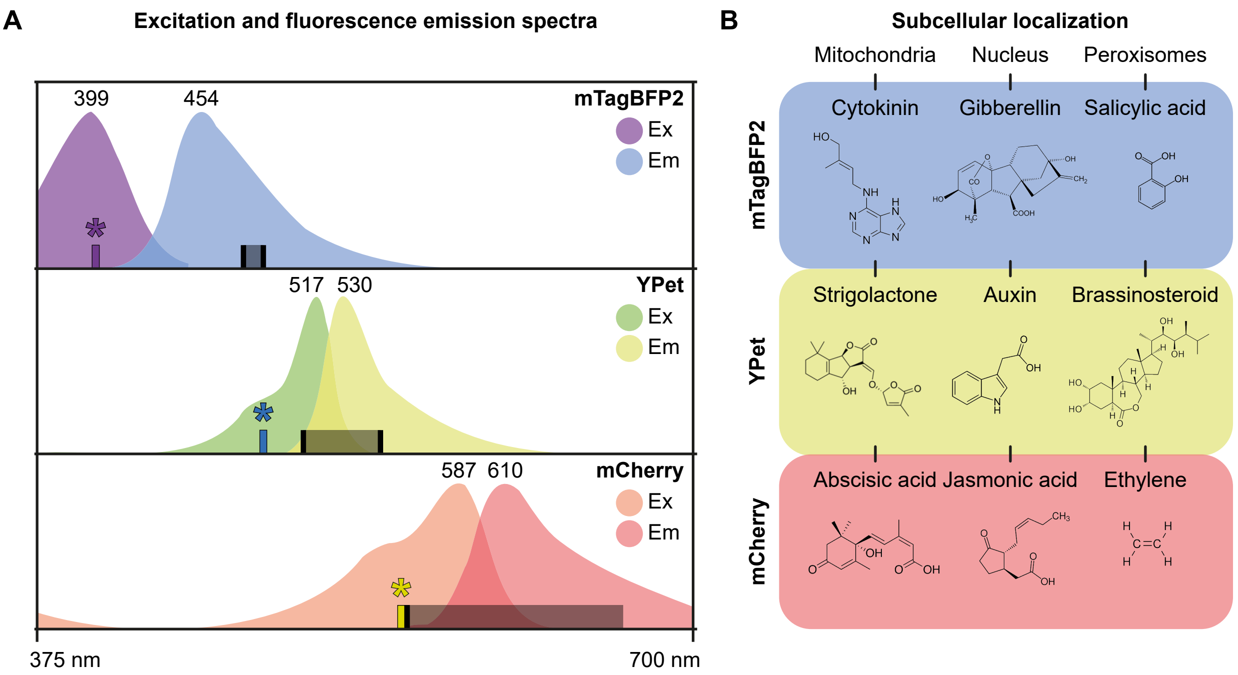


**Supplementary Figure S19. Logic to build a nonuple reporter to monitor phytohormone concentrations in vivo (*aka*, the hormometer).** Combining hormone-specific promoters responsive to all nine major non-peptidic phytohormone classes does not fully resolve the constraints of simultaneous in vivo detection imposed by spectral limitations. (A) Three fluorescence windows are robustly separable in plants: blue (*e.g.*, mTagBFP2), green–yellow (*e.g.*, YPet), and red (*e.g.*, mCherry). This limitation restricts conventional multiplexing strategies. However, spatial segregation of fluorescent proteins through subcellular targeting offers an orthogonal dimension for multiplexing (B). By combining three spectrally distinct fluorophores (mTagBFP2, YPet, and mCherry) with targeting to the nucleus, peroxisomes, and mitochondria, and driving each with distinct hormone-responsive promoters, it is theoretically feasible to generate a nonuple synthetic reporter capable of monitoring nine hormone classes in parallel.

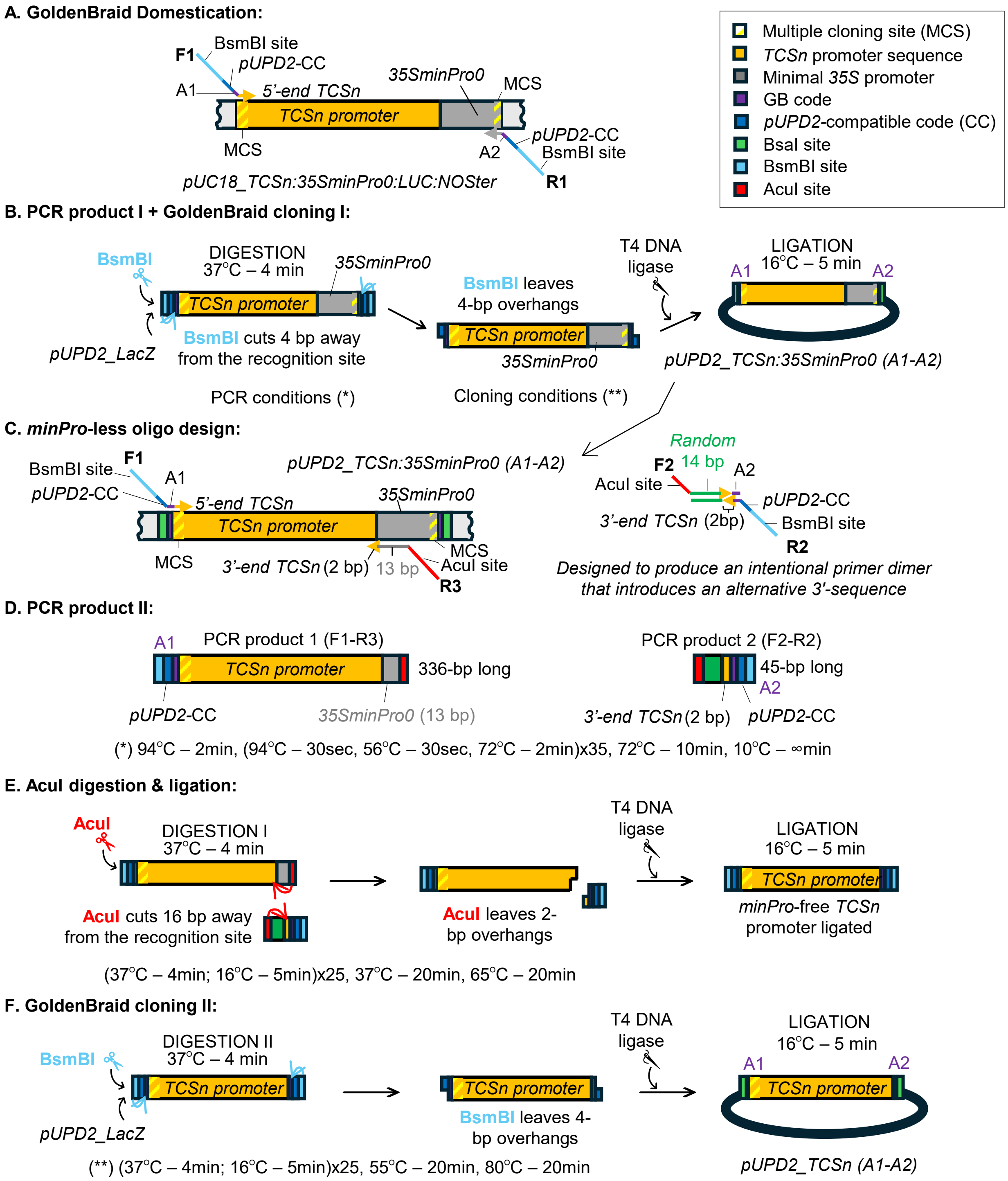


**Supplementary Figure S20. Cytokinin-inducible *TCSn* promoter subcloning strategy.** (A) GoldenBraid (GB) domestication of the *TCSn:35SminPro0* promoter. Oligos F1 and R1 (Supplementary Table S1A) were designed to amplify the *TCSn:35SminPro0* sequence from *pUC18_TCSn:35SminPro0:LUC:NOSter*[48] and introduce the DNA elements needed for its subcloning into *pUPD2* entry vector (with the GB code for a proximo-distal promoter (A1-A2) shown in purple, the *pUPD2* cloning overhang marked in dark blue, and BsmBI recognition sites indicated in cyan) in a similar fashion as previously described[51]. The amplification product included the 5' end of the *35SminPro0* sequence (dark grey) due to the difficulty of designing a reverse oligo over the repetitive *TCSn* promoter (yellow). (B) PCR and GB cloning of the *TCSn:35SminPro0* promoter. The amplification of the *TCSn:35SminPro0* promoter was done using iPROOF High-Fidelity DNA polymerase Kit (Bio-Rad Laboratories, 172-5302) with the following conditions: 94°C – 2min, (94°C – 30sec, 56°C – 30sec, 72°C – 2min)x35, 72°C – 10min, 10°C – ∞min. For subcloning the amplified promoter fragment into the *pUPD2* entry vector, previously described conditions[51] were used. Briefly: (37 °C – 4 min; 16 °C – 5 min)x25, 55 °C – 20 min, 80 °C – 20 min using BsmBI (NEB CutSmart R05580L) restriction enzyme and T4 DNA ligase (NEB M0202L). (C,D)**.** *minPro*-less oligo design and the resulting PCR products. Three additional oligos (F2, R2 and R3, Supplementary Table S1A) were created for the re-amplification of the *pUPD2_TCSn:35SminPro0* fragment and subsequent recloning of it without the *35SminPro0*. The R3 oligo was designed to include the AcuI restriction site (red) 14 bp downstream of the 3'-end of *TCSn* sequence. Together with F1, the two oligos amplified PCR product 1 (336 bp long). Oligos F2 and R2 are overlapping oligos designed to complement the sticky end of PCR product 1 while adding the *pUPD2* cloning overhang to complete a proximo-distal promoter DNA element (A1-A2); they amplified PCR product 2 (45 bp long). A random sequence of 14 bp was included between the AcuI recognition site and the 2 bp long *TCSn* sequence (sticky end). Same PCR conditions as in panel B were used. (E) AcuI digestion and ligation. This step was designed to generate a *minPro*-less *TCSn* sequence flanked by the *pUPD2* cloning overhangs. For that, the AcuI restriction enzyme cut 16 bp away from the recognition site and left 2 bp sticky ends in both PCR products, 1 and 2. When ligated by T4 DNA ligase, PCR product 2 filled the void made by AcuI in PCR product 1, regenerating the *TCSn* promoter while removing the *35SminPro0* sequence. The cloning conditions for this reaction were (37 °C – 4 min; 16 °C – 5 min)x25, 37 °C – 20 min, 65 °C – 20 min. **F.** *minPro*-less *TCSn* GB cloning into *pUPD2* entry clone. Following a similar GB assembly strategy as mentioned in panel B, the *minPro*-less *pUPD2_TCSn* clone was generated (Supplementary Table S1A).

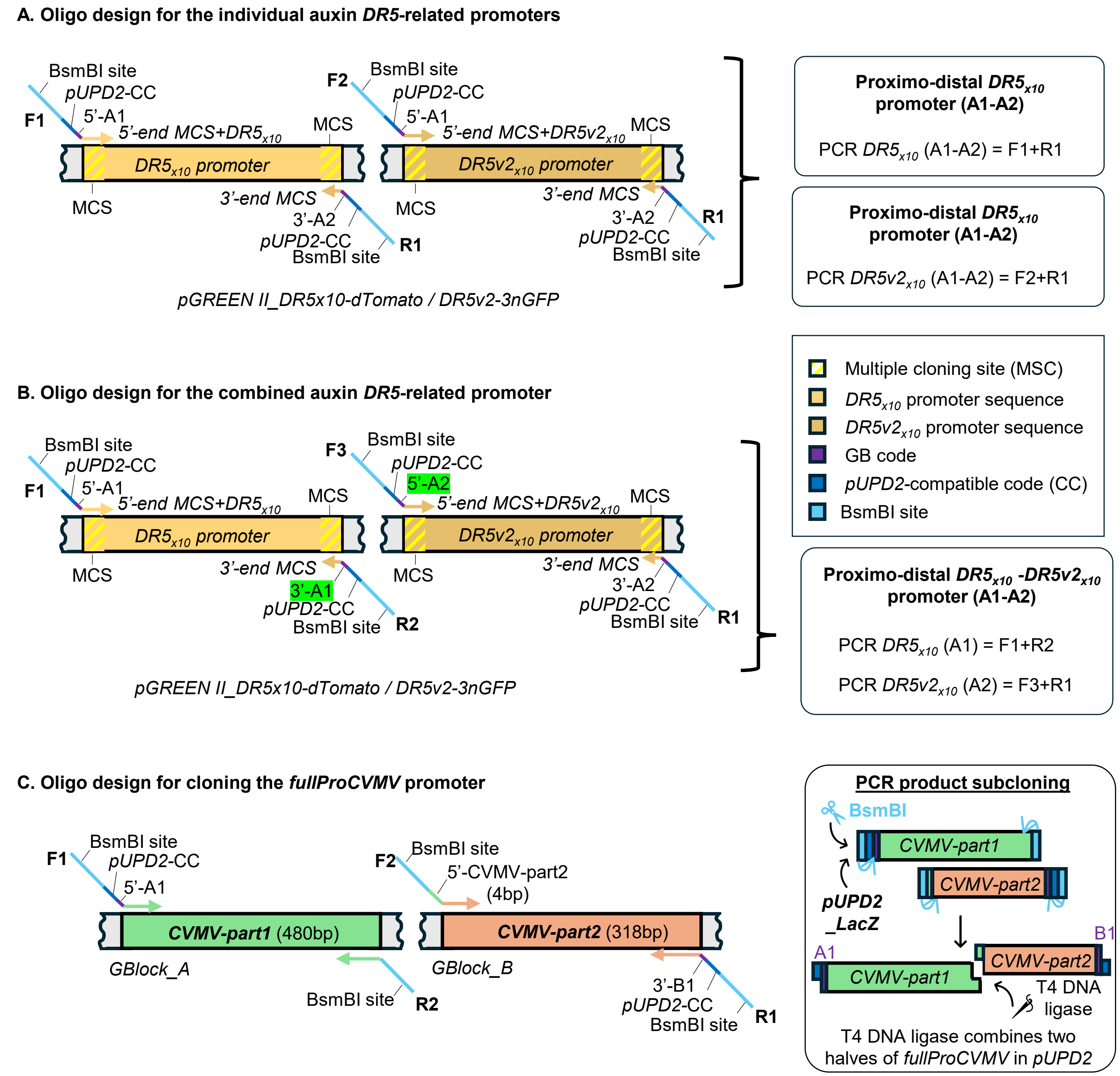


**Supplementary Figure S21. Oligo design strategies for auxin *DR5*-related and *fullProCVMV* promoters.** (A) Oligo design for the individual auxin-inducible *DR5*$_{x10}$ and *DR5v2*$_{x10}$ proximo-distal (A1-A2) promoters. F1 and F2 oligos (Supplementary Table S1A) were designed over the 5'-multiple cloning site (MSC) region and the first *DR5* or *DR5v2* repeat, respectively, to provide sequence specificity while ensuring primer annealing to the first repeat of these repetitive promoters. Additionally, the oligos are flanked by a 5'-A1 GB code (purple), a *pUPD2* cloning overhang (dark blue), and a BsmBI recognition site (cyan) for their subcloning into *pUPD2* entry vector. The reverse oligo (R1) was made complementary to the common 3'-MCS region and was flanked by the same cloning elements as F1 and F2 but adding the 3'-A2 GB code instead. (B) Oligo design for the individual auxin-inducible *DR5*$_{x10}$*:DR5v2*$_{x10}$ proximo-distal (A1-A2) promoter. Two additional oligos were designed (F3 and R2, Supplementary Table S1A) to amplify DR5$_{x10}$ and DR5v2$_{x10}$ as distal (A1) and proximal (A2) promoters, respectively, and allow their assembly during their subcloning into *pUPD2* entry vector to create a final combined *DR5*$_{x10}$*:DR5v2*$_{x10}$ proximo-distal (A1-A2) promoter. For that, the 5'-A1 GB code in F2 oligo was changed to the 5'-A2 GB code (highlighted in green) to create F3. Likewise, the 3'-A2 GB code in R1 oligo was switched to the 3'-A1 GB code (highlighted in green) to create R2. The template used for amplifying *DR5*$_{x10}$, *DR5v2*$_{x10}$, and *DR5*$_{x10}$*:DR5v2*$_{x10}$ proximo-distal promoters was *pGREEN II_DR5x10-dTomato / DR5v2-3nGFP*[47]. The PCR products obtained with each oligo pair combination are displayed on the right side of the figure. (C) Oligo design for subcloning the *Cassava vein mosaic virus* (*CVMV*) promoter. The full-length *CVMV* promoter was split into two independent GBlocks to pass the quality thresholds set by the vendor (Integrative DNA Technologies) for its commercial synthesis. For that, a *CVMV* 480 bp fragment was included in the GBlock_A, and the remaining *CVMV* 318 bp fragment was synthesized as the GBlock_B. Four oligos (F1, F2, R1, and R2, Supplementary Table S1B) were designed to amplify both fragments and allow their scar-less assembly into the *pUPD2* entry clone. The oligos F1 and R1 harbor the GB codes (A1 and B1, respectively) shown in purple, the *pUPD2* cloning overhangs displayed in dark blue, and the BsmBI recognition sites marked in cyan, enabling the fragment subcloning into the entry vector. Oligos F2 and R2 are flanked by the BsmBI recognition sites (cyan) to generate compatible 4 bp sticky ends. During the regular GB assembly conditions[98], the T4 DNA ligase will combine the two PCR fragments with the vector, thus regenerating the full-length of *fullProCVMV* promoter. PCR conditions employed herein are the same as described in Supplementary Figure S20.
