## Supplemental table for "An Integrated Spatially Resolved Mechanistic Model of Hierarchical Auxin–Cytokinin–Ethylene Crosstalk Underlying Root Growth Inhibition in Arabidopsis"

**Supplementary Table S1. *pUPD2* entry clones and selectable markers created for the different GB parts assembled into transcriptional reporters**

**A**

| DNA part | GB category | Clone name | Link to the clone (Benchling supported) | DNA primers for cloning into pUPD2 entry vector: Primer forward | DNA primers for cloning into pUPD2 entry vector: Primer reverse |
|---|---|---|---|---|---|
| **HORMONE-INDUCIBLE PROMOTERS** | | | | **Primer forward** | **Primer reverse** |
| *Auxin Distal-Proximal inducible promoter, $DR5v2_{x10}$* | A1-A2 | *pUPD2_10xDR5v2* | https://benchling.com/s/seq-gW9X2HhWCCG3ShiG3CMM?m=slm-3jC2i4Lh637C | F2: GCGCCGTCTCACTCGGGAGGCCTAGTTGGAATAGGATTTCG | R1: GCGCCGTCTCACTCAGGGACAAGCTTATCGAGGCCTGC |
| *Auxin Hybrid Distal-Proximal inducible promoter, $DR5_{x10}$:$DR5v2_{x10}$* | A1-A2 | *pUPD2_10xDR5-10xDR5v2* | https://benchling.com/s/seq-RWDSCPAoXfFZw37s12Fu?m=slm-ylVz0AJvBjVl8D8 | CGCCGTCTCACTCGGGAGGGTGTCATCTATGTGAATTCGAGAC; F3: GCGCCGTCTCATGACGCCTAGTTGGAATAGGATT | R1: GCGCCGTCTCACTCAGGGACAAGCTTATCGAGGCCTGC; R2: GCGCCGTCTCAGTCACAAGCTTATCGAGGCCTGC |
| *Cytokinin Distal-Proximal inducible promoter, TCSn1* | A1-A2 | *pUPD2_TCSn1* | https://benchling.com/s/seq-2F9mnIZAMAhIkn30jKI5?m=slm-FeIJ9tanjMiLMyHtZA | F1: GCGCCGTCTCGCTCGGGAGGGTCGGCTAAAAGCTTGACTAGTC; F2: GCGCCTGAAGCCTTCGCAAGACCCTATCCC | R3: GCGCCTGAAGGGGTCTTGGCTAGCTAAAGA; R2: GCGCCGTCTCACTCAGGGATAGGGTCTTGCGAAGG |
| *GB-domesticated cytokinin Distal inducible promoter, 1/2 TCSv2* | A1 | *pUPD2_TCSv2_half1* | https://benchling.com/s/seq-M78y70Yj6rzT04PBEBVF?m=slm-KwQ62yFYrhegeF | GCGCCGTCTCGCTCGGGAGGACTAGTCAAAGATTTTGCAAAATC | GCGCCGTCTCGCTCAGTCAGTTGGATTTTGAAAGATCTCTCTA |
| *GB-domesticated cytokinin Proximal inducible promoter, 1/2 TCSv2:35SminPro0:TMV* | A2-B1 | *pUPD2_TCSv2_half2+35Smp0+TMV* | https://benchling.com/s/seq-cprF9wjD9ewc2rL1T8bG?m=slm-Bz5kNkgZcZei8GTx | GCGCCGTCTCGCTCGTGACGACTAGTCAAAGATTTTGCAAAATC | GCGCCGTCTCGCTCAATGGACGCGTTAATTGTAATTGTAAATAGT |

GB: GoldenBraid. *10xDR5* and *10xDR5v2* come from *pGREEN II_DR5x10-dTomato / DR5v2-3nGFP*[47] and were independently amplified via PCR using primers containing GoldenBraid (GB) overhangs for parts A1 and A2, respectively. Both amplicons were then assembled in the *pUPD2* vector in a single-pot reaction, with the PCR fragment $DR5_{x10}$ (GB A1-A2) amplified with primers F1+R1 and the PCR fragment $DR5v2_{x10}$ (GB A1-A2) amplified with primers F2+R1 to generate two alternative proximo-distal promoters (Supplementary Figure S21A). The PCR fragment $DR5_{x10}$ (GB A1) was amplified with primers F1+R2, whereas the PCR fragment $DR5v2_{x10}$ (GB A2) was amplified with primers F2+R3, and the two PCR products were combined together to generate a third proximo-distal promoter (Supplementary Figure S21B).*TCSn* came from *pUC18_TCSn:35SminPro0:LUC:NOSter*[48]. The repetitiveness of the *TCSn* sequence necessitated its subcloning as *pUPD2_TCSn:35SminPro0*, with primers F1+R1 used to amplify the promoter. To make the *TCSn* construct fully modular and compatible with different *minPro* of the collection, the *35SminPro0* was removed with the help of a non-GB Type IIS enzyme AcuI by leveraging PCR1 that used primers F1+R3 and PCR2 that relied on primers F2+R2, and combining the two PCR products in pUPD2 using AcuI, BsmBI, and DNA ligase (Supplementary Figure S20). Original *TCSv2:35SminPro0:TMV* was described in[49]. TMV: Tobacco mosaic virus translational enhancer.

**B**

| DNA part | GB category | Clone name | Link to the clone (Benchling supported) | Primer forward | Primer reverse |
|---|---|---|---|---|---|
| **VIRAL CONSTITUTIVE PROMOTERS** | | | | **Primer forward** | **Primer reverse** |
| CaMV *35Sp* | A1-B1 | *GB0030* | https://goldenbraidpro.com | Ready-to-use piece from the original GB collection (GB0030) | |
| SVBV *Sgt VIp* | A1-B1 | *pUPD2_fullProSVBV* | https://benchling.com/s/seq-8Xtrgip1QqgugPITiaTn?m=slm-sU7UEGAPr47LcqSV | GCGCCGTCTCGCTCGGGAGGACCACTTGTGACTAAGGATC | GCGCCGTCTCGCTCAATGGCCTTCAAATGAAAATACAAGCTCT |
| FMV *Sgt VIp (19S)* | A1-B1 | *pUPD2_fullProFMV* | https://benchling.com/s/seq-rlt4NeA5L7TmVz9NNvTl?m=slm-PaDTSLnICDOxFhev | GCGCCGTCTCGCTCGGGAGCTCGAGCAAGTGTACAAAAAAG | GCGCCGTCTCGCTCAATGGATATTGGTTGAGTATTGATGATCTTC |
| CVMV *Fltp* | A1-B1 | *pUPD2_fullProCVMV* | https://benchling.com/s/seq-SC2C7tGn4YKAKuk9XK2t?m=slm-2RoSi1bcGLno55 | CCGTCTCGCTCGGGAGCTTAATTAATGGTGGACTCCTG; F2: GCGCCGTCTCACTAGTGAAGACGTAAGCACTGACGACAA | CCGTCTCGCTCAATGGTTCGGATCCTTCACCACAAAC; R2: GCGCCGTCTCACTAGTGACACGGAAAAATATAAAAGGATAAGTAGTGG |

Full promoters from the Cauliflower mosaic virus 35S (CaMV 35Sp; 1027 bp)[51], the Strawberry vein banding virus Sgt VI[61,62] (SVBV VIp; 988 bp), the Figwort mosaic virus M3 strain Sgt VI (19S)[63,64] (FMV 19Sp; 416 bp), and the Cassava vein mosaic virus Flt[63-65] (CVMV Fltp; 2895 bp). DNA sequences were amplified from GBlock synthetic DNA. The synthesis of fullProCVMV was split into two GBlocks to be commercially synthesized, with the two fragments amplified (PCR1 used primers F1+R2 and PCR2 relied on primers F2+R1) and combined during regular GB subcloning protocol into pUPD2 (Supplementary Figure S21C). Sgt: subgenomic transcript; Flt: full-length transcript; VI: sixth open reading frame (ORF).

**C**

| DNA part | GB category | Clone name | Link to the clone (Benchling supported) | Primer forward | Primer reverse |
|---|---|---|---|---|---|
| **CORE PROMOTERS** | | | | **Primer forward** | **Primer reverse** |
| CaMV *35S minimal promoter #0 (minPro0)* | A3-B1 | *pUPD2_minPro0* | https://benchling.com/s/seq-3ETRSexQ5VFQmDTUwhub?m=slm-92qr4pd9WRYh | GTAATACGACTCACTATAGGGC | AATTAACCCTCACTAAAGGG |
| CaMV *35S minimal promoter #1 (minPro1)* | A3-B1 | *pUPD2_minPro1* | https://benchling.com/s/seq-fKfGai0ADotCfto01Bro?m=slm-SptfjLesutomW06mN7 | GTAATACGACTCACTATAGGGC | AATTAACCCTCACTAAAGGG |
| CaMV *35S minimal promoter #2 (minPro2)* | A3-B1 | *pUPD2_minPro2* | https://benchling.com/s/seq-OPNeEtn2tjydZlcETOQ3?m=slm-KJZiwZaaWVrVFksll | GTAAAACGACGGCCAGTG | GGAAACAGCTATGACCATG |
| CaMV *35S minimal promoter #3 (minPro3)* | A3-B1 | *pUPD2_minPro3* | https://benchling.com/s/seq-c3NBMHL76ay1MStuGJWS?m=slm-WlPe2Uf9skFLyr | GCGCCGTCTCACTCGTCCCCTCTCTAGCAATGAG | GCGCCGTCTCACTCAATGGTCCTACTTGCCCAAG |
| CaMV *35S minimal promoter #4 (minPro4)* | A3-B1 | *pUPD2_minPro4* | https://benchling.com/s/seq-rANQZTuOCtlOzkd9yPwa?m=slm-sw6zMl7645mww9 | GCGCCGTCTCACTCGTCCCAGTTTAGGGGTCC | GCGCCGTCTCACTCAATGGGGACCCCCAACTC |
| CaMV *35S minimal promoter #5 (minPro5)* | A3-B1 | *pUPD2_minPro5* | https://benchling.com/s/seq-0X7WKErrIFxqCvGWlGK7?m=slm-alpGXqYqlsNF6nc | GCGCCGTCTCACTCGTCCCGGTAGTCATGGTCAT | GCGCCGTCTCACTCAATGGCTCTTCTATCCTAC |
| CaMV *35S minimal promoter #6 (minPro6)* | A3-B1 | *pUPD2_minPro6* | https://benchling.com/s/seq-DvaOPs9puWESALezpvG5?m=slm-R1Eiqy2HTssm9 | GCGCCGTCTCACTCGTCCCGGGGGGGATGAAAGT | GCGCCGTCTCACTCAATGGGAAAGAGTGAATAGTG |
| CaMV *35S minimal promoter #7 (minPro7)* | A3-B1 | *pUPD2_minPro7* | https://benchling.com/s/seq-cUFLKdWgiDSl44gbUy76?m=slm-w3PFFUHL8LVmQ | GCGCCGTCTCACTCGTCCCGAGTAAGGGATTCCC | GCGCCGTCTCACTCAATGGAGCCAAGCTCACTC |
| CaMV *35S minimal promoter #8 (minPro8)* | A3-B1 | *pUPD2_minPro8* | https://benchling.com/s/seq-MLvBj0xWth4rwz1KvzD9?m=slm-cUdgV8xSCwmkmL | GCGCCGTCTCACTCGTCCCGTGGAAAGGCTCAC | GGTCTAGAGATTTCTCCTCCATTGAGTGAGACGGCGC |
| CaMV *35S minimal promoter #9 (minPro9)* | A3-B1 | *pUPD2_minPro9* | https://benchling.com/s/seq-0S1YRhZq5fl9c49EHzJ9?m=slm-RO3zlYoOPUL03fS | GCGCCGTCTCACTCGTCCCATAGGGTCCCCCTTG | GCGCCGTCTCACTCAATGGATAGGCATCCCCTAG |
| CaMV *35S minimal promoter #10 (minPro10)* | A3-B1 | *pUPD2_minPro10* | https://benchling.com/s/seq-H8NgKv9d1rCDFxqR15Cc?m=slm-Q5lyH8NrKi2otEXr | GCGCCGTCTCACTCGTCCCGCAAACCCATGGCAT | GCGCCGTCTCACTCAATGGGGGCCAAGTTAAACAAAG |
| CaMV *35S minimal promoter #11 (minPro11)* | A3-B1 | *pUPD2_minPro11* | https://benchling.com/s/seq-z3aEO7E6VbIIhA9K6IeQ?m=slm-MFKFMGTFGKIOu | GCGCCGTCTCACTCGTCCCTCAAGGTTCTTTAAA | GCGCCGTCTCACTCAATGGGTGCTCTAGCACTAG |
| CaMV *35S minimal promoter #12 (minPro12)* | A3-B1 | *pUPD2_minPro12* | https://benchling.com/s/seq-17SsWNaS2VpKoAAfKdHW?m=slm-AfGcnYyiOjisWz | GGGAATCTCTGGCTCAGTCG | CGTGGAAGCGAATCTAGACC |
| CaMV *35S minimal promoter #13 (minPro13)* | A3-B1 | *pUPD2_minPro13* | https://benchling.com/s/seq-m5yfC9t3tzVPr7gll0Zs?m=slm-lPGsQiUTaVswM5BaX | GGGAATCTCTGGCTCAGTCG | CGTGGAAGCGAATCTAGACC |
| CaMV *35S minimal promoter #14 (minPro14)* | A3-B1 | *pUPD2_minPro14* | https://benchling.com/s/seq-6DoXPFWWSpw0GYgqGdd5?m=slm-BVXXSdccX3Y | GCGCCGTCTCACTCGTCCCCACTAAGGCCTAGG | GGGCCTAAGTCTACCCCATTGAGTGAGACGGCGC |
| CaMV *35S minimal promoter #15 (minPro15)* | A3-B1 | *pUPD2_minPro15* | https://benchling.com/s/seq-Yy95gZvfuaDMLxuKedOQ?m=slm-RLgScBwPJmFR2 | GCGCCGTCTCACTCGTCCCATGTGCAATAGTGTC | GCGCCGTCTCACTCAATGGACCCTTGGAGGCAC |
| CaMV *35S minimal promoter #16 (minPro16)* | A3-B1 | *pUPD2_minPro16* | https://benchling.com/s/seq-vaHHCGEBI1ILpkS65cxw?m=slm-zXzreCCFWNGy9a | GCGCCGTCTCACTCGTCCCTTACATCAAGCAAGT | GCGCCGTCTCACTCAATGGGTGGTATGGGGCTC |
| CaMV *35S minimal promoter #17 (minPro17)* | A3-B1 | *pUPD2_minPro17* | https://benchling.com/s/seq-ECN4YGBizNQQjUMgzl1p?m=slm-xSG8dcSx7LRwlH | GCGCCGTCTCACTCGTCCCTTGCCCACCCCATTA | GCGCCGTCTCACTCAATGGAGTGTGTCCCTCTC |
| CaMV *35S minimal promoter #18 (minPro18)* | A3-B1 | *pUPD2_minPro18* | https://benchling.com/s/seq-BbTdKgtf5n2S5YUnpqll?m=slm-sKcnSykXa52ST9JX | GCGCCGTCTCACTCGTCCCGGATAGCTCAAGCTA | GCGCCGTCTCACTCAATGGGCCCCTCCACAATAG |
| CaMV *35S minimal promoter #19 (minPro19)* | A3-B1 | *pUPD2_minPro19* | https://benchling.com/s/seq-YUFF5YHMSZsYoqDjgmZZ?m=slm-VcDAq2dQNdj7C | GCGCCGTCTCACTCGTCCCAGTTGGGACCATCAA | GCGCCGTCTCACTCAATGGTCCAATGGGCAAGTG |
| CaMV *35S minimal promoter #20 (minPro20)* | A3-B1 | *pUPD2_minPro20* | https://benchling.com/s/seq-dV5xc51RdD1cwlafLTuE?m=slm-7CKrBRszA7x6bEr | GCGCCGTCTCACTCGTCCCGGGACCTAGGTCCAA | GCGCCGTCTCACTCAATGGGGATTAGCAACCCTC |
| FMV *Sgt VI minimal promoter (FMVminPro)* | A3-B1 | *pUPD2_minProFMV* | https://benchling.com/s/seq-1ZlORWL3pvDF5sV5HhUX?m=slm-kgDKK5fZggv6HU | GCGCCGTCTCGCTCGTCCCACGTATGATTTCAAAAAACGC | GCGCCGTCTCGCTCAATGGAGAATTTGTGTTGATGAAAAC |
| HRLV *Sgt VI minimal promoter (HRLVminPro)* | A3-B1 | *pUPD2_minProHRLV* | https://benchling.com/s/seq-eUliyXAQpuGPqSyyF8z1?m=slm-2GWCKKTX9kmA7 | GCGCCGTCTCGCTCGTCCCATGACCGATCAATCCCACTATC | GCGCCGTCTCGCTCAATGGTTCTCCAAATGAAGCAACTTC |

Newly designed DNA variants of the minimal promoter from CaMV 35S gene[51,52] were amplified from GBlock synthetic DNA composed of multiple (three to ten) DNA parts flanked by part-specific GB codes and stacked together as a cost-cutting measure. Construct-specific oligos that include a tail with the GB cloning overhang were designed to amplify minPro sequences prior to their *pUPD2* subcloning. Alternatively, sequences of oligos from our lab collection were used as sequence stuffers in the design of the multisequence GBlocks which enabled selective PCR amplification of the DNA fragments flanked by the *pUPD2* cloning overhang (when allowed by vendor's synthesis thresholds). Minimal promoters from FMV SxD strain Sgt VI (19S) [58-59] and Horseradish latent virus (HRLV) Sgt VI[60] deduced from previously described full promoters were also amplified from GBlock synthetic DNA. Sgt: subgenomic transcript;

**D**

| DNA part | GB category | Clone name | Link to the clone (Benchling supported) | Primer forward | Primer reverse |
|---|---|---|---|---|---|
| **SUBCELULAR LOCALIZATION SIGNALS, N-TERMINAL/C-TERMINAL** | | | | **Primer forward** | **Primer reverse** |
| *Mitrocondrial (MitCox4)* | 32 (N-terminal | *pUPD2_MTS* | https://benchling.com/s/seq-Jdh8SFF6z2KtcJRj5rWA?m=slm-ujqTzd0iRiXlfKZO07 | GTAATACGACTCACTATAGGGC | AATTAACCCTCACTAAAGGG |
| *Peroxisomal (KSRM targeting peptide)* | 35 (C-terminal | *pUPD2_PTS1* | https://benchling.com/s/seq-EWlnAOolRzPaO4NWB4US?m=slm-8SxYP4NArXVs | GCGCCGTCTCGCTCGTTCGAAGTCACGTATGTGA | GCGCCGTCTCGCTCAAAGCTCACATACGTGACTT |
| *Nuclear N7v1* | 35 (C-terminal | *pUPD2_N7v1* | https://benchling.com/s/seq-IbkL6WxCpwBgPnaPrlar?m=slm-jlAbLAjYAV9URuoB | GCGCCGTCTCGCTCGTTCGGCTGCAGCCGCTGAGTTTAA | GCGCCGTCTCGCTCAAAGCTCACTCTTCTTCCTGATCGG |
| *Nuclear N7v2* | 35 (C-terminal | *pUPD2_N7v2* | https://benchling.com/s/seq-5dqyA01vWisy4mc4DGaS?m=slm-qp5Cs9KZbvjrH0Y | GCGCCGTCTCGCTCGTTCGGCAGCCGCTGCAGAATTTAA | GCGCCGTCTCGCTCAAAGCTTATTCCTCTTCTTGGTCAGC |
| *Nuclear N7v3* | 35 (C-terminal | *pUPD2_N7v3* | https://benchling.com/s/seq-2ov3ugLhnKgsR6KqoYRQ?m=slm-80LjiMiARtT052D | GCGCCGTCTCGCTCGTTCGGCCGCTGCAGCAGAATTCAA | GCGCCGTCTCGCTCAAAGCTCATTCTTCCTCTTGATCTGC |

Subcellular localization signals, N-terminal/C-terminal. *Mitochondrial Targeting Signal (MTS)* derived from the *CYTOCHROME C-OXIDASE SUBUNIT 4*[70-71] was amplified from GBlock synthetic DNA. The *MitCox4* sequence was synthesized with the GB cloning overhangs and later amplified with oligos from our lab collection used as stuffers in the synthesis of the multisequence GBlocks. *Peroxisomal Targeting Signal 1 (PTS)* was previously described[74]. The original *Nuclear Localization Signal (NLS) N7* was previously described[89].

**E**

| DNA part | GB category | Clone name | Link to the clone (Benchling supported) | Primer forward | Primer reverse |
|---|---|---|---|---|---|
| **REPORTER PROTEINS, N-TERMINAL/C-TERMINAL** | | | | **Primer forward** | **Primer reverse** |
| *Red Fluorescent Protein, mCherry C-TERMINAL* | B3-B5 | *pUPD2_C-mCherry* | https://benchling.com/s/seq-GZEVPRGSvglvqdAvX2Yw?m=slm-H8ZWaplMHWpa | TCGAGGTCGACGGTATC | CGCTCTAGAACTAGTGGATC |
| *Red Fluorescent Protein, mCherry N-TERMINAL* | B2-B4 | *pUPD2_FIXED_N-mCherryv1* | https://benchling.com/s/seq-xJ1eEZnTpUhAODvsfpRY?m=slm-1PQYElaUhAAFol | GCGCCGTCTCGCTCGCCATGGTGAGTAAGGGTGAAGAA | GCGCCGTCTCGCTCACGAACCTTTATAGAGCTCATCCATACCTC |
| *Blue Fluorescent Protein, mTagBFP* C-TERMINAL* | B3-B5 | *pUPD2_N-mTagBFP** | https://benchling.com/s/seq-2LRTCGSOsbtVAV9hihJM?m=slm-dcffbatv3Nb9BEC | GCGCCGTCTCACTCGAATGTCAGAACTTATCAAG | GCGCCGTCTCACTCAAAGCTTAGTTAAGCTTGTG |
| *Blue Fluorescent Protein, mTagBFP2v1-linker_v1 N-TERMINAL* | B2-B4 | *pUPD2_Full_mTagBFP2v1_linker1_B* | https://benchling.com/s/seq-pTwTpbZwxnTY6rkhYahn?m=slm-pX1B6wrxz02kfgf9 | GCGCCGTCTCGCTCGCCATGGTGTCTAAAGGGGAAGAGCTC | GCGCCGTCTCGCTCACGAACCAGACCCTGATCCGGAACCATTG |
| *Blue Fluorescent Protein, mTagBFP2v1-linker_v1 C-TERMINAL* | B3-B5 | *pUPD2_Full_mTagBFP2v1_linker1_B* | https://benchling.com/s/seq-kfuRCND9OHqnxaQ4RhVH?m=slm-Tp43MEfHZCjOE | GCGCCGTCTCGCTCGAATGGTGTCTAAAGGGGAAGAGCTC | GCGCCGTCTCGCTCAAAGCTTAAGACCCTGATCCGGAACCATTG |
| *3x mTagBFP2 [FPv1_linker_v1+FPv2_linker_v2+FPv3_linker_v3]* | B2-B4 | *pUPD2_3xFull_mTagBFP2_linker_B2* | https://benchling.com/s/seq-35OPT6GKV2zOa6wizXPe?m=slm-OKwjk5EwSS8nx | GCGCCGTCTCGCTCGCCATGGTGTCTAAAGGGGAAGAGCTC | GCGCCGTCTCGCTCACGAACCACTACCAGACCCTGATCCATTC |
| *3x mTagBFP2 [FPv1_linker_v1+FPv2_linker_v2+FPv3_linker_v3]* | B3-B5 | *pUPD2_3xFull_mTagBFP2_linker_B3* | https://benchling.com/s/seq-9Y5jkOhKXoTyj2QLRXz3?m=slm-rg10rr3M3QvGEjifiy | GCGCCGTCTCGCTCGAATGGTGTCTAAAGGGGAAGAGCTC | GCGCCGTCTCGCTCAAAGCTCAACTACCAGACCCTGATCCATTC |
| *Yellow Fluorescent Protein, YPet N-TERMINAL* | B2-B4 | *pUPD2_FixedGrammar_N-YPetv1* | https://benchling.com/s/seq-qWZ4HyH412BC6n9taCsD?m=slm-wTyhfMrUdG4e0u | GCGCCGTCTCGCTCGCCATGGTATCAAAGGGAGAGGAACTT | GCGCCGTCTCGCTCACGAACCCTTATACAATTCATTCATTCCTTCTG |

Initially, mTagBFP* was used in ACE as blue fluorescent protein (hybrid version of mTagBFP that contains an mTagBFP2-like amino acid substitution in the mTagBFP chromophore moiety but does not include the mTagBFP2 N-terminus). Later, it was upgraded to full mTagBFP2[69] in ACE2. FP DNA sequences were amplified from GBlock synthetic DNA. The mCherry C-terminal CDS sequence was synthesized with the GB cloning overhangs and amplified using oligos from our lab collection used as stuffers in the synthesis of multisequence GBlocks.

**F**

| DNA part | GB category | Clone name | Link to the clone (Benchling supported) | Primer forward | Primer reverse |
|---|---|---|---|---|---|
| **TERMINATORS** | | | | **Primer forward** | **Primer reverse** |
| CaMV *35S terminator #0 (ter0)* | C1 | *pUPD2_ter0* | https://benchling.com/s/seq-4mHRgylA35B8VMj9x27P?m=slm-vpAgWwhw3rZMy4 | GCGCCGTCTCACTCGGCTTAAATCACCAGTCTCTCTCTACAAATC | GCGCCGTCTCACTCAAGCGCTGGATTTTGGTTTTAGGAATTAG |
| CaMV *35S terminator #1 (ter1)* | C1 | *pUPD2_ter1* | https://benchling.com/s/seq-5Rg4HEjCAGtG0Zc7P21z?m=slm-91P4vQBw4W9Kx | GTAATACGACTCACTATAGGGC | AATTAACCCTCACTAAAGGG |
| CaMV *35S terminator #2 (ter2)* | C1 | *pUPD2_ter2* | https://benchling.com/s/seq-NyFGnExVRlqYwbimTHxq?m=slm-eS1mTbMnoeZia8 | GTAAAACGACGGCCAGTG | GGAAACAGCTATGACCATG |
| CaMV *35S terminator #3 (ter3)* | C1 | *pUPD2_ter3* | https://benchling.com/s/seq-KbRS4DnnoyBGgOhQ3TE3?m=slm-TL8ZpoFfKx3Yt | GCGCCGTCTCACTCGGCTTGTTGGCTCAAATTATG | GCGCCGTCTCACTCAAGCGTTAATTATGGTGGGAG |
| CaMV *35S terminator #4 (ter4)* | C1 | *pUPD2_ter4* | https://benchling.com/s/seq-oVVMBdcl6xaYIbSnGhiK?m=slm-Uz94tKZNrOlqyV2t | GCGCCGTCTCACTCGGCTTATATAGATCCATCTATG | GCGCCGTCTCACTCAAGCGTTGTGAATAGTCTAT |
| CaMV *35S terminator #5 (ter5)* | C1 | *pUPD2_ter5* | https://benchling.com/s/seq-fnb4xgUe8b2jXsHDUHeu?m=slm-0nuoUb0JJcNmhG | GCGCCGTCTCACTCGGCTTCTCACACAATTAGC | GCGCCGTCTCACTCAAGCGTATTTTGTAACATGG |
| CaMV *35S terminator #6 (ter6)* | C1 | *pUPD2_ter6* | https://benchling.com/s/seq-OLU6z6PUMgEAoJq95uTu?m=slm-l98DMKCCvg6DN | GCGCCGTCTCACTCGGCTTATTAATATTGGCCCC | GCGCCGTCTCACTCAAGCGTAGTTAGACAATTAGG |
| CaMV *35S terminator #7 (ter7)* | C1 | *pUPD2_ter7* | https://benchling.com/s/seq-xkhYL3WSSKb4hCgBNiKd?m=slm-KxbwHwyglsROtN | GCGCCGTCTCACTCGGCTT TGTTTAGTTATGTGATAG | GCGCCGTCTCACTCAAGCGTCAAGTTATATATTATTG |
| CaMV *35S terminator #8 (ter8)* | C1 | *pUPD2_ter8* | https://benchling.com/s/seq-4u4K6YdGKJs7jcOVoIE4?m=slm-5xgoxFNkzJWUfxY | GCGCCGTCTCACTCGGCTTACTTGGTGAGAGGTC | GCGCCGTCTCACTCAAGCGTTTGGTTCTAAGTTGAG |
| CaMV *35S terminator #9 (ter9)* | C1 | *pUPD2_ter9* | https://benchling.com/s/seq-9ZdkuHrm706Qe9m29aBF?m=slm-yRFkCOEQzuKU1 | GCGCCGTCTCACTCGGCTTGTAACAAGAATTTATTG | GCGCCGTCTCACTCAAGCGAACTTATATTGTGAG |
| CaMV *35S terminator #10 (ter10)* | C1 | *pUPD2_ter10* | https://benchling.com/s/seq-EPfnAJQdu7EtYZ5Bvyos?m=slm-3atVUCQ9BWqJlA | GCGCCGTCTCACTCGGCTTCCTTTTCACATTTAG | CAACTATACACCACCTTCGCTTGAGTGAGACGGCGC |
| CaMV *35S terminator #11 (ter11)* | C1 | *pUPD2_ter11* | https://benchling.com/s/seq-2BoBSBMxCB6pBiiXTI50?m=slm-iNU9jmHpGkEACm | GGGAATCTCTGGCTCAGTCG | CGTGGAAGCGAATCTAGACC |
| CaMV *35S terminator #12 (ter12)* | C1 | *pUPD2_ter12* | https://benchling.com/s/seq-jKcAISaMSymnUUhz4QnQ?m=slm-kBP6mhIQBX0Nj | GGAGCCTTCTGGTTTGTATCTC | CCACTAGCATGTTCTGGCAC |
| CaMV *35S terminator #13 (ter13)* | C1 | *pUPD2_ter13* | https://benchling.com/s/seq-y2qr8TZDt6OA0jFtPW9z?m=slm-Twc0BlsLsyVZGTJc | GCGCCGTCTCACTCGGCTTACTCTCATTAGTAATC | GCGCCGTCTCACTCAAGCGTACTTATGTGGTGTATATG |
| CaMV *35S terminator #14 (ter14)* | C1 | *pUPD2_ter14* | https://benchling.com/s/seq-D3URv5dz1rVLgUegdKDR?m=slm-AdoxNCotlNWpaw | GGAGCCTTCTGGTTTGTATCTC | CCACTAGCATGTTCTGGCAC |
| CaMV *35S terminator #15 (ter15)* | C1 | *pUPD2_ter15* | https://benchling.com/s/seq-cYyZFVCJ2QACsliTxYEK?m=slm-QB6Kua2bRNSkai | GCGCCGTCTCACTCGGCTTGTAAGCCTAGTCTTTC | GCGCCGTCTCACTCAAGCGGCTTGATTATTGAGGG |
| CaMV *35S terminator #16 (ter16)* | C1 | *pUPD2_ter16* | https://benchling.com/s/seq-PI3AfOoF74xrezQITAkl?m=slm-kozFAV3bg5V95GST | TCCAATGGCTCAGAGCTTCG | CGCATCTCCAAGCTAATGTGG |
| CaMV *35S terminator #17 (ter17)* | C1 | *pUPD2_ter17* | https://benchling.com/s/seq-ewSzon40sU8HcZYtw5BM?m=slm-SJ80xuQqmK0b1 | GGGAATCTCTGGCTCAGTCG | CGTGGAAGCGAATCTAGACC |
| CaMV *35S terminator #18 (ter18)* | C1 | *pUPD2_ter18* | https://benchling.com/s/seq-3szinAYtAgXQYhIeBZea?m=slm-p8AF6HVAy2ojqGh | TCCAATGGCTCAGAGCTTCG | CGCATCTCCAAGCTAATGTGG |
| CaMV *35S terminator #19 (ter19)* | C1 | *pUPD2_ter19* | https://benchling.com/s/seq-kusUriCVPeiONuQ2uxu0?m=slm-vYGiZsyb1HH3igSy | GCGCCGTCTCACTCGGCTTTAACATCATGGTCCTATC | GCGCCGTCTCACTCAAGCGATTATTTGATTGATAGGC |
| CaMV *35S terminator #20 (ter20)* | C1 | *pUPD2_ter20* | https://benchling.com/s/seq-H6tTFhtu1i6nTTvoKN14?m=slm-6Dnf9S5alHrR3iH5P | GCGCCGTCTCACTCGGCTTGACCCTTGTTGGTGG | GCGCCGTCTCACTCAAGCGCTTAGTGGAATAATTG |

Newly designed DNA variants of the terminator from CaMV 35S gene [51,52] were amplified from GBlock synthetic DNA composed of multiple (three to ten) DNA parts flanked by part-specific GB codes and stacked together as a cost-cutting measure. Individual terminator amplification was carried out in a similar manner as described above for *minPro*.

**G**

| DNA part | GB category | Clone name | Link to the clone (Benchling supported) | Primer forward | Primer reverse |
|---|---|---|---|---|---|
| **NOS- AND 35S-FREE SELECTABLE MARKERS** | | | | **Primer forward** | **Primer reverse** |
| *CVMV Fltp:NPTII:HSP18.2ter* | pDGB3α1 | *pDGB3α1_CVMVp:NPTII:tHsp18.2* | https://benchling.com/s/seq-ptQNY1eL21vpc5ELEH4G?m=slm-QuhWPp0RZ7wX | *CVMV Fltp* and *FMV 19Sp* promoters were PCR-amplified and cloned into *pUPD2* using the primers shown in (B) | |
| *CVMV Fltp:NPTII:HSP18.2ter* | pDGB3α2 | *pDGB3α2_CVMVp:NPTII:tHsp18.2* | https://benchling.com/s/seq-YdpCkonJwT54VeW1UDM3?m=slm-RlOS7tLnGq4gu | | |
| *CVMV Fltp:Bar:HSP18.2ter* | pDGB3α1 | *pDGB3α1_CVMVp:Bar:tHsp18.2* | https://benchling.com/s/seq-VZ8nc5wxvzDPvIAEKIQz?m=slm-wpz5StcxFghqnIiZs | Redomestication of *NPTII* as B3-B5: | |
| *CVMV Fltp:Bar:HSP18.2ter* | pDGB3α2 | *pDGB3α2_CVMVp:Bar:tHsp18.2* | https://benchling.com/s/seq-udvRghiv3D78bk3y8Dsg?m=slm-eOIYfmfeZkgW3se | GCGCCGTCTCGCTCGAATGATTGAACAAGATGGATTGCAC | GCGCCGTCTCGCTCAAAGCTCAGAAGAACTCGTCAAGAAG |
| *FMV 19Sp:NPTII:HSP18.2ter* | pDGB3α1 | *pDGB3α1_FMVp:NPTII:tHsp18.2* | https://benchling.com/s/seq-DqeS51t2J4iZxNNCMlcy?m=slm-tBcnoVDVqvU4hTx | Redomestication and reorientation of *BAR* as B3-B5: | |

| | | | | | |
|---|---|---|---|---|---|
| *FMV 19Sp:NPTII:HSP18.2ter* | pDGB3α2 | *pDGB3α2_FMVp:NPTII:tHsp18.2* | https://benchling.com/s/seq-lgE5TiHmugmY9RK1GFIw?m=slm-uYRTh9PK6ng3c[illegible] | GCGCCGTCTCGCTCGAATGAGCCCAGAACGACGCCC | GCGCCGTCTCGCTCAAAGCTCAGATTTCGGTGACGGGCA |
| *FMV 19Sp:Bar:HSP18.2ter* | pDGB3α1 | *pDGB3α1_FMVp:Bar:tHsp18.2* | https://benchling.com/s/seq-3tdVucAN1hFXk7RIjA4y?m=slm-5qCQPXYKRpCj17[illegible] | *HSP18.2ter*: | |
| *FMVp:Bar:HSP18.2ter* | pDGB3α2 | *pDGB3α2_FMVp:Bar:tHsp18.2* | https://benchling.com/s/seq-UHY3SocH30mmJB9h6kz0?m=slm-7DOqASMjrD75P[illegible] | GCGCCGTCTCGCTCGGCTTATATGAAGATGAAGATGAAATATTTGG | GCGCCGTCTCGCTCAAGCGATGAAGGGAGGTTTATAGGTC |

CVMV Fltp: 2895 bp-promoter from the CVMV full-length transcript[63 65] was amplified from GBlock synthetic DNA. FMV 19Sp: 416 bp-promoter from the FMV M3 strain Sgt VI (19S)[63 64] was amplified from GBlock synthetic DNA. HSP18.2ter: terminator from Arabidopsis thaliana HEAT SHOCK PROTEIN18.2[116].

**H**

| **OTHER *pUPD2* CLONES PREVIOUSLY REPORTED** | | | | **Primer forward** | **Primer reverse** |
|---|---|---|---|---|---|
| *Ethylene Distal-Proximal Inducible Promoter, EBSn* | A1-A2 | *pUPD2_EBSn* | ref 52 | ref 52 | ref 52 |
| *Nuclear Signal, 3xSV40 N-terminal* | B2 | *pUPD2_3xNLS* | ref 51, 66 | ref 51, 66 | ref 51, 66 |
| *Yellow Fluorescent Protein, 3xYPet C-TERMINAL* | B3-B5 | *pUPD2_3xYPet* | ref 51 | ref 51 | ref 51 |
| *BASTA resistance, pUPD_NOSter:BAR:NOSp* | A1-C1 (TU) | *GB0023* | ref 51 | ref 51 | ref 51 |
| *KANAMYCIN resistance, pUPD_NOSp:NPTII:NOSter* | A1-C1 (TU) | *GB0184* | ref 51 | ref 51 | ref 51 |

A1-C1 (TU) refers to the full transcriptional unit (*promoter:CDS:terminator*) subcloned into an entry vector, *pUPD*, from the GoldenBraid collection (GB0023 or GB0184). They were all described in a previous work[52].

**I**

| **ADDITIONAL DUAL LUCIFERASE/RENILLA *pUPD2* CLONES** | | | | **Primer forward** | **Primer reverse** |
|---|---|---|---|---|---|
| *35S distal-proximal promoter* (*minPro0*-less) | A1-A2 | *pUPD2_35S_distal-proxProm* | https://benchling.com/s/seq-nrNQR677BPz8hDHm2Buw?m=slm-HLzzdjVMerSugcxAK0Eg | GCGCCGTCTCGCTCGGGAGGGAGACTAGAGCCAAGCTGA | GCGCCGTCTCGCTCAGGGAGAAGGATAGTGGGATTGTGC |
| *LUCIFERASE CDS* | B3-B5 | *GB0096* | https://goldenbraidpro.com | Ready-to-use piece from the original GB collection (GB0096) | |
| *LUCIFERASE CDS* | B2-B5 | *pUPD2_LUCIFERASE_b2b5* | https://benchling.com/s/seq-xA2k9YLk7imM2eJoCxe2?m=slm-8wtmpLFCJ5cetfSWWuv3 | GCGCCGTCTCGCTCGCCATGGAAGACGCCAAAAACATAAAG | GCGCCGTCTCGCTCAAAGCTTACACGGCGATCTTTCCGCCC |
| *LUCIFERASE CDS - 35S terminator #0* | B2-C1 | *pUPD2_LUCIFERASE_b2b5:ter0* | https://benchling.com/s/seq-ag3Q932nn5T0WO29x3I4?m=slm-S6gBH59wxtpj1u4G6Stw | GCGCCGTCTCGCTCGCCATGGAAGACGCCAAAAACATAAAG | GCGCCGTCTCACTCAAGCGCTGGATTTTGGTTTTAGGAATTAG |
| *RENILLA CDS* | B3-B5 | *pUPD2_RENILLA* | https://benchling.com/s/seq-qvMaUOfjCZ5zDCDk2XM8?m=slm-k1sCrHptclAfRVp88Ts9 | GCGCCGTCTCGCTCGAATGACTTCGAAAGTTTATGATCCA | GCGAGTTCTCAAAAATGAACAATAAGCTTTGAGCGAGACGGCGC |

The GoldenBraid website is given as reference for the *pUPD* entry clone GB0096 from the GoldenBraid collection. The *minPro-less* 35S proximodistal promoter (A1-A2) sequence was amplified from the *pUPD* entry clone GB0030 (*pUPD_fullPro35S*) from the GoldenBraid collection. The *RENILLA* CDS sequence from the clone GB0109 (*pEGB_35S:REN:tNOS*) was redomesticated into *pUPD2* entry clone to modify the flanking GB codes and make them compatible with the other DNA elements to assemble.

**Support information:** Transcriptional Reporter GB parts (in blue), their GB codes for directional assembly (in grey) and the corresponding units in the final reporter (in **bolded black**)

*For **N-Tag** signals:*

| GB code | A1-A2 | A2-A3 | A3-B1 | B1-B2 | B2-B3 | B3-B4 | B4-B5 | B5-C1 |
|---|---|---|---|---|---|---|---|---|
| DNA part | | | | | | | | |
| GB Category | Distal (A1) | Proximal (A2) | Core+UTR | N-TAG (B2) | CDS (B3-B5) | | | Term (C1) |
| **Description** | **Hormone-inducible** | | **Core** | **Subcellular** | **Coding sequence for the reporter** | | | **Terminator** |

*For **C-Tag** signals:*

| GB code | A1-A2 | A2-A3 | A3-B1 | B1-B2 | B2-B3 | B3-B4 | B4-B5 | B5-C1 |
|---|---|---|---|---|---|---|---|---|
| DNA part | | | | | | | | |
| GB Category | Distal (A1) | Proximal (A2) | Core+UTR | CDS (B2-B4) | | | C-TAG (B5) | Term (C1) |
| **Description** | **Hormone-inducible promoter** | | **Core promoter** | **Coding sequence for the reporter protein (fluorescence, histochemical)** | | | **Subcellular localization signal** | **Terminator** |

*For **minPro#:LUC** transcriptional unit:*

| GB code | A1-A2 | A2-A3 | A3-B1 | B1-B2 | B2-B3 | B3-B4 | B4-B5 | B5-C1 |
|---|---|---|---|---|---|---|---|---|
| DNA part | | | | | | | | |
| GB Category | Distal (A1) | Proximal (A2) | Core+UTR | CDS (B2-B5) | | | | Term (C1) |
| **Description** | **Proximo-distal region from the full *35S* promoter** | | **Core promoter to test** | ***LUCIFERASE* CDS (B2-B5) compatible with the core promoters and fused to the original terminator from the *35S* gene (*ter0*)** | | | | |

*For **LUC:term#** transcriptional unit:*

| GB code | A1-A2 | A2-A3 | A3-B1 | B1-B2 | B2-B3 | B3-B4 | B4-B5 | B5-C1 |
|---|---|---|---|---|---|---|---|---|
| DNA part | | | | | | | | |
| GB Category | Distal (A1) | Proximal (A2) | Core+UTR | N-TAG (B2) | CDS (B2-B5) | | | Term (C1) |
| **Description** | **Full *35S* promoter (GB0030)** | | | | ***LUCIFERASE* CDS (GB0096)** | | | **Terminator to test** |

*For **Constitutive viral promoters (same type of display as for LUC:term transcriptional units):***

| GB code | A1-A2 | A2-A3 | A3-B1 | B1-B2 | B2-B3 | B3-B4 | B4-B5 | B5-C1 |
|---|---|---|---|---|---|---|---|---|
| DNA part | | | | | | | | |
| GB Category | Distal (A1) | Proximal (A2) | Core+UTR | N-TAG (B2) | CDS (B2-B5) | | | Term (C1) |
| **Description** | **Constitutive viral promoters** | | | | **Reporter gene** | | | **Terminator** |
